# Anti-inflammatory dopamine- and serotonin-based endocannabinoid epoxides reciprocally regulate cannabinoid receptors and the TRPV1 channel

**DOI:** 10.1101/2020.05.05.079624

**Authors:** William R. Arnold, Lauren N. Carnevale, Zili Xie, Javier L. Baylon, Emad Tajkhorshid, Hongzhen Hu, Aditi Das

## Abstract

The endocannabinoid (eCB) system is a promising target to mitigate pain as the eCBs are endogenous ligands of the pain-mediating receptors—cannabinoid receptors 1 and 2 (CB1 and CB2) and TRPV1. Here we report on a novel class of lipids formed by the epoxidation of N-arachidonoyl-dopamine (NADA) and N-arachidonoyl serotonin (NA5HT) by cytochrome P450 (CYP) epoxygenases. These epoxides (epoNADA and epoNA5HT) are dual-functional rheostat modulators (varying strength of agonism or antagonism) of the eCB-TRPV1 axis. In fact, epoNADA is a 6-fold stronger agonist of TRPV1 than NADA while epoNA5HT is a 30-fold stronger antagonist of TRPV1 than NA5HT and displays a significantly stronger inhibition on TRPV1-mediated responses in primary afferent neurons. Moreover, epoNA5HT is a full CB1 agonist. The epoxides reduce the pro-inflammatory biomarkers IL-6, IL-1β, TNF-α and nitrous oxide (NO) and raise anti-inflammatory IL-10 in activated microglial cells. The epoxides are spontaneously generated by activated microglia cells and their formation is potentiated in the presence of another eCB, anandamide (AEA). We provide evidence for the direct biochemical mechanism of this potentiation using human CYP2J2, a CYP epoxygenase in the human brain, using detailed kinetics studies and molecular dynamics simulations. Taken all together, inflammation leads to an increase in the metabolism of NADA, NA5HT and other eCBs by CYP epoxygenases to form the corresponding epoxides. The epoxide metabolites are bioactive lipids that are more potent, multi-faceted endogenous molecules, capable of influencing the activity of CB1, CB2 and TRPV1 receptors. The identification of these molecules will serve as templates for the synthesis of new multi-target therapeutics for the treatment of inflammatory pain.

## INTRODUCTION

Opioids are highly addictive pain medications that are susceptible for abuse. The age-adjusted death rate by opioid overdose was determined to be nearly 20 per 100,000 people in the United States in 2016, according to a report by the Centers for Disease Control and Prevention^1^. Hence, there is a need for therapeutic alternatives to opioids that combat inflammation and the associated pain.

Pain is regulated primarily by sensory afferent neurons and immune cells. Both of these cell types are rich sources of lipid mediators. Lipid mediators are generated via the enzymatic oxidation of dietary omega-3 and omega-6 polyunsaturated fatty acids (PUFAs). The pro-inflammatory lipid mediators contribute to pain sensitivity by activating the GPCRs in the sensory neurons to increase membrane excitability and pain response^2^. For instance, cyclooxygenases generate pro-inflammatory lipids such as prostaglandin E_2_ (PGE_2_) at the sites of inflammation. Non-steroidal anti-inflammatory drugs (NSAIDs) are used to inhibit cyclooxygenases, leading to a decrease in the synthesis of pro-inflammatory lipid metabolites, thereby decreasing inflammatory pain^3^. On the other hand, anti-inflammatory and pro-resolving lipid mediators suppress and resolve the inflammatory process, and thus attenuate inflammatory pain^4,5^. Hence, lipid mediators can fine-tune the pain response and have been at the center for the development of alternative non-opioid pain therapeutics^6,7^.

Additionally, cannabis has been used for centuries to reduce nociceptive pain either alone or in combination with opioids^8^. The primary components of cannabis interact in the body with cannabinoid receptors 1 and 2 (CB1 and CB2) and other GPCRs^9-11^. An endogenous class of bioactive lipids, known as endocannabinoids (eCBs), activates cannabinoid receptors and suppresses inflammation and pain sensitization^12,13^. The eCBs are derivatives of dietary omega-3 and omega-6 PUFAs and are generated by damaged neurons and inflamed tissues. CB1 receptors are highly expressed in the central nervous system, mostly in the presynaptic region, and there is substantial CB1 expression in the nociceptive sensory neurons. It has been shown that under different pain conditions there is a concomitant increase in CB1 expression^14,15^. Hence, there is sufficient evidence that CB1 mediates the psychotropic effects of cannabinoids such as modulating nociceptive pain, as well as modulating inflammation^9,16-21^. CB2 is mostly expressed in immune cells and mediates the anti-inflammatory effects of cannabinoids, which indirectly contributes to the anti-nociception of acute inflammatory pain^19,21-23^. CB2 receptor activation exerts profound anti-nociceptive effects in animal models of acute, inflammatory, and neuropathic pain^24^.

In addition to CB1 and CB2, a subclass of the eCBs act through transient receptor potential vanilloid 1 (TRPV1)^25-27^. TRPV1 is a non-selective cation channel that is activated by noxious temperatures, pH, and chemical stimuli such as inflammatory agents^28-31^. TRPV1 often spearheads nociceptive pain signaling, and thus antagonizing TRPV1 can reduce pain. Paradoxically, the activation of TRPV1 by small molecules such as capsaicin (CAP), the spicy component of chili peppers, can also alleviate pain by desensitizing TRPV1 signaling and creating a numbing effect^32,33^. In addition to mediating the pain associated with these stimuli, TRPV1 exhibits pro- and anti-inflammatory effects^34-37^.

Recently, it has been postulated that there is a crosstalk between CB1 and TRPV1 receptors, which are co-localized in dorsal root ganglion and in neuron-enriched mesencephalic cultures, hippocampus, and cerebellum^38^. Therefore, the eCB system and TRPV1 axis provides a promising target to develop pain and inflammation therapeutics. For example, CMX-020 (Patent US8658632B2) is a novel drug based on the structure of eCBs and is in development to alleviate pain by binding to both cannabinoid receptors and TRPV1. Endogenous molecules and their synthetic derivatives may provide insight into effective therapeutic strategies with which to target the eCB-TRPV1 axis, as these molecules effectively act upon both receptors unlike cannabinoids or vanilloids separately.

The best-studied eCB is anandamide (N-arachidonoyl-ethanolamine: AEA), which is derived from the omega-6 PUFA arachidonic acid (AA)^39-42^. Besides AEA, other eCBs such as N-arachidonoyl-dopamine (NADA) and N-arachidonoyl-serotonin (NA5HT) are also derivatives of AA (Figure 1). These dopamine (DA)^43-45^ and serotonin (5HT)^46-48^ derivatives were identified *in vivo* in brain and intestinal tissues. It was shown that NADA binds with a higher affinity to CB1 than to CB2^43^; however, conclusive evidence for NA5HT binding to CB1/2 has yet to emerge.

**Figure 1.**
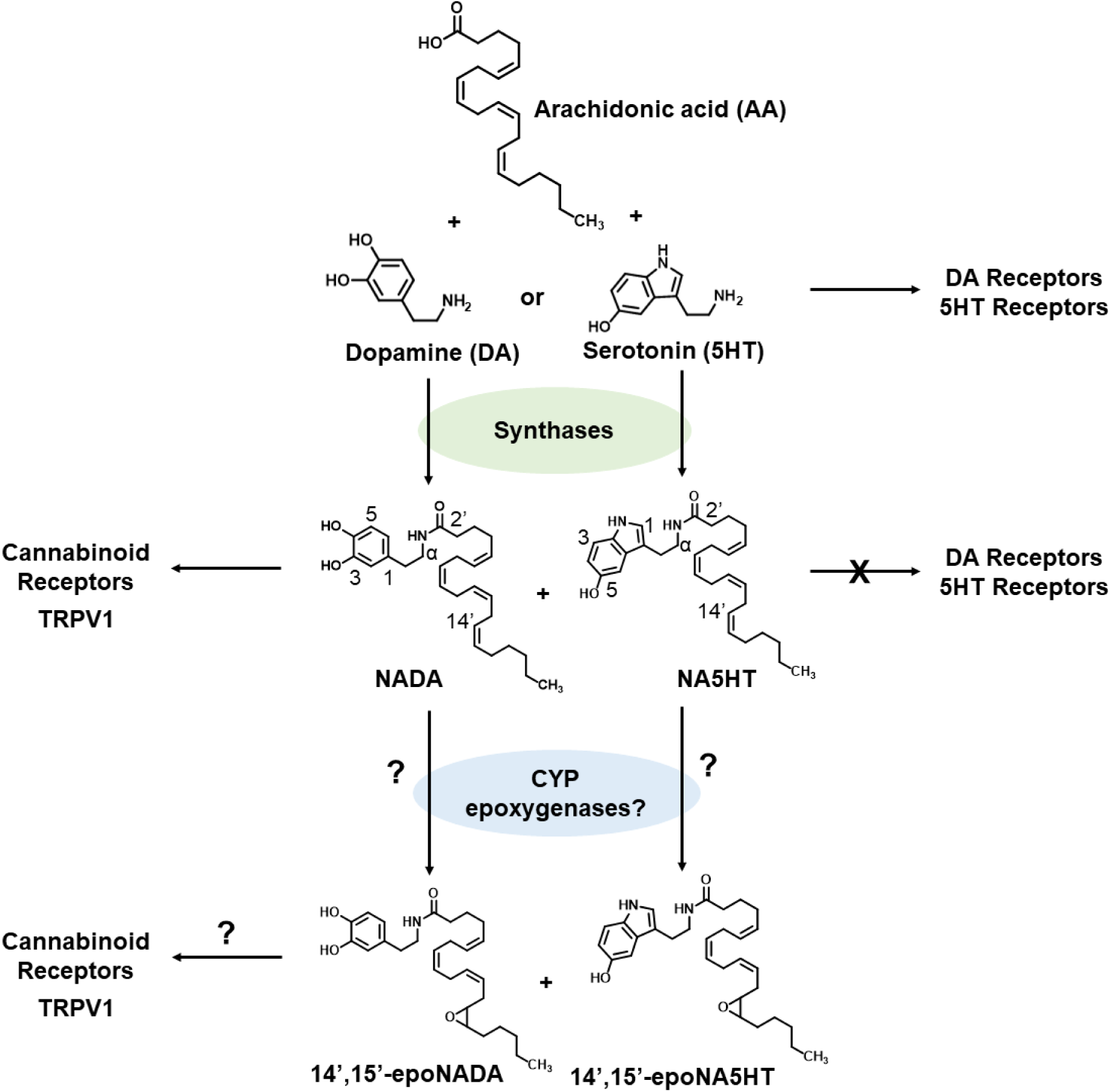
The endovanilloid (eVD) pathway and structures of 14’,15’-epoNADA and 14’,15’-epoNA5HT.

Additionally, NADA was shown to be an agonist of TRPV1^49,50^ and NA5HT is an antagonist of TRPV1, which results in analgesia^51^. Hence, NADA and NA5HT are classified as endovanilloids (eVDs) for their actions at the TRPV1 “vanilloid” receptor. Interestingly, NADA has low-affinity binding to DA receptors^43^, and the stimulation of degranulation in mast cells by NA5HT suggests it does not activate the 5HT receptors^52^. Therefore, NADA and NA5HT do not display typical DA or 5HT responses and are instead regulators of TRPV1.

To add to the complexity of lipid metabolism, cytochromes P450 (CYPs) are known to epoxidize lipids into anti-inflammatory and anti-pain mediators that are more effective than the parent molecules. For example, CYPs convert AA and omega-3 PUFAs into epoxy-PUFAs that have been shown to decrease pain^53-55^. Recently, the CYP-mediated metabolism of eCBs was shown to produce epoxy-eCBs that exhibit CB2 receptor selectivity and are anti-inflammatory and anti-tumorigenic^56-58^. For instance, CYPs epoxidize AEA into epoxyeicosatrienoic acid ethanolamides (EET-EAs) that bind to CB2 receptors^56,59,60^. Similarly, omega-3 PUFA-derived eCBs are also converted by CYPs into epoxide metabolites that are anti-inflammatory and anti-tumorigenic^57^.

We therefore asked whether epoxide metabolites of eVDs can be more effective than parent eVDs at targeting cannabinoid receptors and TRPV1, and whether they are anti-inflammatory. Herein, we report a novel class of dual-functional epoxides of NADA and NA5HT (epoNADA and epoNA5HT, respectively) that reciprocally regulate both cannabinoid receptors (CB1 and CB2) and TRPV1 (Figure 1). We synthesized NADA, NA5HT, 14’,15’-epoNADA, and 14’,15’-epoNA5HT. Using targeted lipidomics, we were able to identify the eVDs and epoxy-eVDs in porcine brain tissue, but because of the low amounts of these and the variability of the results we cannot definitively conclude their presence and levels *in vivo*. We then explored if NADA and NA5HT are potentially epoxidized under inflammatory conditions. We show that epoNADA and epoNA5HT are formed under inflammatory conditions by CYP epoxygenases and show that AEA potentiates the formation of 14’,15’-epoNADA in microglial cells, which might occur through multiple mechanisms. We show that one mechanism of this potentiation may be through multiple-ligand binding to CYP-Nanodiscs (nanoscale lipid bilayers) using *in vitro* kinetics methods and molecular dynamics (MD) simulations. We further demonstrate that epoNA5HT is a potent TRPV1 antagonist suppressing intracellular Ca^2+^ response and membrane currents provoked by the TRPV1 ligand CAP in the primary afferent neurons. Together, we show that epoNADA and epoNA5HT act as dual CB1/2 and TRPV1 ligands and exhibit anti-inflammatory activity. These molecules may be interesting candidates on which to develop drugs.

## EXPERIMENTAL PROCEDURES

*Materials and methods concerning RT-qPCR, IL-6 ELISA kit based detection, Griess Assay to detect nitric oxide, cell viability assay using MTT, cell proliferation assay using BrdU, protein expression and purification methods, nanodisc assembly*, in vitro *lipid metabolism, and full-scan mass spectrometry analysis of the metabolites can be found in the supplementary information*.

### Synthesis of NADA [(5Z,8Z,11Z,14Z)-N-(3,4-dihydroxyphenethyl)icosa-5,8,11,14-tetraenamide] and NA5HT [(5Z,8Z,11Z,14Z)-N-(2-(5-hydroxy-1H-indol-3-yl)ethyl)icosa-5,8,11,14-tetraenamide]

Dopamine hydrochloride (24.9 mg, 0.131 mmol, 2.0 equiv.) or serotonin hydrochloride (27.9 mg, 0.131 mmol, 2.0 equiv.) was added to N,N-diisopropylethylamine (DIPEA, 27.4 μL, 0.145 mmol, 2.2 equiv.) in an anhydrous solution of DMF/CH_2_Cl_2_ (1/1 v/v, 25 mL) under an argon atmosphere. This mixture was cooled in an ice-water bath and then AA (21.7 μL, 0.066 mmol, 1.0 equiv.), 1-ethyl-3-(3-dimethylaminopropyl)-carbodiimide (EDC, 63 mg, 0.33 mmol, 5.0 equiv.), and 4-dimethylaminopyridine (DMAP, 2 mg, 0.0099 mmol, 0.15 equiv.) were added. After 1 hour, the reaction was warmed to room temperature and allowed to incubate at the same temperature for 8 hours. Afterwards, toluene was added, and the reaction was concentrated under reduced pressure. The product was extracted 3× from water with CH_2_Cl_2_ and then washed 1× with brine. The organic layer was dried over sodium sulfate and concentrated under reduced pressure to yield a yellow-brown oil. NADA and NA5HT were purified using HPLC Method 2 below. Products were dried under reduced pressure, stored in ethanol, and quantified via HPLC as stated below. Yields were 13.0 mg NADA (45%) and 12.8 mg NA5HT (45%). **[NADA]** ^1^H-NMR (400 MHz, CDCl_3_) δ 6.96 (s, 1H); 6.80 (d, *J* = 8.0 Hz, 1H); 6.74 (d, *J* = 2.0 Hz, 1H); 6.58 (dd, *J* = 8.1, 2.0 Hz, 1H); 5.61 (s, 1H); 5.54 (d, J = 6.5 Hz, 1H); 5.44 – 5.27 (m, 8H); 4.12 (q, J = 7.1 Hz, 1H) (ethyl acetate); 3.48 (q, *J* = 6.7 Hz, 2H); 2.88 – 2.75 (m, 6H) (part ethyl acetate); 2.70 (t, *J* = 7.1 Hz, 2H); 2.19 – 2.12 (m, 2H); 2.12 – 1.99 (m, 6H); 1.68 (p, *J* = 7.4 Hz, 2H); 1.41 – 1.19 (m, 8H) (part ethyl acetate); 0.88 (t, *J* = 6.7 Hz, 3H). HRMS (m/z): [M + H^+^] cal’d 440.3165, observed 440.3161 (−0.9 ppm); C_28_H_42_NO_3_. **[NA5HT]**. ^1^H-NMR (500 MHz, CDCl_3_) δ 7.90 (s, 1H); 7.04 – 6.98 (m, 2H); 6.82 –6.76 (m, 1H); 5.53 (s, 1H); 5.44 – 5.29 (m, 8H); 3.57 (q, J = 6.6 Hz, 2H); 2.90 (t, J = 6.8 Hz, 2H); 2.80 (dt, J = 18.2, 5.9 Hz, 6H); 2.08 (ddt, J = 31.1, 14.6, 7.4 Hz, 6H); 1.73 – 1.63 (m, 2H); 1.40 – 1.16 (m, 7H) (part ethanol); 0.88 (t, J = 6.7 Hz, 3H). HRMS (m/z): [M + H^+^] cal’d 463.3325, observed 463.3322 (−0.6 ppm); C_30_H_43_N_2_O_2._

### Synthesis of 14’,15’-epoNADA [(5Z,8Z,11Z)-N-(3,4-dihydroxyphenethyl)-13-(3-pentyloxiran-2-yl)trideca-5,8,11-trienamide] and 14’,15’-epoNA5HT [(5Z,8Z,11Z)-N-(2-(5-hydroxy-1H-indol-3-yl)ethyl)-13-(3-pentyloxiran-2-yl)trideca-5,8,11-trienamide]

Synthesis of EETs was performed using *m*CPBA as previously described, which yields 4-8 mg of 14,15-EET (7.6-15% yield)^57^. 14,15-epoNADA and 14,15-epoNA5HT were synthesized by coupling DA and 5HT to 14,15-EET using the above coupling method. Yield from 14,15-EET: 3.6 mg (65%) 14’,15’-epoNADA, 4.7 mg (40%) 14’,15’-epoNA5HT. **[14’,15’-epoNADA]** ^1^H-NMR (400 MHz, CDCl_3_) δ 6.80 (d, *J* = 8.1 Hz, 1H); 6.72 (s, 1H); 6.58 (d, *J* = 7.9 Hz, 1H); 5.81 (s, 1H); 5.65 – 5.25 (m, 6H); 3.47 (q, *J* = 6.5 Hz, 2H); 3.01 (d, *J* = 5.4 Hz, 1H); 2.80 (d, *J* = 20.6 Hz, 4H); 2.69 (t, *J* = 6.8 Hz, 2H); 2.45 (m, *J* = 7.3 Hz, 1H); 2.24 (m, *J* = 6.8 Hz 1H); 2.17 – 1.92 (m, 4H); 1.79 – 1.62 (m, 2H); 1.34 (q, *J* = 5.1, 3.7 Hz, 4H); 0.98 – 0.77 (m, 3H). HRMS (m/z): [M + H^+^] cal’d 456.3114, observed 456.3104 (−2.2 ppm); C_28_H_42_NO_4._ **[14’,15’-epoNA5HT]** ^1^H-NMR (500 MHz, CDCl_3_) δ 7.97 (s, 1H); 7.13 – 6.89 (m, 2H); 6.79 (dd, J = 8.7, 2.5 Hz, 1H); 5.60 (s, 1H); 5.55 – 5.29 (m, 7H); 5.23 (s, 1H); 3.56 (p, J = 8.1, 7.3 Hz, 2H); 2.96 (d, J = 7.2 Hz, 2H); 2.89 (q, J = 7.0 Hz, 2H); 2.79 (dt, J = 23.0, 6.3 Hz, 4H); 2.42 (d, J = 6.7 Hz, 1H); 2.21 (dt, J = 13.8, 6.8 Hz, 1H); 2.09 (dq, J = 20.9, 7.4 Hz, 4H); 1.68 (p, J = 7.6 Hz, 2H); 1.47 – 1.19 (m, 6H), 0.89 (d, J = 7.1 Hz, 3H). HRMS (m/z): [M + H^+^] cal’d 479.3274, observed 479.3269 (−1.0 ppm); C_30_H_43_N_2_O_3._

### Extraction of NADA and NA5HT from porcine brain regions

Porcine brains were obtained from the Meat Science Laboratory at the University of Illinois at Urbana-Champaign. The brains were dissected into regions (cerebellum; “central core” comprising the hippocampus, hypothalamus, and thalamus; and the cerebrum containing the cerebral cortex) and diced immediately after removal from the pig. The tissue was then flash frozen and stored at −80° C until used. For extraction, the tissue was homogenized in 5 volumes of ice-cold methanol (MeOH) containing 0.03 mM of the soluble epoxide hydrolase inhibitor 4-[[trans-4-[[(tricyclo[3.3.1.13,7]dec-1-ylamino)carbonyl]amino]cyclohexyl]oxy]-benzoic acid (t-AUCB) and 1 mM of FAAH inhibitor phenylmethylsulfonyl fluoride (PMSF). Debris was filtered using a filter column. The extract was then dried under reduced pressure and resuspended in 50% MeOH. The extracts were then loaded onto 1-gram Bond Elut C-18 cartridges (Varian, Harbor City, CA) (one column per gram of tissue), preconditioned with 25% MeOH. Cartridges were washed with 4 mL of 10% MeOH and eluted with 4 mL of 100% acetonitrile (MeCN). The eluates were dried under reduced pressure in amber vials and stored under Ar_(*g*)_ at −80° C until resuspended in 150 μL of 180-proof ethanol for LC-MS/MS analysis (the day after). Samples of tissue were spiked with 1 µg of NADA or NA5HT standards to test the recovery of this method, and we were able to recover 40% of the material.

### HPLC analysis of NADA, NA5HT, and metabolites

Compounds were analyzed via high-performance liquid chromatography (HPLC) consisting of an Alliance 2695 analytical separation module (Waters, Milford, MA) and a Waters 996 photodiode diode array detector (Waters). Epoxy-eVDs for synthesis purification were separated in reverse phase using a SunFire™ Prep C_18_ OBD™ 5 μm 19 x 50 mm column (Waters) and a 3.0 mL/min flow rate, and for quantification using a Phenomenex Prodigy® 5μm ODS-2, 150 x 4.60 mm column (Phenomenex, PN 00F-3300-E0, Torrance, CA) with a 1 mL/min flow rate. Mobile Phase A consisted of 95:5% H_2_O (0.1% acetic acid):MeCN and Mobile Phase B consisted of 5:95% H_2_O (0.1% acetic acid):MeCN. A full-scan method (Method 1) was developed to investigate all potential products from *in vitro* enzyme reactions as follows: 0-1 min, 100% A; 1-60 min, linear gradient of 100% A to 100% B; 60-65 min, 100% B. NADA and NA5HT elution times were confirmed using authentic standards (59.5 min and 60 min, respectively). A shorter method (Method 2) was developed to analyze the hydrophobic products and for synthesis purification and quantification. 0-30 min: 100% A to 100% B; 30-40 min: 100% B. All wavelengths from 190-600 nm were monitored. 14’,15’-epoNADA and 14’,15’-epoNA5HT were quantified at 281 nm and 277 nm wavelengths, respectively, using a NADA and NA5HT standard curve, respectively.

### LC-MS/MS quantification of NADA, NA5HT, and AEA from tissue

Samples were analyzed with the 5500 QTRAP LC/MS/MS system (Sciex, Foster City, CA) in Metabolomics Lab of Roy J. Carver Biotechnology Center, University of Illinois at Urbana-Champaign. Software Analyst 1.6.2 was used for data acquisition and analysis. The 1200 series HPLC system (Agilent Technologies, Santa Clara, CA) includes a degasser, an autosampler, and a binary pump. The LC separation was performed on an Agilent SB-Aq column (4.6 x 50mm, 5μm) with mobile phase A (0.1% formic acid in water) and mobile phase B (0.1% formic acid in acetontrile). The flow rate was 0.3 mL/min. The linear gradient was as follows: 0-1min, 90%A; 8-13min, 0%A; 13.5-18min, 90%A. The autosampler was set at 10°C. The injection volume was 10 μL. Mass spectra were acquired under positive electrospray ionization (ESI) with the ion spray voltage of 5500 V. The source temperature was 450 °C. The curtain gas, ion source gas 1, and ion source gas 2 were 32 psi, 50 psi, and 65 psi, respectively. Multiple reaction monitoring (MRM) was used for quantitation: AEA m/z 348.3 → m/z 203.2; NADA m/z 440.2 → m/z 287.1; 14’,15’-epoNADA m/z 456.3 → m/z 137.1; 14’,15’-epoNA5HT m/z 479.3 → m/z 160.1; NA5HT m/z 463.3 → m/z 287.2. Internal standard AEA-d4 was monitored at m/z 352.3 → m/z 287.2.

### Cell culture

HEK cells stably transfected with human TRPV1 (HEK-hTRPV1) were a gift from Prof. Bradshaw (University of Indiana, Bloomington), which were originally constructed by Merck Research. Cells were grown in Eagle Minimum Essential Media supplemented with L-glutamine (EMEM) (ATCC) and 10% fetal bovine serum (FBS) and were incubated at 37°C with 5% CO_2_. Cells were sub-cultured at 80-90% confluency by trypsinization in a 1:6-1:10 ratio. HTLA cells for PRESTO-TANGO and BV2 microglia were grown as previously described^57^.

### Isolation and short-term culture of mouse DRG neurons

Mice were killed by cervical dislocation following CO_2_ asphyxia and spinal columns were removed and placed in ice-cold HBSS. Laminectomies were performed and bilateral DRGs were dissected out. After removal of connective tissues, DRGs were digested in 1 mL of Ca^2+^/Mg^2+^-free HBSS containing 20 U of papain (Worthington, Lakewood, NJ), 0.35 mg of L-cysteine and 1 μL of saturated NaHCO_3_ and incubated at 37°C for 10 minutes. The DRG suspension was centrifuged, the supernatant was removed, and 1 mL of Ca^2+^/Mg^2+^-free HBSS containing 4 mg of collagenase type II and 1.25 mg of Dispase type II (Worthington) was added and incubated at 37°C for 15 minutes. After digestion, neurons were pelleted; suspended in neurobasal medium containing 1% L-glutamine, 2% B-27 supplement, 100 U · mL^−1^ penicillin plus 100 μg · mL^−1^ streptomycin, and 50 ng · mL^−1^ nerve growth factor. The cells were plated on a 12-mm coverslip coated with poly-l-lysine (10 μg · mL^−1^) and cultured under a humidified atmosphere of 5% CO_2_/95% air at 37°C for 24 hours.

### Metabolism of NADA and NA5HT by BV2 microglia with and without lipopolysaccharide (LPS) stimulation

BV2 microglia were plated on 6-well plates at 5 x 10^5^ cells per well and grown to 80-90% confluency. Cell growth media was exchanged for 2 mL of serum-free DMEM and cells were then stimulated with 100 ng/mL of LPS for 12 hours; control cells were without LPS stimulation. Afterwards, 1 μM of t-AUCB with or without 1 μM of the CYP inhibitor SKF 525A were added for 30 min. 10 μM of NADA or NA5HT were then added for 30 min with or without 10 μM or 30 μM AEA. Cells were scraped into media and combined with 2 mL ice-cold methanol. Cells were lysed using three consecutive 30-second on/off cycles on a water-bath sonicator. Cell debris was pelleted via centrifugation and the supernatant was purified using 100-mg Bond Elut C-18 cartridges (Varian, Harbor City, CA). Elution fractions were dried under reduced pressure, resuspended in 150 μL of 180-proof ethanol, and analyzed as stated above for tissue extractions. To account for batch-to-batch variability, data in the presence of AEA were analyzed based on a percentage to controls without AEA.

### TRPV1 binding/activation measurements

Binding of NADA, NA5HT, and epoxy-eVDs to TRPV1 was determined using an intracellular Ca^2+^ fluorescent quantification method. HEK-hTRPV1 cells were grown for 3 passages after recovery from frozen stocks before plating on Corning CellBind black, clear-bottom 96-well fluorescence plates coated with poly-L-lysine. After 24 hrs, media was removed, and cells were loaded with 3 μM Fura-2 AM dye (Molecular Probes) in sterile-filtered HEPES-Tyrode Buffer (HTB) (Alfa Aesar) supplemented with 0.01% Plurionic F-127 (Molecular Probes) for 20 min at room temperature. Analytes were prepared from DMSO stocks in 150 μL HTB on separate 96-well plates so that <0.1% DMSO was introduced to the cells. Dye was removed and cells were washed twice with HTB and 100 μL of HTB was added to the cells for the assay. To confirm binding to TRPV1, 0.5 μM of the TRPV1-specific antagonist AMG-9810 was added to this 100 μL of HTB prior to stimulating with agonists. Cells were then incubated at room temperature for 20 min to allow for the de-acetylation of the dye. Fluorescence readings were conducted on a SpectraMax Gemini EM (Molecular Devices, San José, Ca) plate reader using the following settings: bottom-read; channel 1—340 nm excitation, 510 nm emission; channel 2—380 nm excitation, 510 nm emission; 2-sec mix before experiment; read every 14 sec; 5-min experiment. The assays were conducted at room temperature (25°C). 100 μL of agonists were transferred in triplicate via multi-channel pipette to initiate the assay and the fluorescence intensities of both channels were measured over 5 min. The intensity from channel 1 (Ca^2+^-bound Fura-2) was divided by the intensity from channel 2 (Ca^2+^-free Fura-2) to achieve the Fluorescence Ratio 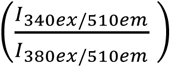. The Fluorescence Ratio was then plotted over time. Due to variations in the activation, the AUC of the Fluorescence Ratio from 84-252 seconds was used to determine activation. The average AUC of DMSO from 84-252 seconds was considered “baseline” and subtracted from each data point. The AUCs were plotted as a function of concentration and fitted to a dose-response curve (Equation 5) using OriginPro. CAP was used as a full-agonist positive control, and the *B*_*max*_ of the CAP was defined as 100% activation. Antagonism experiments for NA5HT, 14’,15’-epoNA5HT, and AMG-9810 were determined by adding varying concentrations of antagonist in 100 μL of HTB 15 min prior to stimulating with 250 nM CAP.

### Live cell Ca^2+^ imaging

Cultured DRG neurons were loaded with 4 μM Fura-2 AM (Life Technologies) in culture medium at 37°C for at least 60 minutes before use. Cells were washed 3 times and incubated in HBSS at room temperature for 30 minutes. Fluorescence at 340 nm and 380 nm excitation wavelengths was recorded on an inverted Nikon Ti-E microscope equipped with 340-, 360-, and 380-nm excitation filter wheels with NIS-Elements imaging software (Nikon Instruments Inc.). Fura-2 ratios (F340/F380) reflect changes in [Ca^2+^]_i_ upon stimulation. Values were obtained from 50–100 cells in time-lapse images from each coverslip. Threshold of activation was defined as 3 SD above the average (∼20% above the baseline).

### Whole-cell Patch-clamp recording

Whole-cell patch-clamp recordings were performed using an Axon 700B amplifier (Molecular Devices, Sunnyvale, CA, USA) with Clampex 10.4 software (Molecular Devices). at room temperature (22–24°C) on the stage of an inverted phase-contrast microscope equipped. Pipettes were pulled from borosilicate glass (BF 150-86-10; Sutter Instrument, Novato, CA, USA) with a Sutter P-1000 pipette puller, which had resistances of 2–4 megaohms when filled with pipette solution containing 140 mM KCl, 1 mM EGTA, 1 mM MgCl_2_, 5 mM MgATP and 10 mM HEPES with pH 7.3 and 320 mOsm/L osmolarity. Cells were continuously perfused with extracellular solution containing the following: 140 mM NaCl, 2 mM CaCl2, 1 mM MgCl2, 5 mM KCl, 10 mM glucose and 10 mM HEPES, pH adjusted to 7.4 with NaOH and the osmolarity was adjusted to ≈340 mOsm/L with sucrose. Data were analyzed and plotted using Clampfit 10 (Molecular Devices).

### PRESTO-TANGO binding with CB1 and CB2

Binding of eVDs to CB1 and CB2 was performed in HTLA cells using the PRESTO-TANGO assay as previously described^57^. HTLA cells, (an HEK293 cell line stably expressing a tTA-dependent luciferase reporter and a β-arrestin2-TEV fusion gene) were a gift from Brian Roth’s lab. CP-55940 was used as a full-agonist positive control for both receptors, and its *B*_*max*_ was defined as 100% activity. Cells were incubated with 1 μM of t-AUCB for 30 minutes prior to stimulation with analyte for 8 hrs. Luminescence readings were then recorded and processed as previously described.^57^ For antagonism experiments, analytes were co-administered at varying concentrations with 50 nM CP-55940.

### Quantitation of 14’,15’-epoNADA and 14’,15’-epoNA5HT using LC-MS/MS

Samples were analyzed with the 5500 QTRAP LC/MS/MS system (Sciex, Framingham, MA) in Metabolomics Lab of Roy J. Carver Biotechnology Center, University of Illinois at Urbana-Champaign. Software Analyst 1.6.2 was used for data acquisition and analysis. The 1200 series HPLC system (Agilent Technologies, Santa Clara, CA) includes a degasser, an autosampler, and a binary pump. The LC separation was performed on an Agilent SB-Aq (4.6 x 50mm, 5μm) with mobile phase A (0.1% formic acid in water) and mobile phase B (0.1% formic acid in MeCN). The flow rate was 0.3 mL/min. The linear gradient was as follows: 0-1min, 90%A; 8-13min, 0%A; 13.5-18min, 90%A. The autosampler was set at 10°C. The injection volume was 10 μL. Mass spectra were acquired under positive electrospray ionization (ESI) with the ion spray voltage of +5000 V. The source temperature was 450 °C. The curtain gas, ion source gas 1, and ion source gas 2 were 30, 65, and 55, respectively. Multiple reaction monitoring (MRM) was used for quantitation: 14’,15’-epoNA5HT m/z 479.3 → m/z 160.0; 14’,15’-epoNADA m/z 456.3 → m/z 137.1. Internal standard Anadamide-d4 was monitored at m/z 352.3 → m/z 287.2.

### Quantitation of EET-EAs and either 14’,15’-epoNADA or 14’,15’-epoNA5HT using LC-MS/MS

Samples were analyzed with the 5500 QTRAP LC/MS/MS system (Sciex, Framingham, MA) in Metabolomics Lab of Roy J. Carver Biotechnology Center, University of Illinois at Urbana-Champaign. Software Analyst 1.6.2 was used for data acquisition and analysis. The 1200 series HPLC system (Agilent Technologies, Santa Clara, CA) includes a degasser, an autosampler, and a binary pump. The LC separation was performed on an Agilent Agilent Eclipse XDB-C18 (4.6 x 150mm, 5μm) with mobile phase A (0.1% formic acid in water) and mobile phase B (0.1% formic acid in MeCN). The flow rate was 0.4 mL/min. The linear gradient was as follows: 0-2min, 90%A; 8min, 55%A; 13-25min, 40%A; 30min, 30%A; 35min, 25%A; 36-44min, 0%A; 45-50min, 90%A. The autosampler was set at 10°C. The injection volume was 10 μL. Mass spectra were acquired under positive electrospray ionization (ESI) with the ion spray voltage of +5000 V. The source temperature was 450 °C. The curtain gas, ion source gas 1, and ion source gas 2 were 32, 65, and 55, respectively. Multiple reaction monitoring (MRM) was used for quantitation: 14,15-epoNA5HT m/z 479.3 → m/z 160.0; 14,15-epoNADA m/z 456.3 → m/z 137.1; 5,6-EET-EA, 8,9-EET-EA, 11,12-EET-EA, and 14,15-EET-EA are all measured with m/z 264.2 → m/z 62.0. Internal standards Anadamide-d4 and 14,15-EET-EA-d8 were monitored at m/z 352.3 → m/z 287.2 and m/z 372.2 → m/z 63.0, respectively.

### Binding equations

The general one-site binding equation used was Equation 1 below

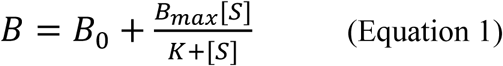

where *B*_*o*_ is the baseline response, *B*_*max*_ is the maximum response, *K* is the binding parameter, and [*S*] is the substrate concentration. For kinetic experiments, *B* and *K* represent velocity and *K*_*m*_, respectively; for Soret binding experiments, *B* and *K* represent Δ*A* and *K*_*D*_, respectively. For metabolism and Soret experiments, *B*_*o*_ = 0; for NADPH metabolism *B*_*o*_ is a nonzero value.

Inhibition experiments were described by either a competitive model (Equation 2) or noncompetitive model (Equation 3)

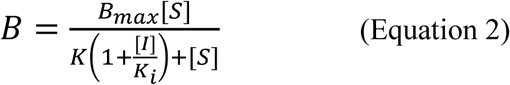

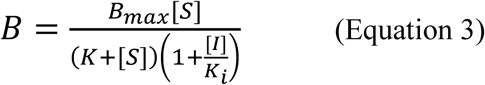

where *K*_*i*_ is the affinity of the inhibitor and [*I*] is the concentration of the inhibitor.

A general two-site binding equation (Equation 4) was used to describe the metabolism of NADA and NA5HT

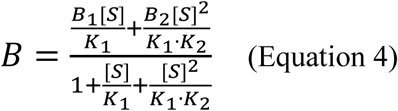

where *B*_1_ and *B*_2_ are the maximum metabolism at the first site and second site, respectively, and *K*_1_ and *K*_2_ are the affinities at the first site and second site, respectively.

*In cellulo* data were fitted to a dose-response equation (Equation 5)

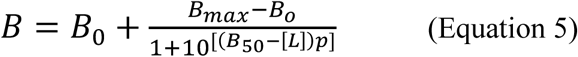

where [*L*] is the concentration of the ligand, *p* is the Hill coefficient, *B*_50_ is the half-maximal response (EC_50_ for agonism and IC_50_ for antagonism experiments), *B*_*o*_ is the baseline response (bottom asymptote), and *B*_*max*_ is the maximum response (top asymptote).

### Statistical analysis

Unless stated otherwise, statistical significance was determined by a one-way ANOVA followed by a Tukey’s post-hoc analysis. P values < 0.05 were considered statistically significant.

### Modeling and simulation of membrane-bound CYP2J2 with AEA and NADA or NA5HT

Initial structural models of membrane-bound CYP2J2 bound to AEA and NADA or NA5HT were generated with molecular docking performed with AutoDock Vina^61^ in a stepwise manner as described below. A grid box of dimension 22 Å in x, y and z and centered in the active site of CYP2J2 was employed for docking. We first docked NADA or NA5HT to our previous membrane-bound models of CYP2J2 ^62,63^. From this docking step, configurations of AEA/NADA/NA5HT in which the main epoxidation site (carbons C14 and C15) were close to the heme moiety (with distance < 5 Å) and with a high docking score, resulting in over 100 initial structures of each molecule in complex with CYP2J2. The resulting structures were then employed as receptors for docking of a second molecule (AEA for NADA/NA5HT in active site, or NADA/NA5HT for AEA in active site). This second step allowed us to explore potential peripheral binding sites (i.e., outside the central active site cavity of CYP2J2), which were hypothesized from the experimental. From docking, two potential peripheral binding sites were identified. For each peripheral binding configuration, the corresponding CYP2J2 complexes with two molecules were sorted by docking score, and the models with the highest score were employed as starting configurations for MD simulations. Each simulation system was minimized for 2,000 steps, and equilibrated for 1 ns with the C_α_ atoms of CYP2J2 and the heavy atoms of the ligands (AEA, NADA and NA5HT) harmonically restrained (with force constant k=1 kcal/mol/Å^2^). Following this preparation step, the 2-molecule systems were simulated for 50 ns.

### Simulation protocol

The simulations were performed using NAMD2^64^. The CHARMM27 force field with cMAP^65,66^ corrections was used for CYP2J2. The CHARMM36^67,68^ force field was used for lipids. Force field parameters for AEA, NADA and NA5HT were generated by analogy from the CHARMM General Force Field^69^. The TIP3P model was used for water^70^. Simulations were performed with the NPT ensemble with a time step of 2 fs. A constant pressure of 1 atm was maintained using the Nosé-Hoover Langevin piston method^71,72^. Temperature was maintained at 310 K using Langevin dynamics with a damping coefficient γ of 0.5 ps^-1^applied to all atoms. Nonbonded interactions were cut off at 12 Å, with smoothing applied at 10 Å. The particle mesh Ewald (PME) method^73^ was used for long-range electrostatic calculations with a grid density of > 1 Å^-3^.

### Data availability statement

All plasmids used in the study are available from Addgene or can be obtained on request. All other data or resources are available in the Source Data file or from the corresponding author upon reasonable request.

## RESULTS

### Biosynthesis of epoxy-endovanilloids by microglial cells and their endogenous levels in porcine brains

We first determined the levels of NADA and NA5HT in porcine brain, as pigs and humans express similar CYP isozymes. To achieve this, we synthesized the authentic standards (Figure 2a). As the epoxidation of PUFAs and eCBs by CYPs occurs mainly on the terminal alkene^57,59,62,74^, we synthesized 14’,15’-epoNADA and 14’,15’-epoNA5HT (Figure 2a) and separated them using a reverse-phase HPLC-UV method and confirmed the structures using NMR and LC-MS/MS (Figure S1-S5). As shown in Figure 2b Method 1, we developed an LC-MS/MS quantitation method in the selected reaction monitoring (SRM) mode to quantify NADA, NA5HT, 14’,15’-epoNADA, 14’,15’-epoNA5HT, and AEA from brain tissue and microglial cells. Previously, NADA was found in the cerebellum and hippocampus/thalamus regions in bovine and rat brains^46,75,76^, while NA5HT was identified in bovine brain extracts and in the intestine^47,48^. The levels of NADA and NA5HT have been reported to be <1-10 pmol · g^-1^ in these brain tissues, which is lower than literature AEA levels (∼20-80 pmol · g^-1^)^77^ and lower than our reported levels for AEA in rat brains (150 pmol · g^-1^) and pig brains (2800 pmol · g^-1^)^57^. It is important to note that AEA levels are known to rise postmortem due to denaturation of the enzymes that degrade AEA^78^. Additionally, during tissue extraction, NADA and NA5HT are unstable due to the potential oxidation of the headgroups. Hence, the levels of the NADA and NA5HT are expected to be lower than AEA.

**Figure 2.**
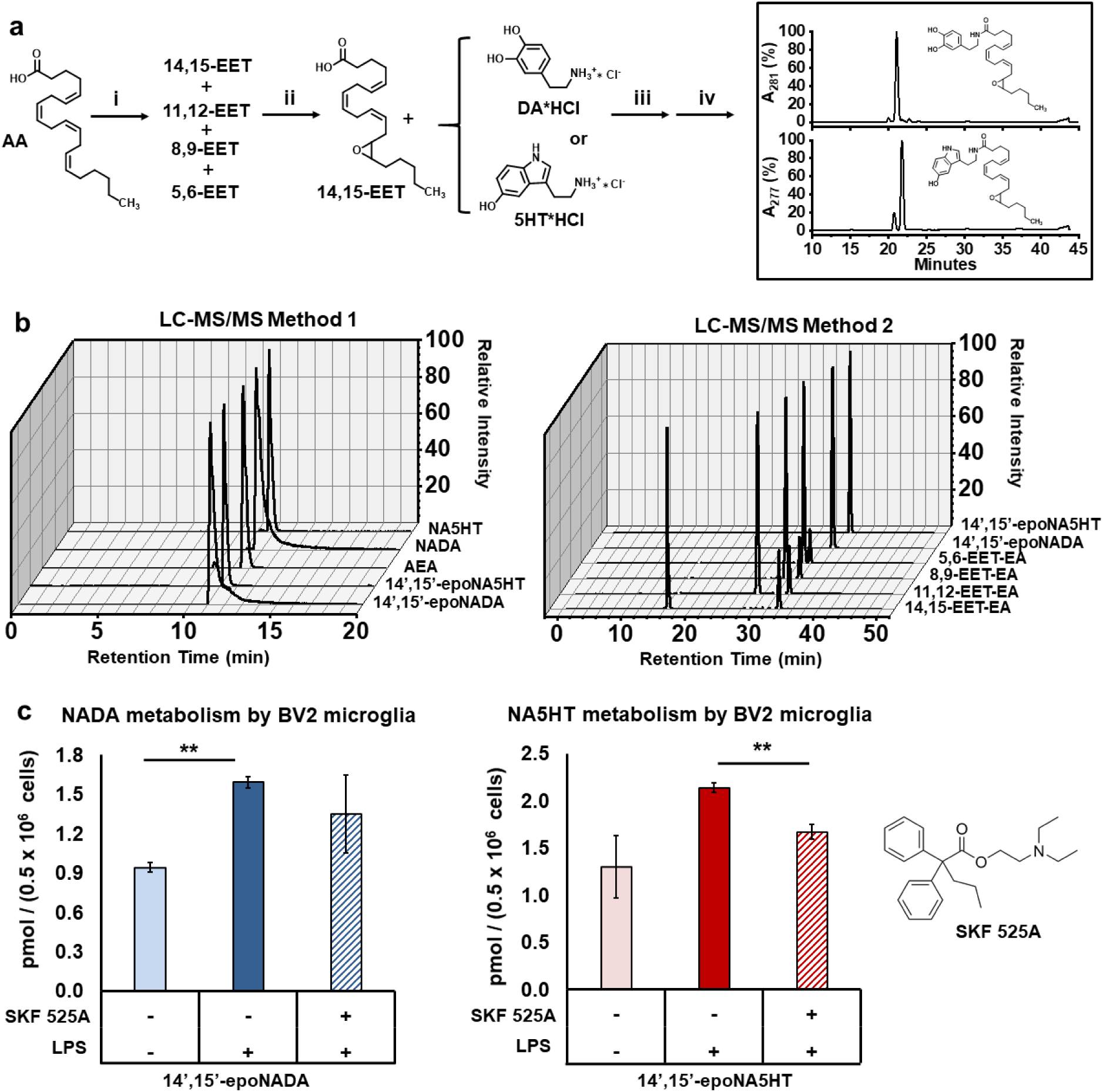
Synthesis of authentic standards, targeted lipidomics method, and production of epo-NADA and epo-NA5HT by microglial cells. **(a)** Authentic standards 14’,15’-epoNADA and 14’,15’-epoNA5HT of were synthesized. (i) *m-*CPBA, RT, 1 hr, MeCN (ii) reverse-phase HPLC purify (iii) EDC, DMAP, DIPEA; ice bath 1 hr; RT 8 hrs; 50:50 DCM:DMF (iv) reverse-phase HPLC purify. **(b)** Development of LC-MS/MS method for the separation of NADA, epo-NADA, NA5HT, epo-NA5HT and AEA (Method 1) and the different regioisomers of EET-EAs and epo-NADA and epo-NA5HT (Method 2). AEA m/z 348.3 → m/z 203.2; NADA m/z 440.2 → m/z 287.1; 14’,15’-epoNADA m/z 456.3 → m/z 137.1; 14’,15’-epoNA5HT m/z 479.3 → m/z 160.1; NA5HT m/z 463.3 → m/z 287.2; EET-EA m/z 264.2 → m/z 62.0. **(c)** NADA and NA5HT metabolism by BV2 microglial cells under lipopolysaccharide (LPS) stimulation in the presence of 1 µM t-AUCB (sEH inhibitor) and 10 µM NADA or NA5HT. The reversible CYP epoxygenase inhibitor SKF 525A was used to demonstrate CYP-mediated metabolism. Data represents the mean ± SEM of 3 experiments. ** p < 0.01

Herein, we measured 1.7 and 1.9 pmol of NADA (0.182 ± 0.081 pmol · g^-1^ wet tissue) and 11.3 and 4.9 pmol of NA5HT (0.686 ± 0.006 pmol · g^-1^ wet tissue) from the cerebella of two pig brains; and although they were also detected in two more pig brains the values were very low. From the hippocampus-thalamus-hypothalamus regions, we recovered 0.4 and 0.5 pmol of NADA (0.039 ± 0.009 pmol · g^-1^ wet tissue) and 0.2 and 0.1 pmol of NA5HT (0.010 ± 0.003 pmol · g^-1^ wet tissue) in two brains. The epoxy-eVDs were variable and often below detection limit likely due to the low abundance of the parent molecules. Overall, the levels of NADA and NA5HT are much less than AEA in rat and pig brains reported from our laboratory^57^.

Additionally, the levels of the NADA and NA5HT were variable in the four pig brains that were analyzed. Of note, two extraction methods were used to detect eVDs and AEA (methanol/C-18 cartridges for eVDs, acetate:hexanes/silica column for AEA^57^). AEA was very poorly detected using the methanol/C-18 cartridges and the eVDs quickly degraded on during the acetate:hexanes extraction and silica column purification. Therefore, we cannot simultaneously quantify AEA and eVDs in the same sample.

In the field of lipid metabolites, there is strong evidence that the production of lipid metabolites is localized to the site of action and that there is a rapid subsequent degradation^79^. This leads to low plasma/tissue levels of most lipid metabolites. Hence, we studied the epoxidation of these lipid metabolites in microglial cells. Microglial cells, the innate immune cells (macrophages) of the brain, are activated during neuroinflammation and play important roles in pain modulation^80,81^. Previously, it was demonstrated that CYPs are upregulated during neuroinflammation^82^. Hence, we used BV2 microglial cells to determine the production of eVD epoxides from NADA or NA5HT under an inflammatory stimulus. We stimulated microglial cells with lipopolysaccharide (LPS), followed by the addition of NADA or NA5HT^57^. We found that within 30 minutes of incubation, 14’,15’-epoNADA and 14’,15’-epoNA5HT were formed under LPS stimulation. Interestingly, these molecules are also produced without LPS stimulation showing that these molecules are made spontaneously by resting microglial cells (Figure 2c). Importantly, the production of these metabolites were partially inhibited in the presence of SKF 525A (a reversible CYP epoxygenase inhibitor) demonstrating that the eVD epoxides are produced partly by enzymatic oxidation by CYP epoxygenases^83^ (Figure 2c). SKF 525A is a nonspecific CYP inhibitor that is known inhibits CYP450-dependent arachidonic acid conversion to EET metabolites. However, there are several other CYPs in the microglial cells that may be producing these epoxidized metabolites, which can explain the partial inhibition.

### Epoxygenation of NADA or NA5HT by BV2 microglia in the presence of AEA

Several studies have shown that the endogenous levels of AEA are much higher than NADA and NA5HT. Furthermore, it has been shown that various brain CYPs such as CYP2J2 and CYP2D6 convert AEA into AEA epoxides (EET-EAs). Therefore, to understand the substrate specificity of CYPs when both substrates (AEA and NADA or NA5HT) are present, we further used the activated microglial cells to study the co-metabolism of AEA with NADA or NA5HT (Figure 3a and b). Interestingly, we observed that AEA potentiated the formation of 14’,15’-epoNADA (∼ 2-fold) in a concentration-dependent manner (Figure 3a). Contrariwise, AEA inhibited 14’,15’-epoNA5HT formation (∼0.5-fold) (Figure 3b). It is possible that AEA acts as a potentiator of CYP-mediated NADA metabolism and an inhibitor of CYP-mediated NA5HT metabolism, either directly or indirectly. Several ligands have shown to either act as direct potentiators or inhibitors of CYP metabolism through cooperative or allosteric binding^84-86^. To explore this possibility, we delineated the mechanism of eVD metabolism using a recombinantly expressed CYP.

**Figure 3.**
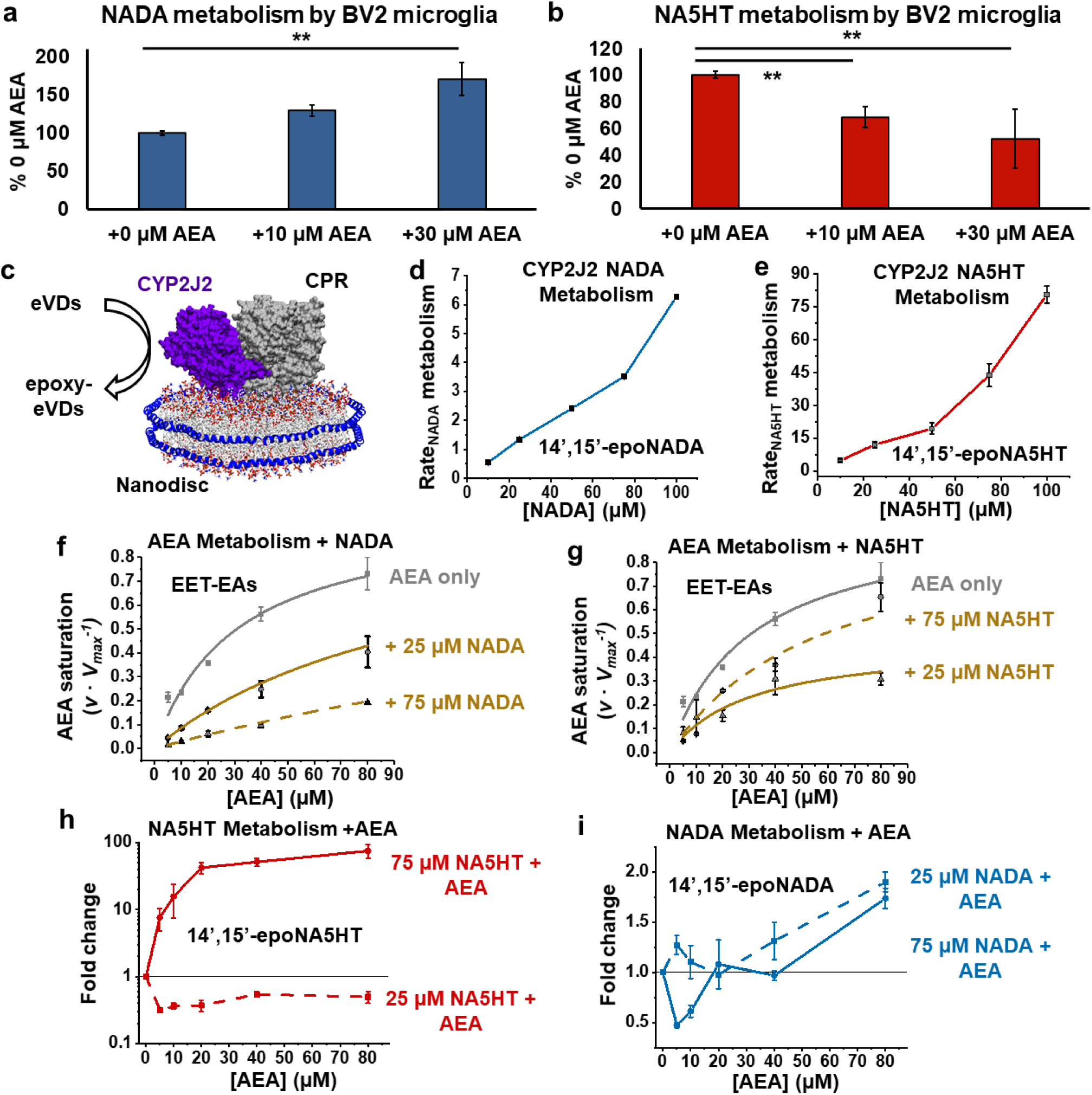
Potentiation of eVD metabolism by AEA. **(a)** Metabolism 10 µM NADA and **(b)** 10 µM NA5HT in the presence of increasing concentrations of AEA by BV2 microglia. Metabolism as conducted similarly to data in Figure 2c. Data shown are the mean ± SEM of *n* = 6 (two sets of triplicate performed on separate days). **p < 0.01. **(c)** Schematic of the CYP2J2-CPR-Nanodisc. **(d-e)** Metabolism of NADA and NA5HT by CYP2J2-CPR-NDs. Rates are in pmol_epoxy-eVD_ · min^-1^ · nmol_CYP2J2_^-1^ and data represents the mean ± SEM of 3 experiments. **(f-i)** Co-substrate metabolism of eVDs with AEA. Data represents the mean ± SEM of multiple replicates. **(f)** AEA metabolism in the presence of NADA and **(g)** NA5HT. Rates are shown as the fraction of the *V*_*max*_ (135 pmol_EET-EAs_ · min^-1^ · nmol_CYP2J2_^-1^) for AEA metabolism (grey squares) as previously published (see text). **(h)** Fold change of NA5HT metabolism and **(i)** NADA metabolism in the presence of increasing AEA concentrations compared to the metabolism without AEA.

Of the common CYP epoxygenases in mouse, CYP2J9 and CYP2J12 were shown to be highly expressed in mouse brain tissues while CYP2Cs showed low expression^87,88^. We confirmed that CYP2J9 and CYP2J12 are expressed in the BV2 microglial cells used for our studies (Figure S6). CYP2J9 and CYP2J12 are highly homologous to human CYP2J2, which is the only CYP2J found in humans. CYP2J2 is also the most highly expressed CYP epoxygenase in the human brain^89-91^. Hence, we elucidated the biochemistry of eVD metabolism in the presence and absence of AEA by human CYP2J2.

### Human CYP2J2 epoxidizes NADA and NA5HT to form epoNADA and epoNA5HT

CYP2J2 is highly expressed in the brain, cardiovascular, and cerebrovascular systems and is one of the major epoxygenases in these tissues^89-92^. Additionally, CYP2J2 is known to epoxidize several endocannabinoids including AEA^57,60,93^. Therefore, we recombinantly expressed human CYP2J2 and its redox partner cytochrome P450 reductase (CPR) and studied the metabolism of eVDs by CYP2J2.

In order to study the direct metabolism of NADA and NA5HT, we incubated CYP2J2 with each eVD and detected the metabolites using UV-Vis HPLC and LC-MS/MS. All of the oxidized products of NADA and NA5HT are detailed in the supplementary material (Figures S7-S26 and Table S2). The metabolism of NADA resulted in the formation of eight mono-oxygenated products (PD1-8) (Figure S7-S9 and Table S2). We determined that 14’,15’-epoNADA (PD5) is a product based on the fragmentation and co-elution with the synthesized standard. Two di-oxygenated products were also found (PD9 and PD10) (Figure S10). Additionally, a mass corresponding to a hydroxyquinone product of NADA (PD11) was detected within 6 ppm of the expected mass (Figure S11). UV-Vis HPLC demonstrates that PD11 has an absorbance profile between 270 and 310 nm that is typical of hydroxyquinones (Figure S11c). It is known that dopamine is oxidized to form 6-hydroxy-*p*-quinone,^94,95^ and therefore, NADA-6-hydroxy-*p*-quinone is likely the identity of PD11 (Figure S7c).

The metabolism of NA5HT by CYP2J2 produced 4 mono-oxygenated products (PS1-4) (Figure S12-S14 and Table S2). PS2 was confirmed to be 14’,15’-epoNA5HT. PS3 and PS4 appear to be oxidized at the 5HT headgroup (Figure S13d-e and Figure S14b-c). Acyl-chain di-oxygenated products and quinonized products of the 5HT headgroup were also observed (Figures S15-S18). To confirm these metabolites of eVDs, we also investigated CAP as a substrate of CYP2J2, and found headgroup-oxidized products among other oxygenated products (Figures S19-S26 and Table S2). After establishing eVDs as substrates of CYP2J2, we then proceeded to measure the kinetics of metabolism by CYP2J2-CPR-Nanodiscs.

### Kinetics of NADA and NA5HT metabolism by CYP2J2-CPR-Nanodiscs

We next incorporated CYP2J2 and CPR into stable nanoscale lipid bilayers known as Nanodiscs (ND) and proceeded to determine the kinetics of the 14’,15’-epoxide formation (Figure 3c). The metabolism of NA5HT by CYP2J2 is in a similar range as the metabolism of AEA and other lipid PUFAs,^59,62^ but the metabolism of NADA is low. Interestingly, the data demonstrate the presence of multiple binding sites, as the kinetics plots strongly deviate from a typical Michaelis-Menten model. The plots resemble the beginning of a sigmoidal curve indicating positive binding interactions. However, saturation was not achieved as the eVDs are poorly metabolized and insoluble beyond 100 µM (Figure 3d and e). We therefore hypothesized that there are at least two binding sites. To further probe the kinetics of eVD metabolism, we measured the rate of NADPH oxidation by CYP2J2-CPR-ND in the presence of the eVDs.

CPR shuttles electrons from NADPH to CYPs to facilitate the metabolism of substrates. Therefore, the rate of NADPH oxidation increases in the presence of CYP substrates. In this case, NADPH oxidation in the presence of NA5HT showed biphasic kinetics (Figure S27). During Phase I, the rates are close to the baseline (without NA5HT). In Phase II (after one minute) the rates increase to produce a second linear phase that demonstrate a hyperbolic curve as a function of the NA5HT concentration (Equation 1) (Figure S27). This increase in NADPH oxidation occurs only when CYP2J2 is included in the reaction (Figure S27c). The *K*_*m*_ and *V*_*max*_ of Phase II are 27.3 ± 11.3 µM and 178 ± 28 nmol_NADPH_ · min^-1^ · nmol_CYP2J2_^-1^, respectively.

Therefore, these data suggest NA5HT binds to two sites, where one is silent for the formation of 14’,15’-epoNA5HT and the other increases NADPH oxidation. Although the drug ebastine (EBS) shows monophasic NADPH oxidation with similar kinetics as NA5HT (*K*_*m*_ = 18.9 ± 9.5 µM and *V*_*max*_ = 64.8 ± 12.1 nmol_NADPH_ · min^-1^ · nmol_CYP2J2_^-1^), AA, AEA, and NADA do not increase NADPH oxidation and therefore the biphasic NADPH kinetics is unique to NA5HT (Figure S27a).

### Relative binding affinities of NADA and NA5HT as determined by ebastine (EBS) binding inhibition

Substrate binding to CYPs is typically measured by observing direct spectroscopic changes to the heme Soret band at 417 nm. As eCBs do not produce substantial Soret shift upon binding, we used an EBS competitive inhibition assay to measure the binding affinity of NADA and NA5HT^61,64^. Both NADA and NA5HT displayed competitive inhibition of EBS binding (Equation 2) suggesting that the binding of NADA and NA5HT overlap the binding of EBS (*K*_*i*_ for NADA is 71.1 ± 20.0 μM and *K*_*i*_ for NA5HT is 49.3 ± 6.2 μM).

### Co-metabolism of AEA with NADA or NA5HT

We next determined the co-substrate kinetics of AEA with the eVDs to determine if we can explain the observed effects of AEA on BV2-mediated metabolism. We first developed an LC-MS/MS method to simultaneously measure the four different regioisomers of AEA epoxides (EET-EAs) and either 14’,15’-epoNADA or 14’,15’-epoNA5HT (Figure 2b, Method 2). We had previously determined the kinetics of AEA metabolism by CYP2J2-CPR-NDs^59^, and we repeated these experiments using two concentrations (25 and 75 μM) of either NADA or NA5HT. NADA inhibited AEA metabolism following a competitive inhibition model (Figure 3f). A 3D global fit of the data (Figure S28) yields a *K*_*i*_ of 7.50 ± 0.88 μM for the inhibition of AEA by NADA, which is among the strongest endogenous inhibitors of CYP2J2 as compared to virodhamine^96^. NA5HT was a noncompetitive inhibitor (Equation 3) at 25 μM (*K*_*i*_ = 21.4 ± 3.6 μM) and a competitive inhibitor at 75 μM (*K*_*i*_ = 86.6 ± 18.5 μM) (Figure 3g). NADA and NA5HT also altered the regioselectivity of AEA epoxidation in a concentration-dependent manner (Figure S29).

Interestingly, AEA showed a biphasic potentiation of NADA and NA5HT metabolism (Figure 3h and i) when we measured the epoxy-eVD formation. With increasing concentrations of AEA, the metabolism of NADA and NA5HT slightly decreases but then increases at higher concentrations of AEA. AEA significantly and hyperbolically increases 75 μM NA5HT metabolism by almost 100-fold with an apparent *V*_*max*_ and *K*_*m*_ of 5.46 ± 1.91 nmol · min^-1^ · nmol_CYP2J2_^-1^ and 53.7 ± 35.7 μM, respectively (Figure 3h). NADA is potentiated by almost 200% at the highest concentrations of AEA (Figure 3i). Overall, the potentiation of eVD metabolism by AEA and the altered AEA regioselectivity in the presence of eVDs demonstrate that eVDs and AEA are binding to CYP2J2 at multiple sites. Furthermore, these data support the observed crosstalk of AEA and eVDs in microglial cells (Figure 3a and b), as at similar concentrations within each experiment AEA potentiates NADA and inhibits NA5HT.

### Molecular dynamics (MD) simulations demonstrate AEA stabilizes eVD binding to CYP2J2

In order to characterize the molecular basis of the multi-site kinetics observed with the eVDs, we performed MD simulations starting from membrane-bound structures of CYP2J2 in complex with substrates (AEA, NADA or NA5HT). Initial molecular docking was performed with AEA and either NADA or NA5HT in a stepwise manner^97^. These models allowed us to probe the binding mode of a second molecule to CYP2J2 in the presence of another molecule bound in the active site in an unbiased manner (i.e., without any assumptions about location of peripheral binding pockets). Two distinct configurations of peripheral AEA binding, with either NADA or NA5HT in the active site, were identified (Figure 4a and b). In Configuration 1, AEA was docked in a pocket located below the I-helix, with its ethanolamine group near a residue (R321) that we have previously identified to modulate PUFA binding^62^ (Figure 4a). In Configuration 2, AEA was located closer to the membrane interface (Figure 4b). For the two identified AEA binding configurations, the initial orientation of NADA or NA5HT in the active site was similar, with the main epoxidation site (carbons C14 and C15) close to the heme moiety (distance < 5 Å), and the headgroup (DA or 5HT) pointing away from the heme. These docking results suggested that NADA/NA5HT binding was not modulated by the same PUFA-interacting residues previously reported (T318, R321 and S493)^62^. Due to the larger headgroups of NADA/NA5HT compared to other PUFAs (i.e., DA/5HT vs. carboxylic acid), a different binding orientation (not interacting with the PUFA triad) was necessary to fit these molecules in the active site.

**Figure 4.**
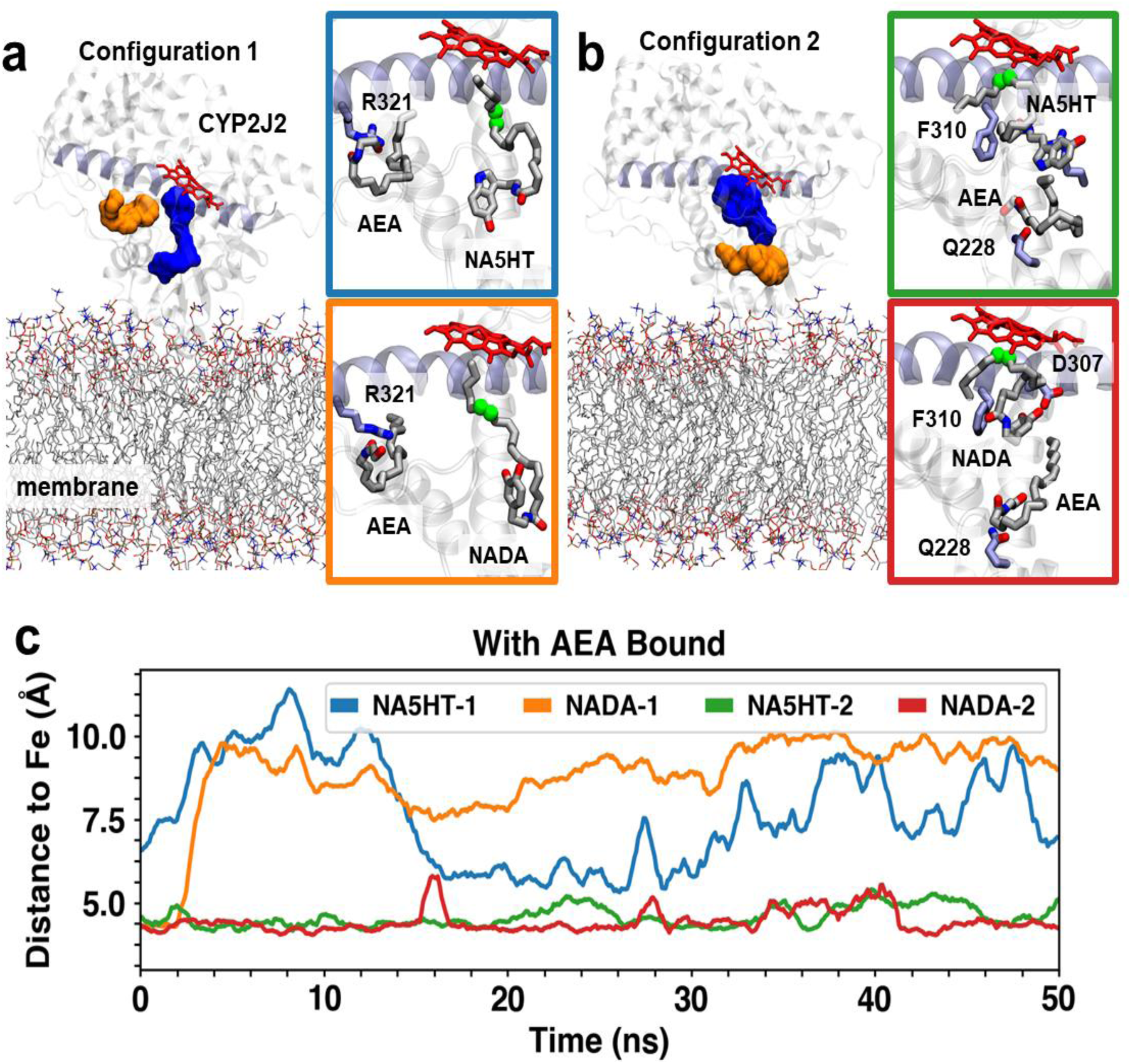
Concurrent CYP2J2 binding of AEA and NADA/NA5HT in MD simulations. Representative snapshots of two distinct configurations of AEA and NADA/NA5HT identified with molecular docking and with MD simulations: **(a)** peripheral binding pocket located near helix I (shown in purple cartoon) and **(b)** peripheral binding site near the membrane interface. In both panels, peripheral and active binding site cavity are shown as orange and blue surfaces, respectively. Lipids are shown in stick representation and CYP2J2 in cartoon representation. AEA, NADA, and NA5HT molecules are shown as sticks. Notable residues involved in AEA/NADA/NA5HT interactions are also shown as sticks. Carbons involved in epoxidation are highlighted as green spheres. **(c)** Time series of carbon-to-heme distances obtained from 50 ns MD simulations for the main epoxidation sites of NADA/NA5HT. Colors correspond to the subpanels shown in **(a)** and **(b)**.

The MD simulations (Figure 4c, Tables S3-S6, and Movies S1-S4) revealed that stable NADA/NA5HT binding (i.e, with the epoxidation site distance to the heme < 5 Å) was only achieved when AEA is bound in Configuration 2 (Figure 4b). When AEA is located in the I-helix pocket (Configuration 1), NADA or NA5HT in the active site gradually move away from the heme, which results in a displacement of its epoxidation site (with the distance to heme between 7.5 and 10 Å during the simulations) (Figure 4c). In contrast, AEA in Configuration 2 constrains the motion of NADA or NA5HT in the active site, which maintain their potentially productive orientation close to the heme (epoxidation site to heme distance < 5 Å) during the simulation (Figure 4c). In these simulations, the ethanolamine group of AEA interacted with Q228, located near the membrane interface, and remained locked in its binding site. Positioning of AEA in turn constrained the mobility of the molecule in the active site (NADA or NA5HT). NADA/NA5HT are further stabilized by hydrophobic interactions (mainly with F310) and transient electrostatic interactions (i.e., NA5HT serotonin group with D307 and E222). These observations suggest that NADA/NA5HT binding is enhanced by concurrent AEA binding to a peripheral site near the membrane interface and provide insights into the protein residues involved in this binding (e.g., Q228 for AEA and F310 for NADA/NA5HT). Overall, the MD simulations in conjunction with the kinetics data concur with the observations from the cell culture studies that AEA enhances the metabolism of NADA.

### Estimate of NA5HT binding at the second site

We can definitively demonstrate that the eVDs bind to at least two sites in CYP2J2 from the kinetic curves; however, due to the solubility issues of the lipids, we cannot determine the kinetic parameters of their metabolism.

Notwithstanding, we can obtain approximate values for the kinetic parameters of NA5HT based on a two-site binding equation using assumptions from other data (Equation 4). The assumptions are the following. (A) Negligible metabolism occurs at the unproductive site (*B*_1_ = 0). (B) The EBS competitive binding and NADPH oxidation data represent the affinity at the first site and can be averaged to obtain *K*_1_ = 38 µM. (C) The apparent *V*_*max*_ as AEA potentiates the metabolism of NA5HT represents the *V*_*max*_ of NA5HT metabolism (i.e., *B*_2_ = 5,460 pmol · min^-1^ nmol_CYP2J2_^-1^). Substituting these values into Equation 4, *K*_2_ is determined to be 5.36 ± 0.27 mM (Figure S30). This shows that the affinity of the eVDs for the second site is weak. Due to a lack of kinetic details in the data, a similar analysis cannot be done for NADA, though the binding to the second site is likely as weak.

### Anti-inflammatory action of the epoxy-NADA and epoxy-NA5HT in microglial cells

After establishing how these metabolites are potentially formed, we proceeded to characterize their pharmacology. Primary sensory afferent neurons and immune cells are rich sources of lipid mediators that regulate pain due to inflammation. Previous studies have demonstrated the anti-inflammatory actions of eVDs^98-100^; thus, we hypothesize that epoxy-eVDs would also be anti-inflammatory. Microglial cells play an important role in neuroinflammation and neuropathic pain. Microglial cells are strongly activated after injury and release proinflammatory cytokines such as IL-6, IL-1β, and TNF-α. Therefore, there is a significant interest in discovering lipid-based molecules that decrease microglial activation. Herein, we show that the two novel lipid mediators, epoNADA and epoNA5HT, exhibit anti-inflammatory action in microglial cells. To investigate the actions of eVDs and epoxy-eVDs, we measured the levels of pro-inflammatory nitric oxide (NO), IL-6, IL-1β, and TNF-α in lipopolysaccharide- (LPS)-stimulated BV2 cells. All eVDs and epoxy-eVDs dose-dependently reduced NO and IL-6 production (Figure 5a-d) and the IC_50_ values are tabulated in Table 1. Together, these data demonstrate that the eVDs and the epoxy-eVDs are anti-inflammatory mediators. As determined by MTT and BrdU assays, these compounds were not toxic at their effective concentrations, though NA5HT and 14’,15’-epoNA5HT increased cell viability in the presence of LPS suggesting they may be pro-proliferative, which was confirmed using a BrdU assay (Figure S32).

**Table 1.** Anti-inflammatory marker inhibition and receptor activation parameters of eVDs and epoxy-eVDs.

|  | IC<sub>50</sub> (μM) |  | TRPV1 |  | CB1 |  | CB2 |  |
| --- | --- | --- | --- | --- | --- | --- | --- | --- |
|  | NO<br>production | IL-6<br>expression | %<br>Activation<sup>a</sup> | EC<sub>50</sub><br>(IC<sub>50</sub>)<sup>b</sup> | %<br>Activation<sup>c</sup> | EC<sub>50</sub><sup>b</sup> | %<br>Activation<sup>c</sup> | EC<sub>50</sub><sup>b</sup> |
| NADA | 0.909 ± 0.076 | 0.507 ± 0.065 | 83.8 ± 13.6 | 962 ± 249 | 83.9 ± 9.2 | 0.045 ± 0.01 | --- | DNB<sup>d</sup> |
| 14',15'-<br>epoNADA | 0.888 ± 0.135 | 0.807 ± 0.440 | 79.7 ± 9.8 | 156 ± 36 | 92.3 ± 11.2 | 0.080 ± 0.03 | --- | DNB<sup>d</sup> |
| NA5HT | 2.14 ± 0.06 | 1.06 ± 0.49 | --- | (7,650 ± 1,300) | --- | DNB<sup>d</sup> | 19.2 ± 3.4 | 8.80 ± 5.57 |
| 14',15'-<br>epoNA5HT | 2.07 ± 0.18 | 1.56 ± 0.22 | 23.6 ± 4.4<br>--- | 120 ± 27<br>(254 ± 38) | 99.9 ± 13.1 | 16.5 ± 11.7 | --- | DNB<sup>d</sup> |

<sup>a</sup> Percent activation compared to the $B_{max}$ of CAP
<sup>b</sup> nM
<sup>c</sup> Percent activation compared to the $B_{max}$ of CP-55940
<sup>d</sup> DNB: Does not activate. Compounds up to 1 μM in concentration do not agonize the receptor and do not antagonize 50 nM of CP-55940.

**Figure 5.**
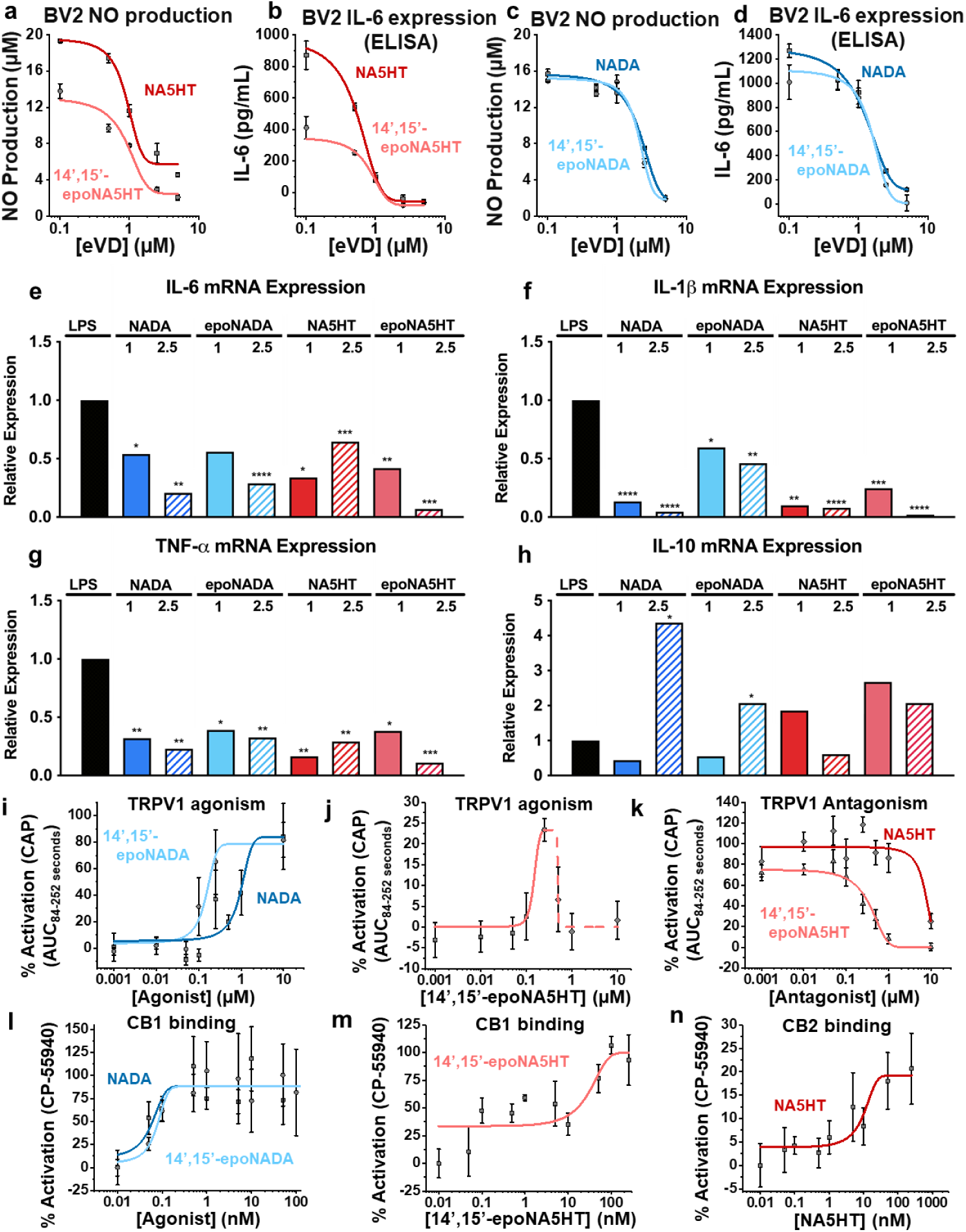
Anti-inflammatory actions of eVDs and receptor activation. **(a-d)** Anti-inflammatory effects of eVDs and epoxy-eVDs were determined by pre-incubating eVDs and epoxy-eVDs with BV2 cells for 4 hrs prior to stimulation by 25 ng/mL LPS. Data represents the mean ± SEM of *n* = 2-3. **(a)** Dose-dependent inhibition of nitric oxide (NO) production by NA5HT and 14,15-epoNA5HT as determined by Griess Assay. **(b)** Dose-dependent reduction of IL-6 protein expression (ELISA) by NA5HT and 14,15-epoNA5HT. **(c)** Dose-dependent inhibition of NO production and **(d)** IL-6 protein expression by NADA and 14,15-epoNADA. **(e-h)** Anti-inflammation experiments were repeated to measure pro-inflammatory *Il-6, Il-1β*, and *Tnf-α* and anti-inflammatory *Il-10* mRNA expression in the presence of LPS. Data represents the average of 3 technical replicates. Data is reported as the mean relative expression. Statistical analysis was performed using a two-tailed t-test (α = 0.05). P-values: 0.0332 (*), 0.0021(**), 0.0002 (***), and < 0.0001 (****). **(i-n)** eVD activation to TRPV1-transfected HEK cells was determined using a Fura 2-AM Ca^2+^-influx assay. The *B*_*max*_ of capsaicin (CAP) activation is defined as 100%. Data represents the mean ± SEM of *n* = 6 (two sets of triplicate experiments on separate days). **(i)** Antagonism was determined by preincubating cells with antagonist prior to stimulating with 250 nM CAP. **(j-l)** eVD activation of CB1 and CB2 was determined by the PRESTO-Tango assay. *B*_*max*_ of CP-55940 is defined as 100%. Data represents the SEM of n = 6 (two sets of triplicate experiments on separate days).

In addition to measuring IL-6 protein levels and NO, we also measured the mRNA expression of *Il-6, Il −10, Il* -*1β*, and *Tnf-α* to determine the transition of the pro-inflammatory phenotype of BV2 to the anti-inflammatory phenotype. Whereas IL-6, IL −1β, and TNF-α are pro-inflammatory, IL-10 is an anti-inflammatory cytokine that facilitates the resolution of inflammation. Concentrations around the IC_50_ of the IL-6 and NO experiments (1 and 2.5 µM of each compound) were tested and compared to cells with LPS only. All the eVDs and epoxy-eVDs downregulated pro-inflammatory *Il-6, Il-1β*, and *Tnf-α* with 14’,15’-epo-NA5HT being the most effective overall (Figure 5e-g). NADA and 14’,15’-epoNADA concomitantly upregulated *Il-10* (Figure 5h). NADA had the greatest effect at 2.5 μM and 14’,15’-epoNADA was second best. Therefore, the eVDs and epoxy-eVDs decrease *Il-6, Il-1β*, and *Tnf-α* expression and increase *Il-10* mRNA expression in activated microglial cells, demonstrating that these compounds are anti-inflammatory.

Most eCBs and eVDs mediate anti-inflammation and anti-pain polypharmacologically through CB1, CB2, and TRPV1 receptors. We determined that the mRNA of *Cnr1* (CB1 gene), *Cnr2* (CB2 gene), and *Trpv1* are expressed in the BV2 cells (Figure S33). Since these receptors are known targets of eVDs and mediate inflammation and pain, we proceeded to measure the activation of CB1, CB2, and TRPV1 by epoxy-eVDs.

### Activation of TRPV1 by eVDs and epoxy-eVDs

Previous studies have shown that NADA mediates anti-inflammation via TRPV1^98^. Therefore, we next investigated the actions of eVDs and epoxy-eVDs at TRPV1 using HEK cells stably expressing human TRPV1. As TRPV1 is a non-selective cation channel, activation was determined by measuring the relative Ca^2+^ influx (Figure S34 and Table 1). CAP was used as a full-agonist positive control and was found to have an EC_50_ of 6.88 ± 3.36 nM, in accordance with reported values^101^ (Figure S35a). We determined that NADA and 14’,15’-epoNADA are full agonists, with ∼80% activation compared to CAP (Figure 5i). The EC_50_ of 14’,15’-epoNADA is about 6-fold tighter than NADA. 14’,15’-epoNADA also promoted a broader duration of Ca^2+^ influx opening compared to NADA (Figure S34). The TRPV1-selective antagonist AMG-9810 was able to fully antagonize the signal from CAP, the eVDs, and the epoxy-eVDs, confirming the Ca^2+^ influx is TRPV1-mediated (Figure S35b and c).

On the contrary, 14’15’-epoNA5HT was found to be a partial agonist of TRPV1 up to 250 nM, with a 24% activation compared to CAP (Figure 5j). This signal was antagonized by AMG-9810, demonstrating it is TRPV1-mediated (Figure S35c). Concentrations above 250 nM, however, resulted in a reduction of the signal, signifying that 14’,15’-epoNA5HT is an antagonist of TRPV1 at higher concentrations (Figure 5j). Previously, NA5HT had been shown to be an antagonist of TRPV1^51^; therefore, we measured the antagonism of TRPV1 by 14’15’-epoNA5HT and compared it to NA5HT. 14’15’-epoNA5HT functioned as an antagonist of CAP at all concentrations, with an IC_50_ of 250 ± 38 nM (Figure 5k). Of note, this IC_50_ correlates to the concentration at which the self-antagonism was observed for 14’,15’-epoNA5HT (Figure 5j). From Table 1, we see that 14’15’-epoNA5HT is a 30-fold stronger antagonist of TRPV1 than NA5HT.

### Activation of CB1 and CB2 by eVDs and epoxy-eVDs using a β-arrestin recruitment PRESTO-Tango assay

Previously, we have shown that epoxy-eCBs show an increased activation of CB2 compared to the parent eCBs^57,58^. Therefore, we determined the activation of CB1 and CB2 by eVDs and the epoxy-eVDs using the PRESTO-Tango assay^57^. CP55940 was used as a positive control, which is a full agonist at both CB1 and CB2 (Figure S36a and b). The EC_50_ values for CP55940 were 0.91 ± 0.38 nM and 5.2 ± 1.9 nM at CB1 and CB2, respectively, which agree with previously reported data^57,58,102^.

Herein, NADA, 14’,15’-epoNADA, and 14’,15’-epoNA5HT, but not NA5HT, demonstrated full-agonist activation of CB1 (∼100% activation as compared to CP55940) (Figure 5l and m, Table 1). Of these, NADA activates CB1 2-fold more potently as compared to 14’,15’-epoNADA. 14’,15’-epoNA5HT activated CB1 with an EC_50_ of 16.5 ± 11.7 nM. Only NA5HT activated CB2. The measured EC_50_ was 8.80 ± 5.57 nM with 19.2% activation compared to CP55940, which makes it a partial agonist.

We further tested if NA5HT is an antagonist for CB1 and if NADA, 14’,15’-epoNADA, and 14’,15’-epoNA5HT are antagonists for CB2. None of these antagonized 50 nM CP55940 activation of these receptors (Figure S36c). Therefore, NA5HT does not act on CB1, and NADA, 14’,15’-epoNADA, and 14’,15’-epoNA5HT do not act on CB2. This interesting dichotomy could be exploited to design pain therapeutics that specifically target one receptor. Overall, our data shows that 14’,15’-epoNA5HT is anti-inflammatory, is an agonist of CB1, and is an antagonist of TRPV1, thereby making it the most efficacious of the eVDs for the development of pain therapeutics.

### Inhibition of TRPV1 mediated responses by epoxy-eVDs in primary afferent DRG neurons monitored using live cell Ca^2+^ imaging and whole-cell Patch-clamp recording

To further determine if NA5HT and 14’,15’-epoNA5HT could inhibit native TRPV1 expressed in mouse dorsal root ganglia (DRG) neurons, we compared CAP-evoked intracellular free Ca^2+^ ([Ca^2+^]_i_) response in cultured mouse DRG neurons with and without pretreatment of NA5HT and 14’,15’-epoNA5HT using live cell Ca^2+^ imaging. Bath application of CAP at 250 nM produced a robust [Ca^2+^]_i_ increase in 47.12% ± 2.18% of DRG neurons (Figure 6a & 6d). However, after pretreatment of 1 μM NA5HT or 1 μM 14’,15’-epoNA5HT for 10 mins, 250 nM CAP could only activate 33.64% ± 1.16% and 20.01% ± 2.39% of DRG neurons, respectively (Figure 6b-d). Consistent with the reduced number of DRG neuron activated by CAP after pretreatment of NA5HT and 14’,15’-epoNA5HT, the amplitude of CAP-induced [Ca^2+^]_i_ increase was also significantly diminished after pretreatment of NA5HT and 14’,15’-epoNA5HT (Figure 6b, 6c, 6e). We also examined the inhibitory effect of NA5HT and 14’,15’-epoNA5HT on CAP-induced excitation of DRG neurons by using current-clamp recording. Consistent with Ca^2+^ imaging results, CAP evoked a large membrane potential depolarization and robust action potential firing, which was also significantly inhibited by pretreatment with NA5HT or 14’,15’-epoNA5HT for 10 mins (Figure 6f-i). Of note, inhibition of TRPV1-mediated [Ca^2+^]_i_ increase membrane potential depolarization by 14’,15’-epoNA5HT was significantly stronger than that produced by NA5HT (Figure 6i), which is consistent with results in TRPV1-expressing HEK293 cells. Together, these results suggest that both NA5HT and epoNA5HT are potent TRPV1 antagonists suppressing TRPV1 function in both heterologous cells and native DRG neurons.

**Figure 6.**
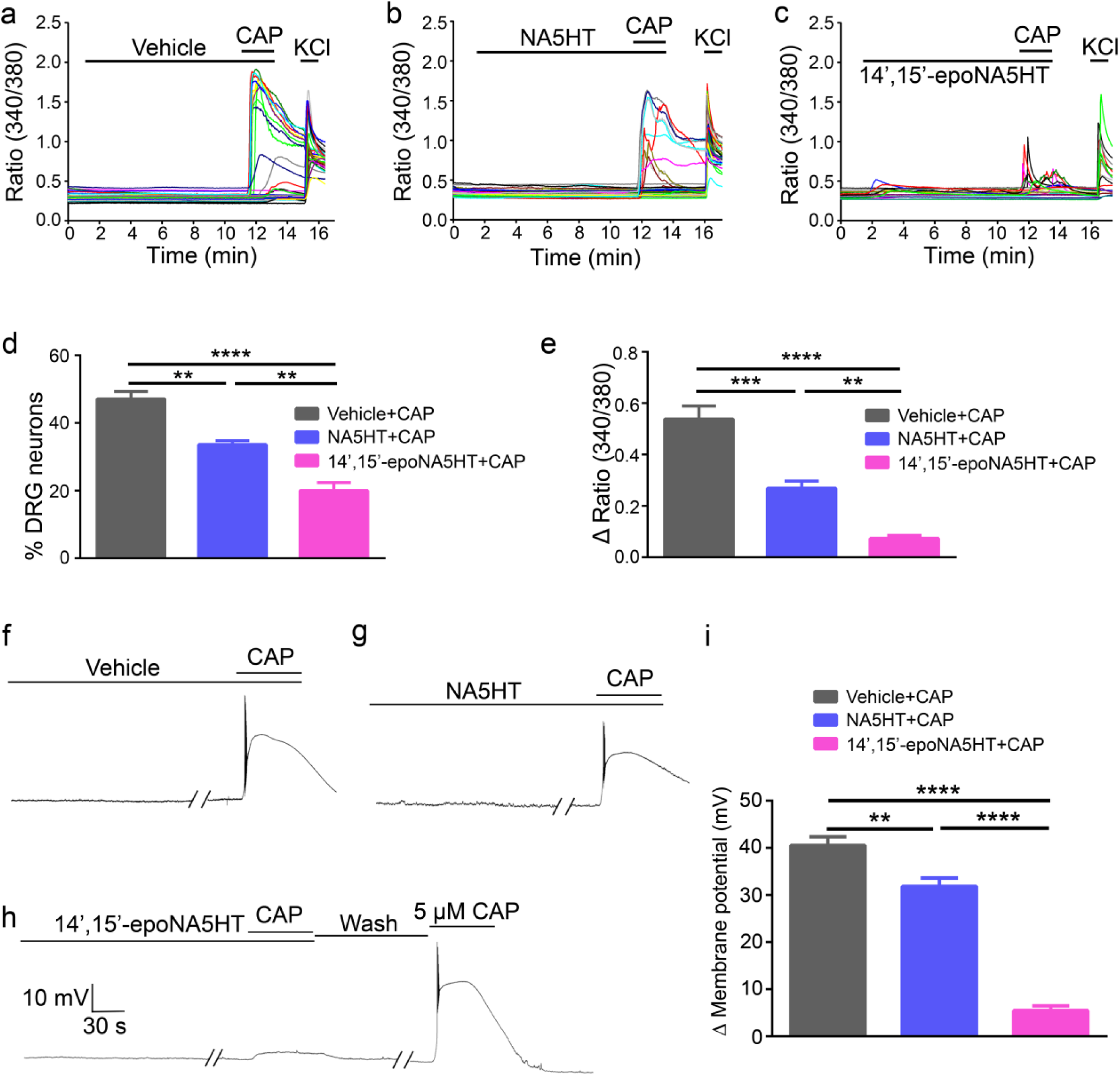
NA5HT and 14’,15’-epoNA5HT inhibit CAP-induced response in cultured mouse DRG neurons. (a-c) Ratiometric Ca^2+^ imaging of cultured wild-type mouse DRG neurons. Each trace corresponds to fluorescence in a single neuron. CAP elicited large [Ca^2+^]_i_ responses in cultured DRG neurons (a), which was significantly inhibited by pretreatment of NA5HT (b) and 14’,15’-epoNA5HT (c) for 10 mins, respectively. (d) Percentage of DRG neurons responding to CAP with or without pretreatment of NA5HT or 14’,15’-epoNA5HT. (*n*=4 to 5). (e) Quantification of CAP-induced [Ca^2+^]_i_ response with or without pretreatment of NA5HT or 14’,15’-epoNA5HT. (*n*=4 to 5). (f) Representative traces showing CAP-induced a robust membrane depolarization and action potential firing in cultured DRG neurons, which was significantly inhibited by pretreatment of NA5HT (g) and 14’,15’-epoNA5HT (h) for 10 mins, respectively. (i) Summarized data showing that CAP-produced membrane depolarization and action potential firing in DRG neurons with or without pretreatment of NA5HT or 14’,15’-epoNA5HT. (*n*=5). Drug concentration: CAP, 250 nM; NA5HT, 1 μM, 14’,15’-epoNA5HT, 1 μM and KCl, 100 mM. **p ≤ 0.01, ***p ≤ 0.001 ****p ≤ 0.0001 (ANOVA).

## DISCUSSION

The endocannabinoid (eCB) system is a promising target to mitigate pain as an alternative to the opioid system. There is also evidence that there is a synergism between the eCB and the opioid system that reduces the need for high doses of opioids^8^. Hence, it is important to understand the function of eCBs and their metabolites as endogenous and exogenous ligands of the receptors that are involved in pain response modulation. It has been previously shown that the activation of CB1 and CB2 is associated with anti-nociceptive and anti-inflammatory actions^12,13^. Control of TRPV1 can also modulate pain by mostly antagonizing TRPV1. While there are ample examples of drugs that target cannabinoid receptors or TRPV1, there is dearth of molecules that are rheostat regulators (varying strength of agonism or antagonism) of both cannabinoid receptors and TRPV1. The primary difference between many drugs and endogenous lipids is that the former are usually structurally rigid and target either CB receptors or TRPV1 whereas the latter are functionally plastic and capable of activating both receptors. NADA and NA5HT are AA derivatives of neurotransmitters—dopamine (DA) and serotonin (5HT). NADA is an agonist of both CB1 and TRPV1^49,50^. NA5HT is an antagonist of TRPV1. PUFAs, through parallel pathways, are converted into epoxide mediators that have been shown to reduce pain, and anti-pain drugs have been developed that prevent the degradation of these molecules^55^. Overall, there is strong evidence for pain modulation by eCB and PUFA epoxides through multiple receptors.

Using a combined biophysical and cell-based approach, we report the pharmacological characterization of NADA and NA5HT epoxides that are produced by the CYP epoxygenase pathway in microglial cells and *in vitro* albeit at low levels. These molecules are anti-inflammatory and function through the eCB-TRPV1 axis. These results can potentially inspire new therapeutics that effectively target this axis. The overall findings of this work are outlined in Figure 7 and discussed below.

**Figure 7.**
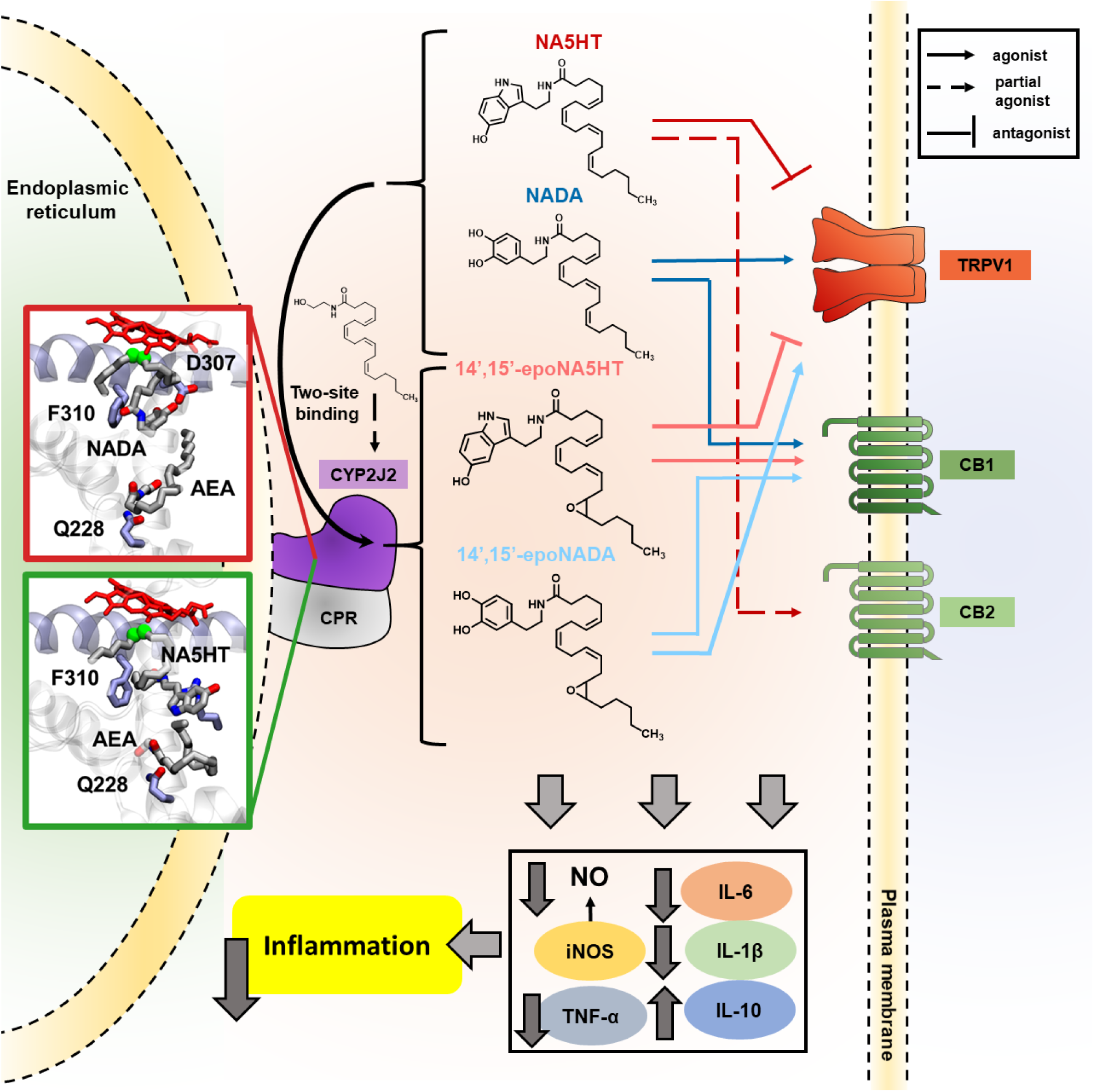
Summary of the eVD metabolism pathway and pharmacology. The eVDs bind in a two-site model to CYP2J2 and other epoxygenases and are metabolized to form epoxy-eVDs. AEA potentiates the metabolism of eVDs as revealed by BV2 metabolism assays, *in vitro* CYP2J2 kinetics, and molecular dynamics simulations. 14’,15’-epoNADA is a better TRPV1 agonist and slightly weaker CB1 agonist compared to NADA. 14’,15’-epoNA5HT is a better TRPV1 antagonist compared to NA5HT, and is a CB1 full agonists as opposed to NA5HT, which is a partial CB2 agonist. Overall, the eVDs and epoxy-eVDs potently downregulate pro-inflammatory NO production and *Il-6, Il-1ß*, and *Tnf-α* expression while increasing *Il-10* expression, thereby demonstrating that they are potently anti-inflammatory.

### Levels of eVDs in porcine brain and metabolism of exogenous eVDs by microglia and CYP2J2-NDs

The biosynthesis of the epoxy-eVDs is facilitated by CYP epoxygenases as the inhibiton of CYP enzymes in microglial cells reduce the levels of these metabolites. In the pig brain, we were able to detect the parent compounds NADA and NA5HT, whose levels were much lower and variable than that of AEA. The epoNADA and epoNA5HT were spontaneously formed by microglia cells in the presence and absence of inflammatory stimulus. As the levels of AEA were high and eVDs were low, we tested the effect of eVD metabolism by CYPs in the presence of AEA. Interestingly, the metabolism of NADA is potentiated by AEA in the microglial cells. This was an interesting observation as there are very few examples where the binding of one ligand at the enzyme active site potentiates the metabolism of another ligand. However, such complicated substrate interactions are common for CYPs with a large active sites such as CYP3A4^84-86^. We demonstrate that AEA is a complicated effector of eVD metabolism in CYP2J2, which potentially explains the observed potentiation of NADA metabolism in BV2 cells. However, AEA may also be inhibiting unknown enzymes that degrade NADA or 14’,15’-epoNADA.

We then proceeded to investigate the the interactions of AEA and the eVDs at the enzymological level to gain better molecular insight into the metabolism. Since reconstituted systems for mouse CYP2J9 and CYP2J12 do not exist, we used a human homologue, CYP2J2, which we have used previously to study CYP2J enzymology. CYP2J9, CYP2J12, and CYP2J2 were all shown to be highly expressed in mouse and human brains^87,88,89-91^ Additionally, CYPs such as CYP2J6 are expressed in DRGs and is involved in the epoxidation of lipids that act on TRPV1 when paclitaxel is administered. In addition, CYP2J4 is found in TRPV1-positive rat trigeminal ganglia, which are also involved in pain-temperature sensing pathways^103^. Comparing the *in cellulo* metabolism data to the *in vitro* kinetic measurements with CYP2J2-NDs reveals a complex network of substrate-substrate interactions. Previously, we determined that PUFAs bind CYP2J2 at two main sites: the substrate access channel and the PUFA binding pocket. Since PUFAs adopt similar conformations, their binding to CYP2J2 is competitive^62^. AEA binds to CYP2J2 with “PUFA-like” properties: that is, by binding similar sites in the substrate access channel and PUFA binding pocket. However, NADA and NA5HT bind in a distinct manner compared to PUFAs and AEA due to their large headgroups. They do not bind the PUFA binding pocket and have unstable binding to the substrate access channel. Therefore, AEA can simultaneously be accommodated in binding to CYP2J2. The MD simulations support that up to three overlapping binding sites are possible, which can help to explain the observed complex AEA and eVD interactions, such as the regioselectivity change in the AEA metabolites (Figure S29) and the potentiation of eVD metabolism (Figures 3 and 4). We have previously observed complex inhibition and regioselectivity changes for endogenous substrates of CYP2J2^59,96,97^; however, this is the first report of a potentiation of CYP2J2 metabolism. Due to the poor solubility of the lipid substrates used in these studies, we could not accurately determine the binding constants to understand the cooperative nature of the potentiation. However, it is evident from the kinetic curves that there is a homotropic potentiation of eVD metabolism and a heterotropic potentiation by AEA. Overall, the finding implies that the co-substrate AEA potentiates the metabolism of NADA and NA5HT by CYP2J2 *in vitro*. Therefore, this study provides another therapeutic route where drugs can potentiate CYP2J2’s epoxidation of endogenous lipids.

### Pharmacological assessment of epoxy-eVDs

There are very few reports on the study of eVD and eCB metabolism by CYPs. One common observation is that the oxidized eVD metabolites exhibit different pharmacology compared to the parent molecules. CYP2U1 metabolizes NA5HT to a 2-oxo derivative that is a weaker FAAH inhibitor compared to NA5HT^47^. NADA was shown to be hydroxylated at the ω and ω-1 positions by rat liver microsomes, which were weaker TRPV1 agonists compared to NADA^49^. Therefore, the 2-oxo-NA5HT and NADA-OH metabolites may represent a degradation pathway of NADA and NA5HT. However, we show that the epoxy-eVDs are potently anti-inflammatory, as they lower IL-6, IL-1β, TNF-α and NO levels while increasing IL-10 levels in activated microglial cells, and therefore may represent an activation pathway.

A key observation is that the epoxidation of eVDs increases their activity on TRPV1. 14’,15’-epoNADA is a 6.5-fold stronger agonist than NADA and 14’,15’-epoNA5HT is a 30-fold stronger antagonist than NA5HT at TRPV1, while also showing partial agonism at lower concentrations in TRPV1-expressing HEK293 cells. Moreover, both NA5HT and 14’,15’-epoNA5HT suppressed TRPV1-mediated [Ca^2+^]_i_ response and membrane potential depolarization in mouse DRG neurons with 14’,15’-epoNA5HT exerting a significantly stronger inhibition of TRPV1-mediated responses than that produced by NA5HT. Based on the cryo-EM structure of TRPV1, it has been proposed that agonists binding at the CAP-binding pocket facilitate channel opening by promoting the lateral movement of the S4-S5 linker^104^. The epoxide could be forming some interactions with H-bond donating groups such as Tyr or Asp residues that populate this linker, which facilitates the channel opening. Since TRPV1 exists as a tetramer, the binding of 14’,15’-epoNA5HT to different sites and perhaps different monomers may help to explain its dual agonism/antagonism. However, since the activation of TRPV1 is self-antagonized at concentrations greater than 250 nM (around the IC_50_), the 14’,15’-epoNA5HT functions overall as an antagonist The epoxidation of NA5HT changes it from being a partial CB2 agonist to a full CB1 agonist. This is intriguing given that 14’,15’-epoNA5HT and NA5HT differ only by an epoxide yet show completely different functions at either CB1 or CB2. Contrariwise, the epoxidation of NADA does not greatly alter their potency towards CB1 receptor activation. Previously it was shown that the epoxidation of omega-6 and omega-3 eCBs have preferential activation towards CB2^57,93^. Herein, we show that eVD epoxides target CB1 receptors. While NADA and 14’,15’-epoNADA are TRPV1 and CB1 agonists, NA5HT and 14’,15’-epoNA5HT are TRPV1 antagonists. The complex functional crosstalk of CB1 and TRPV1 is still being elucidated,^105,106^ but the studies suggest that the plasticity exhibited by endogenous lipids indirectly contributes to this crosstalk.

Overall, the eVDs and epoxy-eVDs lower pro-inflammatory IL-6, IL −1β, TNF-α, and NO levels while increasing anti-inflammatory IL-10. They also potently activate cannabinoid receptors, with affinities between 40 pM and 20 nM, and are potent ligands of TRPV1. The epoxidation of eVDs increases their potency at TRPV1 and alters their pharmacology at cannabinoid receptors. In particular, 14’,15’-epoNA5HT is the most effective epoxy-eVD at reducing pro-inflammatory markers. 14’,15’-epoNA5HT is also a potent antagonist of TRPV1 expressed in either HEK293 cells or native DRG neurons and a potent full agonist of CB1.

Lastly, the formation of epoxy-eVDs by CYPs is potentiated by the co-substrate AEA, which is also metabolized by CYPs to form EET-EA that are potent CB2 ligands. Hence, inflammation and related pain response is accompanied by a storm of epoxy-eVDs and epoxy-eCBs that are multi-faceted endogenous molecules capable of influencing the activity of CB1, CB2 and TRPV1 receptors. The discovery of these molecules will serve as templates for new multi-target therapeutic drugs that will prove useful for the treatment of inflammatory pain as well as of other conditions in which these receptors are targeted in other clinical studies.

## ABBREVIATIONS

AEA: Anandamide
AA: Arachidonic Acid
CB1: cannabinoid receptor 1
CB2: cannabinoid receptor 2
CAP: capsaicin
CYP: cytochrome P450
CPR: cytochrome P450 reductase
DA: dopamine
DRG: dorsal root ganglia
EBS: ebastine
eCB: endocannabinoid
eVD: endovanilloid
EET: epoxyeicosatrienoic acid
epoNADA: epoxyeicosatrienoyl dopamine
EET-EA: epoxyeicosatrienoyl ethanolamide
epoNA5HT: epoxyeicosatrienoyl serotonin
EPOX: epoxygenase
HETE: hydroxyeicosatrienoic acid
HQ: hydroxyquinone
LC-MS/MS: liquid chromatography-tandem mass spectrometry
MD: molecular dynamics
ND: Nanodisc
NADA: N-arachidonoyl-dopamine
NA5HT: N-arachidonoyl-serotonin
PUFA: polyunsaturated fatty acid
5HT: serotonin
TRPV1: transient receptor potential vanilloid 1

## AUTHOR CONTRIBUTIONS

The manuscript was written through contributions of all authors. All authors have given approval to the final version of the manuscript. WRA conceived the project, wrote the manuscript, designed and performed the experiments (synthesis of the compounds, metabolism studies, *in vitro* kinetics, mass spec, receptor activation assays, TRP channel assays, and extractions), made figures, and analyzed the data. LNC designed and performed experiments (synthesis of the compounds, NMR characterization, inflammation assays, metabolism studies, and extractions), wrote the corresponding manuscript sections, prepared figures, and analyzed the data. JLB and ET designed, performed, analyzed, made figures and wrote the sections of the manuscript pertaining to the MD simulations. X.Z and H.H designed, performed, and analyzed the Ca^2+^ imaging and electrophysiology experiments in DRG neurons, prepared figures, and wrote the corresponding section. AD conceived the project, designed the experiments, coordinated the collaborations, and wrote the manuscript.

## ACKNOWLEDGMENTS

We would like to thank Dr. Li Zhong of the Roy J. Carver Metabolomics Center at UIUC for LC-MS/MS analyses and method development. We would also like to thank Dr. Ilia Denisov for helpful discussions regarding the kinetics data. We thank Ms. Hannah Huff for designing the *Cyp2j12* qPCR primers. We thank Ms. Josephine Watson for contributions to the ELISA, NO, MTT, and BrdU assays. We thank Dr. Anuj Yadav and Prof. Jefferson Chan for synthetic assistance to prepare the compounds. We thank Prof. James A. Imlay for thoughtful discussions regarding statistics.

## REFERENCES

1 Hedegaard, H., Warner, M. & Minino, A. M. Drug Overdose Deaths in the United States, 1999-2016. NCHS data brief, 1–8 (2017).

2 Park, K. A. & Vasko, M. R. Lipid mediators of sensitivity in sensory neurons. Trends Pharmacol Sci 26, 571–577 (2005).

3 Cashman, J. N. The mechanisms of action of NSAIDs in analgesia. Drugs 52 Suppl 5, 13–23 (1996).

4 Serhan, C. N. Pro-resolving lipid mediators are leads for resolution physiology. Nature 510, 92–101, doi: 10.1038/nature13479 (2014).

5 Serhan, C. N. & Chiang, N. Resolution phase lipid mediators of inflammation: agonists of resolution. Current opinion in pharmacology 13, 632–640, doi: 10.1016/j.coph.2013.05.012 (2013).

6 Piomelli, D. & Sasso, O. Peripheral gating of pain signals by endogenous lipid mediators. Nature neuroscience 17, 164–174 (2014).

7 Malan, T. P., Jr. & Porreca, F. Lipid mediators regulating pain sensitivity. Prostaglandins & Other Lipid Mediators 77, 123–130 (2005).

8 Gerak, L. R. & France, C. P. Combined Treatment with Morphine and Delta(9)-Tetrahydrocannabinol in Rhesus Monkeys: Antinociceptive Tolerance and Withdrawal. Journal of Pharmacology and Experimental Therapeutics 357, 357–366, doi: 10.1124/jpet.115.231381 (2016).

9 Devane, W. A., Dysarz, F. A., Johnson, M. R., Melvin, L. S. & Howlett, A. C. Determination and characterization of a cannabinoid receptor in rat brain. Molecular Pharmacology 34, 605–613 (1988).

10 Matsuda, L. A., Lolait, S. J., Brownstein, M. J., Young, A. C. & Bonner, T. I. Structure of a cannabinoid receptor and functional expression of the cloned cDNA. Nature 346, 561–564 (1990).

11 Munro, S., Thomas, K. L. & Abu-Shaar, M. Molecular characterization of a peripheral receptor for cannabinoids. Nature 365, 61–65 (1993).

12 Guindon, J. & Hohmann, A. G. The endocannabinoid system and pain. CNS Neurol Disord Drug Targets 8, 403–421 (2009).

13 Di Marzo, V., Bifulco, M. & De Petrocellis, L. The endocannabinoid system and its therapeutic exploitation. Nature reviews. Drug discovery 3, 771–784, doi: 10.1038/nrd1495 (2004).

14 Palazzo, E. et al. Changes in cannabinoid receptor subtype 1 activity and interaction with metabotropic glutamate subtype 5 receptors in the periaqueductal gray-rostral ventromedial medulla pathway in a rodent neuropathic pain model. CNS Neurol Disord Drug Targets 11, 148–161 (2012).

15 Amaya, F. et al. Induction of CB1 cannabinoid receptor by inflammation in primary afferent neurons facilitates antihyperalgesic effect of peripheral CB1 agonist. Pain 124, 175–183 (2006).

16 Croci, T. et al. In vitro functional evidence of neuronal cannabinoid CB1 receptors in human ileum. British Journal of Pharmacology 125, 1393–1395, doi: 10.1038/sj.bjp.0702190 (1998).

17 El-Talatini, M. R. et al. Localisation and Function of the Endocannabinoid System in the Human Ovary. PLoS ONE 4, e4579, doi: 10.1371/journal.pone.0004579 (2009).

18 Fride, E. Endocannabinoids in the central nervous system-an overview. Prostaglandins, Leukotrienes and Essential Fatty Acids 66, 221–233, doi: http://dx.doi.org/10.1054/plef.2001.0360 (2002).

19 Galiègue, S. et al. Expression of Central and Peripheral Cannabinoid Receptors in Human Immune Tissues and Leukocyte Subpopulations. European Journal of Biochemistry 232, 54–61, doi: 10.1111/j.1432-1033.1995.tb20780.x (1995).

20 Lazenka, M. F., Selley, D. E. & Sim-Selley, L. J. Brain regional differences in CB1 receptor adaptation and regulation of transcription. Life Sciences 92, 446–452, doi: http://dx.doi.org/10.1016/j.lfs.2012.08.023 (2013).

21 Fagerberg, L. et al. Analysis of the human tissue-specific expression by genome-wide integration of transcriptomics and antibody-based proteomics. Mol Cell Proteomics 13, 397–406, doi: 10.1074/mcp.M113.035600 [pii] (2014).

22 Manzanares, J., Julian, M. D. & Carrascosa, A. Role of the cannabinoid system in pain control and therapeutic implications for the management of acute and chronic pain episodes. Curr Neuropharmacol 4, 239–257, doi: Doi 10.2174/157015906778019527 (2006).

23 Clayton, N., Marshall, F. H., Bountra, C. & O’Shaughnessy, C. T. CB1 and CB2 cannabinoid receptors are implicated in inflammatory pain. Pain 96, 253–260 (2002).

24 Guindon, J. & Hohmann, A. G. Cannabinoid CB2 receptors: a therapeutic target for the treatment of inflammatory and neuropathic pain. British Journal of Pharmacology 153, 319–334, doi: 10.1038/sj.bjp.0707531 (2008).

25 Costa, B., Giagnoni, G., Franke, C., Trovato, A. E. & Colleoni, M. Vanilloid TRPV1 receptor mediates the antihyperalgesic effect of the nonpsychoactive cannabinoid, cannabidiol, in a rat model of acute inflammation. Br J Pharmacol 143, 247–250, doi: 10.1038/sj.bjp.0705920 (2004).

26 Morales, P., Hurst, D. P. & Reggio, P. H. Molecular Targets of the Phytocannabinoids: A Complex Picture. Prog Chem Org Nat Pr 103, 103–131, doi: 10.1007/978-3-319-45541-9_4 (2017).

27 Bisogno, T. et al. Molecular targets for cannabidiol and its synthetic analogues: effect on vanilloid VR1 receptors and on the cellular uptake and enzymatic hydrolysis of anandamide. Br J Pharmacol 134, 845–852, doi: 10.1038/sj.bjp.0704327 (2001).

28 Caterina, M. J. et al. Impaired nociception and pain sensation in mice lacking the capsaicin receptor. Science 288, 306–313, doi: 8443 [pii] (2000).

29 Tominaga, M. & Caterina, M. J. Thermosensation and pain. J Neurobiol 61, 3–12, doi: 10.1002/neu.20079 (2004).

30 Szallasi, A. & Blumberg, P. M. Vanilloid (Capsaicin) receptors and mechanisms. Pharmacological Reviews 51, 159–212 (1999).

31 Choi, S. I., Yoo, S., Lim, J. Y. & Hwang, S. W. Are sensory TRP channels biological alarms for lipid peroxidation? Int J Mol Sci 15, 16430–16457, doi: 10.3390/ijms150916430 (2014).

32 Jara-Oseguera, A., Simon, S. A. & Rosenbaum, T. TRPV1: on the road to pain relief. Current molecular pharmacology 1, 255–269 (2008).

33 Anand, P. & Bley, K. Topical capsaicin for pain management: therapeutic potential and mechanisms of action of the new high-concentration capsaicin 8% patch. British journal of anaesthesia 107, 490–502, doi: 10.1093/bja/aer260 (2011).

34 Wang, Y. & Wang, D. H. TRPV1 ablation aggravates inflammatory responses and organ damage during endotoxic shock. Clin Vaccine Immunol 20, 1008–1015, doi: 10.1128/CVI.00674-12 [pii] (2013).

35 Tsuji, F. & Aono, H. Role of transient receptor potential vanilloid 1 in inflammation and autoimmune diseases. Pharmaceuticals (Basel) 5, 837–852, doi: 10.3390/ph5080837 [pii] (2012).

36 Yoshida, A. et al. TRPV1 is crucial for proinflammatory STAT3 signaling and thermoregulation-associated pathways in the brain during inflammation. Sci Rep-Uk 6, doi:Artn 26088 10.1038/Srep26088 (2016).

37 Schwartz, E. S. et al. TRPV1 and TRPA1 antagonists prevent the transition of acute to chronic inflammation and pain in chronic pancreatitis. J Neurosci 33, 5603–5611, doi: 10.1523/JNEUROSCI.1806-12.2013 33/13/5603 [pii] (2013).

38 Grabiec, U. & Dehghani, F. N-Arachidonoyl Dopamine: A Novel Endocannabinoid and Endovanilloid with Widespread Physiological and Pharmacological Activities. Cannabis and cannabinoid research 2, 183–196, doi: 10.1089/can.2017.0015 (2017).

39 Devane, W. A. et al. Isolation and structure of a brain constituent that binds to the cannabinoid receptor. Science 258, 1946+ (1992).

40 Mechoulam, R. et al. Identification of an endogenous 2-monoglyceride, present in canine gut, that binds to cannabinoid receptors. Biochemical Pharmacology 50, 83–90, doi: http://dx.doi.org/10.1016/0006-2952(95)00109-D (1995).

41 Sugiura, T. et al. 2-Arachidonoylgylcerol: A Possible Endogenous Cannabinoid Receptor Ligand in Brain. Biochemical and Biophysical Research Communications 215, 89–97, doi: http://dx.doi.org/10.1006/bbrc.1995.2437 (1995).

42 Leishman, E. & Bradshaw, H. B. N-Acyl Amides: Ubiquitous Endogenous Cannabimimetic Lipids That Are in the Right Place at the Right Time. Endocannabinoidome: The World of Endocannabinoids and Related Mediators, 33–48 (2015).

43 Bisogno, T. et al. N-acyl-dopamines: novel synthetic CB(1) cannabinoid-receptor ligands and inhibitors of anandamide inactivation with cannabimimetic activity in vitro and in vivo. Biochem J 351 Pt 3, 817–824 (2000).

44 Bisogno, T. et al. Arachidonoylserotonin and other novel inhibitors of fatty acid amide hydrolase. Biochem Biophys Res Commun 248, 515–522, doi: S0006291X9898874X [pii] (1998).

45 Bezuglov, V. V., Bobrov, M. & Archakov, A. V. Bioactive amides of fatty acids. Biochemistry (Mosc) 63, 22–30 (1998).

46 Huang, S. M. et al. An endogenous capsaicin-like substance with high potency at recombinant and native vanilloid VR1 receptors. Proc Natl Acad Sci U S A 99, 8400–8405, doi: 10.1073/pnas.122196999 99/12/8400 [pii] (2002).

47 Siller, M. et al. Oxidation of endogenous N-arachidonoylserotonin by human cytochrome P450 2U1. Journal of Biological Chemistry 289, 10476–10487, doi: 10.1074/jbc.M114.550004 [pii] (2014).

48 Verhoeckx, K. C. et al. Presence, formation and putative biological activities of N-acyl serotonins, a novel class of fatty-acid derived mediators, in the intestinal tract. Biochim Biophys Acta 1811, 578–586, doi: 10.1016/j.bbalip.2011.07.008 S1388-1981(11)00133-8 [pii (2011).

49 Rimmerman, N. et al. Microsomal omega-hydroxylated metabolites of N-arachidonoyl dopamine are active at recombinant human TRPV1 receptors. Prostaglandins & Other Lipid Mediators 88, 10–17, doi: 10.1016/j.prostaglandins.2008.08.004 (2009).

50 De Petrocellis, L. & Di Marzo, V. N-Acyldopamines and N-Acylserotonins: From Synthetic Pharmacological Tools to Endogenous Multitarget Mediators. Endocannabinoidome: The World of Endocannabinoids and Related Mediators, 67–84 (2015).

51 Maione, S. et al. Analgesic actions of N-arachidonoyl-serotonin, a fatty acid amide hydrolase inhibitor with antagonistic activity at vanilloid TRPV1 receptors. British Journal of Pharmacology 150, 766–781, doi: 10.1038/sj.bjp.0707145 (2007).

52 Yoo, J. M., Sok, D. E. & Kim, M. R. Effect of endocannabinoids on IgE-mediated allergic response in RBL-2H3 cells. Int Immunopharmacol 17, 123–131, doi: 10.1016/j.intimp.2013.05.013 (2013).

53 Green, D. et al. Central activation of TRPV1 and TRPA1 by novel endogenous agonists contributes to mechanical allodynia and thermal hyperalgesia after burn injury. Molecular pain 12, doi: 10.1177/1744806916661725 (2016).

54 Schunck, W. H., Konkel, A., Fischer, R. & Weylandt, K. H. Therapeutic potential of omega-3 fatty acid-derived epoxyeicosanoids in cardiovascular and inflammatory diseases. Pharmacol Ther 183, 177–204, doi: 10.1016/j.pharmthera.2017.10.016 (2018).

55 Zhang, G., Kodani, S. & Hammock, B. D. Stabilized epoxygenated fatty acids regulate inflammation, pain, angiogenesis and cancer. Prog Lipid Res 53, 108–123, doi: 10.1016/j.plipres.2013.11.003 (2014).

56 Zelasko, S., Arnold, W. R. & Das, A. Endocannabinoid metabolism by cytochrome P450 monooxygenases. Prostag Oth Lipid M 116, 112–123, doi: 10.1016/j.prostaglandins.2014.11.002 (2015).

57 McDougle, D. R. et al. Anti-inflammatory omega-3 endocannabinoid epoxides. Proc Natl Acad Sci U S A 114, E6034–E6043, doi: 10.1073/pnas.1610325114 (2017).

58 Roy, J., Watson, J. E., Hong, I., Fan, T. M. & Das, A. Anti-Tumorigenic Properties of Omega-3 Endocannabinoid Epoxides. J Med Chem, doi: 10.1021/acs.jmedchem.8b00243 (2018).

59 Arnold, W. R., Weigle, A. T. & Das, A. Cross-talk of cannabinoid and endocannabinoid metabolism is mediated via human cardiac CYP2J2. J Inorg Biochem 184, 88–99, doi: 10.1016/j.jinorgbio.2018.03.016 (2018).

60 McDougle, D. R., Kambalyal, A., Meling, D. D. & Das, A. Endocannabinoids anandamide and 2-arachidonoylglycerol are substrates for human CYP2J2 epoxygenase. J Pharmacol Exp Ther 351, 616–627, doi: 10.1124/jpet.114.216598 (2014).

61 Trott, O. & Olson, A. J. AutoDock Vina: improving the speed and accuracy of docking with a new scoring function, efficient optimization, and multithreading. J Comput Chem 31, 455–461, doi: 10.1002/jcc.21334 (2010).

62 Arnold, W. R., Baylon, J. L., Tajkhorshid, E. & Das, A. Asymmetric Binding and Metabolism of Polyunsaturated Fatty Acids (PUFAs) by CYP2J2 Epoxygenase. Biochemistry 55, 6969–6980, doi: 10.1021/acs.biochem.6b01037 (2016).

63 McDougle, D. R. et al. Incorporation of charged residues in the CYP2J2 F-G loop disrupts CYP2J2-lipid bilayer interactions. Biochim Biophys Acta 1848, 2460–2470, doi: S0005-2736(15)00237-0 [pii] 10.1016/j.bbamem.2015.07.015 (2015).

64 Phillips, J. C. et al. Scalable molecular dynamics with NAMD. J Comput Chem 26, 1781–1802, doi: 10.1002/jcc.20289 (2005).

65 MacKerell, A. D. et al. All-atom empirical potential for molecular modeling and dynamics studies of proteins. J Phys Chem B 102, 3586–3616, doi: 10.1021/jp973084f (1998).

66 Mackerell, A. D., Jr., Feig, M. & Brooks, C. L., 3rd. Extending the treatment of backbone energetics in protein force fields: limitations of gas-phase quantum mechanics in reproducing protein conformational distributions in molecular dynamics simulations. J Comput Chem 25, 1400–1415, doi: 10.1002/jcc.20065 (2004).

67 Hart, K. et al. Optimization of the CHARMM additive force field for DNA: Improved treatment of the BI/BII conformational equilibrium. J Chem Theory Comput 8, 348–362, doi: 10.1021/ct200723y (2012).

68 Klauda, J. B. et al. Update of the CHARMM all-atom additive force field for lipids: validation on six lipid types. J Phys Chem B 114, 7830–7843, doi: 10.1021/jp101759q (2010).

69 Vanommeslaeghe, K. et al. CHARMM general force field: A force field for drug-like molecules compatible with the CHARMM all-atom additive biological force fields. J Comput Chem 31, 671–690, doi: 10.1002/jcc.21367 (2010).

70 Jorgensen, W. L., Chandrasekhar, J., Madura, J. D., Impey, R. W. & Klein, M. L. Comparison of Simple Potential Functions for Simulating Liquid Water. J Chem Phys 79, 926–935, doi:Doi 10.1063/1.445869 (1983).

71 Feller, S. E., Zhang, Y. H., Pastor, R. W. & Brooks, B. R. Constant-Pressure Molecular-Dynamics Simulation - the Langevin Piston Method. J Chem Phys 103, 4613–4621, doi:Doi 10.1063/1.470648 (1995).

72 Martyna, G. J., Tobias, D. J. & Klein, M. L. Constant-Pressure Molecular-Dynamics Algorithms. J Chem Phys 101, 4177–4189, doi:Doi 10.1063/1.467468 (1994).

73 Darden, T., York, D. & Pedersen, L. Particle Mesh Ewald - an N.Log(N) Method for Ewald Sums in Large Systems. J Chem Phys 98, 10089–10092, doi:Doi 10.1063/1.464397 (1993).

74 Arnold, C. et al. Arachidonic Acid-metabolizing Cytochrome P450 Enzymes Are Targets of ω-3 Fatty Acids. Journal of Biological Chemistry 285, 32720–32733, doi: 10.1074/jbc.M110.118406 (2010).

75 Hu, S. S. et al. The biosynthesis of N-arachidonoyl dopamine (NADA), a putative endocannabinoid and endovanilloid, via conjugation of arachidonic acid with dopamine. Prostaglandins, leukotrienes, and essential fatty acids 81, 291–301, doi: 10.1016/j.plefa.2009.05.026 (2009).

76 Marinelli, S. et al. N-arachidonoyl-dopamine tunes synaptic transmission onto dopaminergic neurons by activating both cannabinoid and vanilloid receptors. Neuropsychopharmacol 32, 298–308, doi: 10.1038/sj.npp.1301118 (2007).

77 Buczynski, M. W. & Parsons, L. H. Quantification of brain endocannabinoid levels: methods, interpretations and pitfalls. Br J Pharmacol 160, 423–442, doi: 10.1111/j.1476-5381.2010.00787.x BPH787 [pii] (2010).

78 Schmid, P. C. et al. Occurrence and postmortem generation of anandamide and other long-chain N-acylethanolamines in mammalian brain. Febs Lett 375, 117–120 (1995).

79 Seki, H., Tani, Y. & Arita, M. Omega-3 PUFA derived anti-inflammatory lipid mediator resolvin E1. Prostaglandins Other Lipid Mediat 89, 126–130, doi: 10.1016/j.prostaglandins.2009.03.002 (2009).

80 Inoue, K. & Tsuda, M. Microglia in neuropathic pain: cellular and molecular mechanisms and therapeutic potential. Nat Rev Neurosci 19, 138–152, doi: 10.1038/nrn.2018.2 (2018).

81 Zhao, H. et al. The role of microglia in the pathobiology of neuropathic pain development: what do we know? Br J Anaesth 118, 504–516 (2017).

82 Snider, N. T., Nast, J. A., Tesmer, L. A. & Hollenberg, P. F. A Cytochrome P450-Derived Epoxygenated Metabolite of Anandamide Is a Potent Cannabinoid Receptor 2-Selective Agonist. Molecular Pharmacology 75, 965–972, doi: 10.1124/mol.108.053439 (2009).

83 Graber, M. N., Alfonso, A. & Gill, D. L. Recovery of Ca2+ pools and growth in Ca2+ pool-depleted cells is mediated by specific epoxyeicosatrienoic acids derived from arachidonic acid. Journal of Biological Chemistry 272, 29546–29553, doi: DOI 10.1074/jbc.272.47.29546 (1997).

84 Atkins, W. M. Non-Michaelis-Menten kinetics in cytochrome P450-catalyzed reactions. Annual Review of Pharmacology and Toxicology 45, 291–310, doi:DOI 10.1146/annurev.pharmtox.45.120403.100004 (2005).

85 Denisov, I. G., Frank, D. J. & Sligar, S. G. Cooperative properties of cytochromes P450. Pharmacol Ther 124, 151–167, doi: 10.1016/j.pharmthera.2009.05.011 S0163-7258(09)00117-X [pii] (2009).

86 Korzekwa, K. R. et al. Evaluation of atypical cytochrome P450 kinetics with two-substrate models: evidence that multiple substrates can simultaneously bind to cytochrome P450 active sites. Biochemistry 37, 4137–4147, doi: 10.1021/bi9715627 (1998).

87 Graves, J. P. et al. Quantitative Polymerase Chain Reaction Analysis of the Mouse Cyp2j Subfamily: Tissue Distribution and Regulation. Drug Metab Dispos 43, 1169–1180, doi: 10.1124/dmd.115.064139 (2015).

88 Graves, J. P. et al. Characterization of the Tissue Distribution of the Mouse Cyp2c Subfamily by Quantitative PCR Analysis. Drug Metab Dispos 45, 807–816, doi: 10.1124/dmd.117.075697 (2017).

89 Ferguson, C. S. & Tyndale, R. F. Cytochrome P450 enzymes in the brain: emerging evidence of biological significance. Trends Pharmacol Sci 32, 708–714, doi: 10.1016/j.tips.2011.08.005 (2011).

90 Nishimura, M., Yaguti, H., Yoshitsugu, H., Naito, S. & Satoh, T. Tissue distribution of mRNA expression of human cytochrome P450 isoforms assessed by high-sensitivity real-time reverse transcription PCR. Yakugaku zasshi : Journal of the Pharmaceutical Society of Japan 123, 369–375 (2003).

91 Dutheil, F. et al. Xenobiotic-Metabolizing Enzymes and Transporters in the Normal Human Brain: Regional and Cellular Mapping as a Basis for Putative Roles in Cerebral Function. Drug Metabolism and Disposition 37, 1528–1538, doi: 10.1124/dmd.109.027011 (2009).

92 Wu, S., Moomaw, C. R., Tomer, K. B., Falck, J. R. & Zeldin, D. C. Molecular cloning and expression of CYP2J2, a human cytochrome P450 arachidonic acid epoxygenase highly expressed in heart. J Biol Chem 271, 3460–3468 (1996).

93 Snider, N. T., Nast, J. A., Tesmer, L. A. & Hollenberg, P. F. A cytochrome P450-derived epoxygenated metabolite of anandamide is a potent cannabinoid receptor 2-selective agonist. Mol Pharmacol 75, 965–972, doi: 10.1124/mol.108.053439 (2009).

94 Napolitano, A., Pezzella, A. & Prota, G. New reaction pathways of dopamine under oxidative stress conditions: nonenzymatic iron-assisted conversion to norepinephrine and the neurotoxins 6-hydroxydopamine and 6, 7-dihydroxytetrahydroisoquinoline. Chem Res Toxicol 12, 1090–1097, doi:tx990079p [pii] (1999).

95 Napolitano, A., Crescenzi, O., Pezzella, A. & Prota, G. Generation of the neurotoxin 6-hydroxydopamine by peroxidase/H2O2 oxidation of dopamine. J Med Chem 38, 917–922 (1995).

96 Carnevale, L., Arango, A., Arnold, W. R., Tajkhorshid, E. & Das, A. Endocannabinoid Virodhamine is an Endogenous Inhibitor of Human Cardiovascular CYP2J2 Epoxygenase. Biochemistry, doi: 10.1021/acs.biochem.8b00691 (2018).

97 Arnold, W. R., Baylon, J. L., Tajkhorshid, E. & Das, A. Arachidonic Acid Metabolism by Human Cardiovascular CYP2J2 Is Modulated by Doxorubicin. Biochemistry 56, 6700–6712, doi: 10.1021/acs.biochem.7b01025 (2017).

98 Lawton, S. K. et al. N-Arachidonoyl Dopamine Modulates Acute Systemic Inflammation via Nonhematopoietic TRPV1. J Immunol 199, 1465–1475, doi: 10.4049/jimmunol.1602151 (2017).

99 Wilhelmsen, K. et al. The endocannabinoid/endovanilloid N-arachidonoyl dopamine (NADA) and synthetic cannabinoid WIN55,212-2 abate the inflammatory activation of human endothelial cells. J Biol Chem 289, 13079–13100, doi: 10.1074/jbc.M113.536953 (2014).

100 Costa, B. et al. The dual fatty acid amide hydrolase/TRPV1 blocker, N-arachidonoyl-serotonin, relieves carrageenan-induced inflammation and hyperalgesia in mice. Pharmacological research 61, 537–546, doi: 10.1016/j.phrs.2010.02.001 (2010).

101 Lam, P. M. et al. Activation of recombinant human TRPV1 receptors expressed in SH-SY5Y human neuroblastoma cells increases [Ca(2+)](i), initiates neurotransmitter release and promotes delayed cell death. Journal of neurochemistry 102, 801–811, doi: 10.1111/j.1471-4159.2007.04569.x (2007).

102 Pertwee, R. G. Pharmacology of cannabinoid receptor ligands. Curr Med Chem 6, 635–664 (1999).

103 Ruparel, S. et al. Plasticity of cytochrome P450 isozyme expression in rat trigeminal ganglia neurons during inflammation. Pain 153, 2031–2039, doi: 10.1016/j.pain.2012.04.027 (2012).

104 Gao, Y., Cao, E., Julius, D. & Cheng, Y. TRPV1 structures in nanodiscs reveal mechanisms of ligand and lipid action. Nature 534, 347–351, doi: 10.1038/nature17964 (2016).

105 Chen, J. et al. Spatial Distribution of the Cannabinoid Type 1 and Capsaicin Receptors May Contribute to the Complexity of Their Crosstalk. Sci Rep-Uk 6, doi:Artn 33307 10.1038/Srep33307 (2016).

106 Musella, A. et al. A novel crosstalk within the endocannabinoid system controls GABA transmission in the striatum. Sci Rep-Uk 7, doi:Artn 7363 10.1038/S41598-017-07519-8 (2017).

