## Supplemental Information for "Anti-inflammatory dopamine- and serotonin-based endocannabinoid epoxides reciprocally regulate cannabinoid receptors and the TRPV1 channel"

[†]Department of Comparative Biosciences, [1]Department of Biochemistry, [2]Department of Anesthesiology, The Center for the Study of Itch & Sensory Disorders, Washington University School of Medicine, [3]Center for Biophysics and Quantitative Biology, [4]Center for Macromolecular Modeling and Bioinformatics, [5]Beckman Institute for Advanced Science and Technology, [6]Department of Bioengineering, Neuroscience program, University of Illinois Urbana-Champaign, Urbana IL 61801

**Corresponding Author**

* To whom correspondence should be addressed:

Aditi Das, Ph.D., University of Illinois Urbana-Champaign, 3836 VMBSB, 2001 South Lincoln Avenue, Urbana IL 61802, Phone: 217-244-0630.

§ These authors have contributd equally to the manuscript

#### Materials and Methods

**Materials.** Human CYP2J2 cDNA was obtained from OriGene (Catalog No. SC321730) and modified as published before[1]. Ampicillin, arabinose, chloramphenicol, isopropyl β-D-1-thiogalactopyranoside (IPTG), and Ni-NTA resin were obtained from Gold Biotechnology. δ-aminolevulinic acid was obtained from Frontier Scientific. NADPH and $NADP^+$ were obtained from P212121.com. 1-palmitoyl-2-oleoyl-sn-glycero-3-phosphocholine (POPC) and 1-hexadecanoyl-2-(9Z-octadecenoyl)-sn-glycero-3-phospho-L serine (POPS) were purchased from Avanti Polar Lipids, Inc. AA, NADA, and NA5HT, SKF 525A (α-phenyl-α-propyl-benzeneacetic acid, 2-(diethylamino)ethyl ester, monohydrochloride), AMG-9810 ((2E)-N-(2,3-dihydro-1,4-benzodioxin-6-yl)-3-[4-(1,1-dimethylethyl)phenyl]-2-propenamide), capsaicin (N-[(4-hydroxy-3-methoxyphenyl)methyl]-6E-8-methyl-nonenamide), and CP55940 (*rac*-5-(1,1-dimethylheptyl)-2-[(1R,2R,5R)-5-hydroxy-2-(3-hydroxypropyl)cyclohexyl]-phenol) were obtained from Cayman Chemical. AA for syntheses was obtained from NuChek. All other materials and reagents used were purchased from Sigma-Aldrich and Fisher Scientific.

**Reverse Transcription quantitative Polymerase Chain Reaction (RT-qPCR).** To measure IL-6, IL-10, IL-1β, TNF-α, CYP2J9, CYP2J12, GAPDH, TRPV1, CB1, and CB2 mRNA, total RNA was isolated from BV2 microglial cells using the Direct-zol RNA Kit, according to the manufacturer's instructions (Zymo Research). One microgram of RNA and random primers were used to synthesize the first strand of cDNA using the High-Capacity cDNA Reverse Transcription Kit (Applied Biosciences). Total cDNA reaction samples were diluted 5-fold in RNAse/DNase-free water and were used as templates for amplification of each gene using the 7500 Real Time PCR System (Applied Biosciences). In addition, corresponding primers purchased from Integrated DNA Technologies (Table S1) and the Power SYBR green PCR Master Mix (Applied Biosciences) were utilized during each qPCR reaction. The mRNA levels of IL-6, IL-10, IL-1β, TNF-α, CYP2J9, CYP2J12, TRPV1, CB1, and CB2 were normalized to those of GAPDH, the internal control, and were analyzed using the $2^{-\Delta\Delta C_T}$ method.[2] Relative mRNA expression was calculated by taking the quotient of the treated sample (LPS + eVD) and LPS alone (IL-6, IL-10, IL-1β, and TNF-α mRNA) or the quotient of LPS-treated and LPS-depleted samples (CYP2J9, CYP2J12, TRPV1, CB1, and CB2). Graphs show the relative mRNA expression of each sample for a given gene. The asterisks above each bar indicate the p-value associated with the standard deviation of each triplicate ΔCT value. P-values were calculated using a two-tailed t-test ($\alpha = 0.05$): 0.0332 (*), 0.0021(**), 0.0002 (***), and < 0.0001 (****).Primers used were designed as shown in Table S1.

**BV2 Anti-inflammatory and Cytotoxicity Assays (Griess assay, IL-6 ELISA, and MTT).** Assays were performed as previously described[3]. BV2 cells were seeded in 24-well plates (200,000 cells/mL) and grown to 80-90% confluency. For eVD screening and dose response studies, media was replaced with serum-free media and the cells were pre-incubated with eVDs for 4 hours prior to stimulation with 25 ng/mL LPS (Sigma-Aldrich, USA). For epoxy-eVDs studies, BV2 were treated with the sEH inhibitor, t-AUCB (1 µM), for 30 min prior to addition of epoxy-eVDs. For inhibition studies, CB1 antagonist, Rimonabant (1 µM), CB2 inverse agonist, AM630 (1 µM), TRPV1 antagonist, AMG 9810 (1 µM), were pre-incubated 30 min before addition of eVDs (1 µM) for 4 hrs. Media was collected after 24 hours and measured for NO levels (Griess assay) and IL-6 levels (ELISA) as previously described[3]. Cellular proliferation/cytotoxicity was assessed by testing the media with a commercially available MTT kit (Cayman Chemical Item No: #10009365).

**BrdU Cell Proliferation Assay**. The BV-2 cells were seeded at 20,000 cells/0.1 mL in a 96-well plate. For eVD screening and dose response studies, media was replaced with serum-free media and the cells were pre-incubated with eVDs for 4 hours prior to stimulation with 25 ng/mL LPS for an additional 24 hours (final well vol. 200 µL). For epoxygenated metabolites, serum-media was replaced with media containing 1 µM of sEH inhibitor (t-AUCB) for 30 min before adding epoVDs and LPS stimulation.

Briefly, after a 21 h incubation, BrdU (1x) was added to the plate for 3 hours. Next, the media was aspirated and the BV-2 cells were fixed according to manufacturer's instruction (Millipore Sigma, Cat. No. 2750). BrdU incorporation was detected by the addition of anti-BrdU monoclonal antibody with Goat anti-mouse IgG peroxidase conjugate. Lastly, TMB substrate solution was added for 30 min at room temperature and stop solution was added. The positive wells were measured using a Biotek Synergy 2 microplate reader (Biotek Instruments, Winooski, VT, USA) at an absorbance wavelength of 450 nm.

**Expression and purification of recombinant CYP2J2 in *E. coli*.** Recombinant D34-CYP2J2 containing a $His_5$ tag was expressed and purified as previously performed[1,4]. The D34-CYP2J2 is a 34-residue N-terminal truncation (residues 3-37) of CYP2J2 with a substitution of Leu2 for an Ala residue. These modifications have been previously shown to increase protein yield without affecting activity[38, 45].

**Expression and purification of cytochrome P450 reductase.** Expression of cytochrome P450 reductase (CPR) was performed as described previously[1].

**Use of reactive oxygen species (ROS) scavengers.** All *in vitro* metabolism data using recombinant proteins CYP2J2-CPR in nanodiscs were performed using ROS scavengers as previously described[5].

**CYP2J2-mediated metabolism of NADA and NA5HT.** Metabolism of NADA and NA5HT was performed in a lipid reconstituted system as previously described[5,6]. Briefly, 0.6 μM CYP2J2 and 0.6 μM CPR were incubated in a 20% POPS and 80% POPC reconstituted system in 0.5 mL of 0.1 M potassium phosphate buffer, pH 7.4 at 37$^{o}$ C with 100 μM NADA or NA5HT for 10 min. Reactions were initiated with 1 mM NADPH (final) and reacted for 60 min. Reactions were terminated by vortexing with 0.5 mL ethyl acetate. Adequate amount of $NaCl_{(s)}$ was added to facilitate layer separation. Metabolites were extracted thrice by vortexing the reactions for 1 min with 0.5 mL ethyl acetate followed by centrifugation at 3K RPM at 4$^{o}$ C for 5 minutes to separate the layers. The organic layers were dried under a stream of $N_{2(g)}$ and resuspended in 0.1 mL of acetonitrile for HPLC and LC-MS/MS analysis. For all other experiments, CYP2J2 was incorporated into Nanodiscs.

**Incorporation of CYP2J2 into Nanodiscs.** Nanodiscs (NDs) containing CYP2J2 were prepared in 20% POPS and 80% POPC Nanodiscs as previously described[7]. NDs were used for all experiments using *in vitro* CYP2J2 except for initial product determination.

**Kinetics of NADA and NA5HT metabolism.** The kinetics of NADA and NA5HT metabolism was determined using a CYP2J2- ND/CPR system as previous described[7] with the following modifications. Reactions were performed with ROS scavengers as stated above. NADA and NA5HT (10-100 μM in DMSO) were incubated with CYP2J2-ND/CPR in 0.5 mL of 0.1 M potassium phosphate buffer (pH 7.4) for 10 min. Reactions were initiated with the addition of 0.5 mM NADPH. The formation of epoxy-eVDs was determined to be linear up to 45 min, and so a 30-min reaction was utilized. Reactions were terminated upon the addition of 0.5 mL ethyl acetate and approximately 10 mg $NaCl_{(s)}$ to facilitate layer separation. The products were extract thrice with ethyl acetate, dried under a stream of $N_{2(g)}$, and resuspended in 180-proof ethanol for LC-MS/MS quantification.

**NADPH kinetics.** The rate of NADPH oxidation by CYP2J2-ND/CPR was determined using UV-Vis spectroscopy as previously described[8]. The rate of the NADPH oxidation with CYP2J2-ND/CPR without substrates was considered the "baseline" rate.

**AEA metabolism inhibition.** AEA metabolism was determined as previously described[6] with the following modifications. Reactions were performed in the presence of ROS scavengers as stated above and were terminated using 0.5 mL ethyl acetate and $NaCl_{(s)}$. 25 μM and 75 μM of NADA or NA5HT were used to inhibit AEA metabolism. 40 μM of AEA without inhibitor was used as a control control to compare against our previous measurements for AEA metabolism[6]. The levels of AEA metabolism were similar to previously published and were used to assess saturation level ($v/V_{max}$)[6]. EET-EAs and epoxy-eVDs were simultaneously quantified using LC-MS/MS.

**Ebastine competitive binding**. Relative binding affinities of eVDs binding to CYP2J2 were determined using an ebastine competitive binding assay as previously described.[7]

**LC-MS and LC-MS/MS analysis of CYP2J2 metabolites.** CYP2J2 metabolism products were determined using HPLC Method 1. The Phenomenex column was used. LC-MS/MS detection was performed as previously described[6].

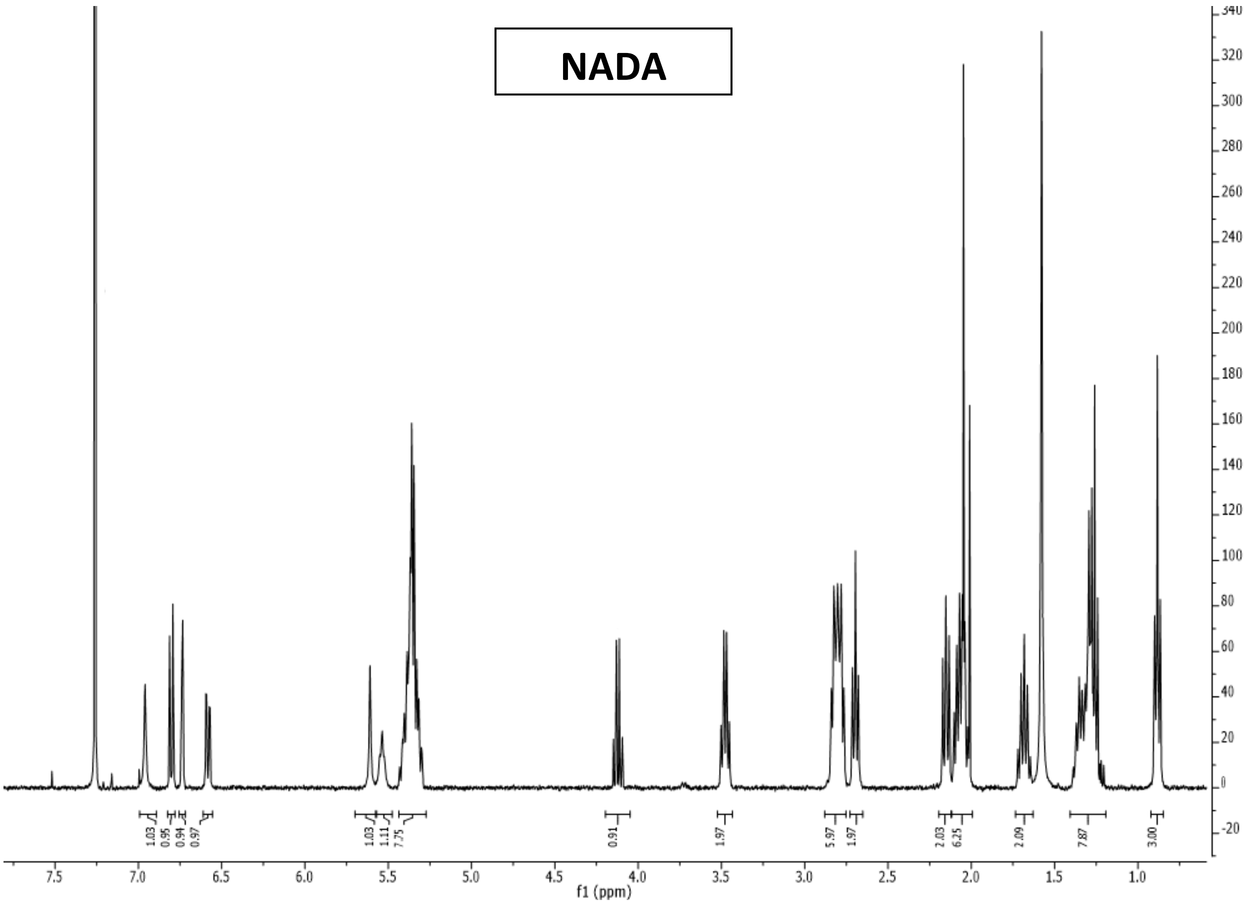

**Figure S1**. $^{1}$H-NMR spectrum for NADA (400 Hz, $CDCl_3$). Peak at 4.12 ppm corresponds to residual ethyl acetate. Other peaks for ethyl acetate overlap with NADA peaks (ethyl acetate: 2.05 and 1.26 ppm in $CDCl_3$)

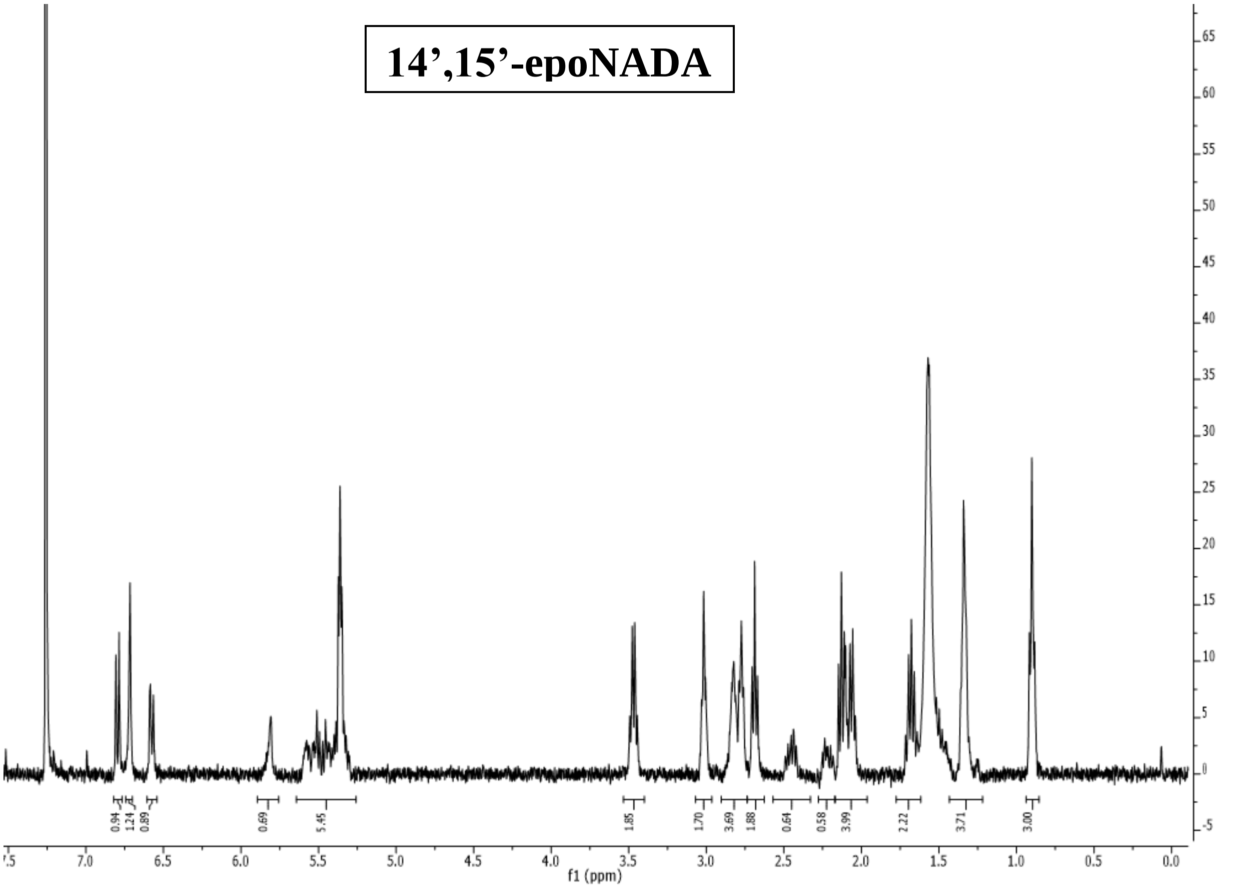

**Figure S2**. [1]H-NMR spectrum for 14’,15’-epoNADA (400 Hz, $CDCl_3$).

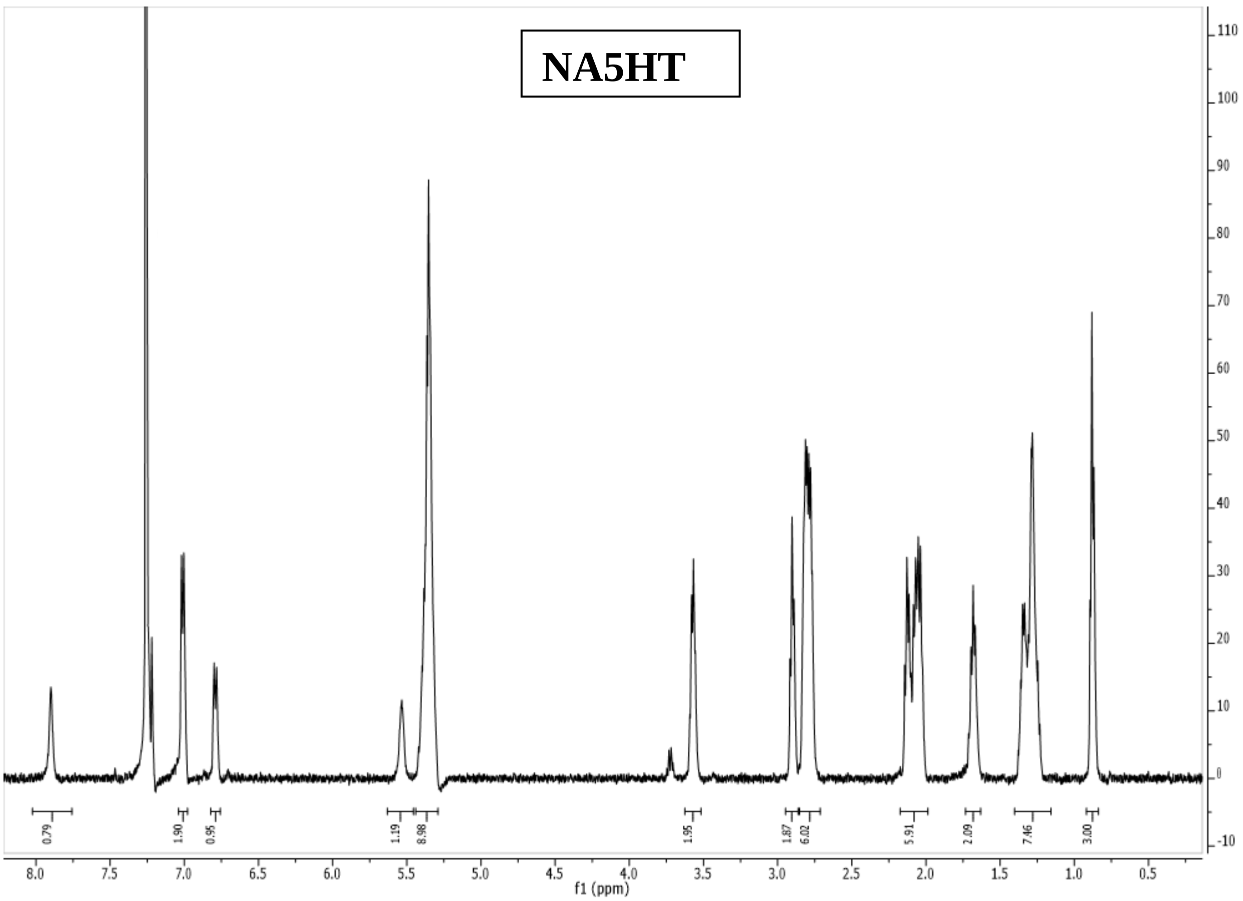

**Figure S3**. $^{1}$H-NMR spectrum for NA5HT (400 Hz, $CDCl_3$).

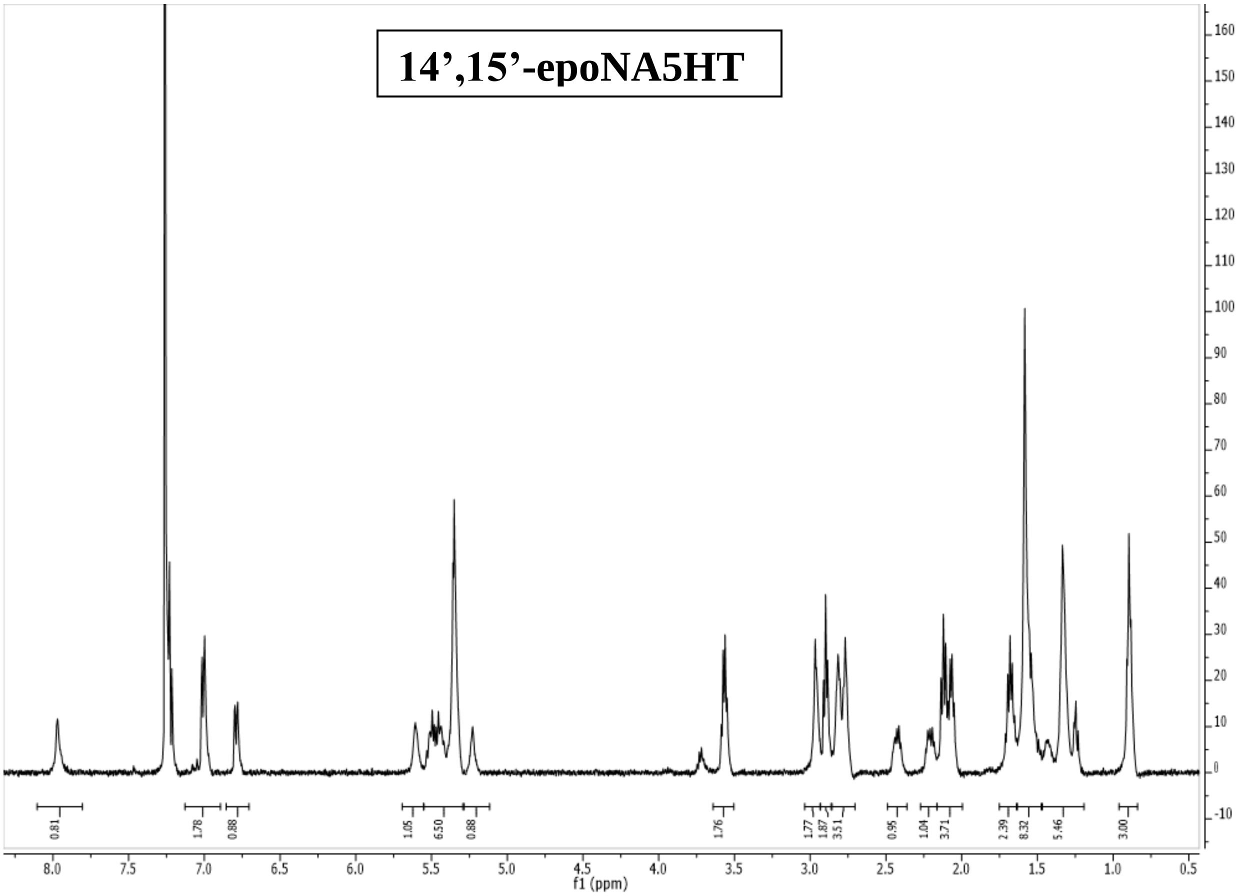

**Figure S4**. [1]H-NMR spectrum for 14’,15’-epoNA5HT (500 Hz, $CDCl_3$).

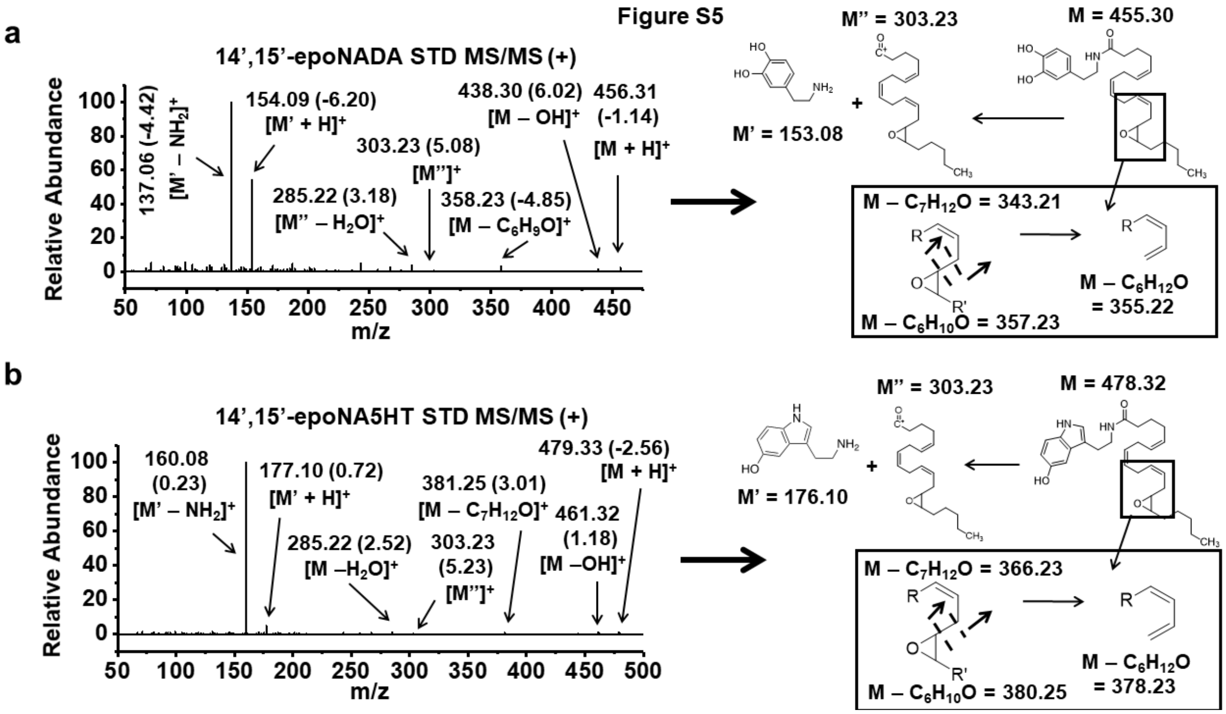

**Figure S5.** MS/MS spectra of synthesized **(a)** 14’,15’-epoNADA and **(b)** 14’,15’-epoNA5HT standards along with fragmentation schemes. Data are obtained from LC-MS/MS analysis. Fragment m/z values are given with deviation from the calculated m/z values (± ppm) given in parentheses.

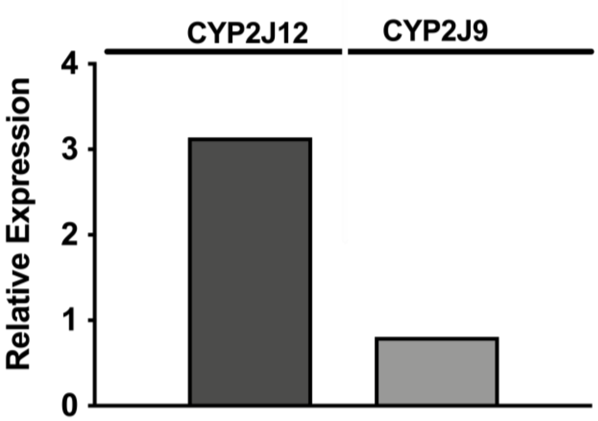

**Figure S6.** *Effect of LPS in BV2 microglial cells on the relative mRNA expression of CYP2J12 and CYP2J9.* BV2 microglial cells seeded at 200,000 cells/well in a 24-well plate were treated with 100 ng/mL LPS for 3 hours. RT-qPCR detected the presence of CYP2J12 and CYP2J9 in BV2 cells. Data is from 3-pooled wells and is performed in technical replicates. Data is reported as the mean relative expression. Statistical analysis was performed using a two-tailed t-test ($\alpha = 0.05$). P-values: 0.0332 (*), 0.0021(**), 0.0002 (***), and < 0.0001 (****).

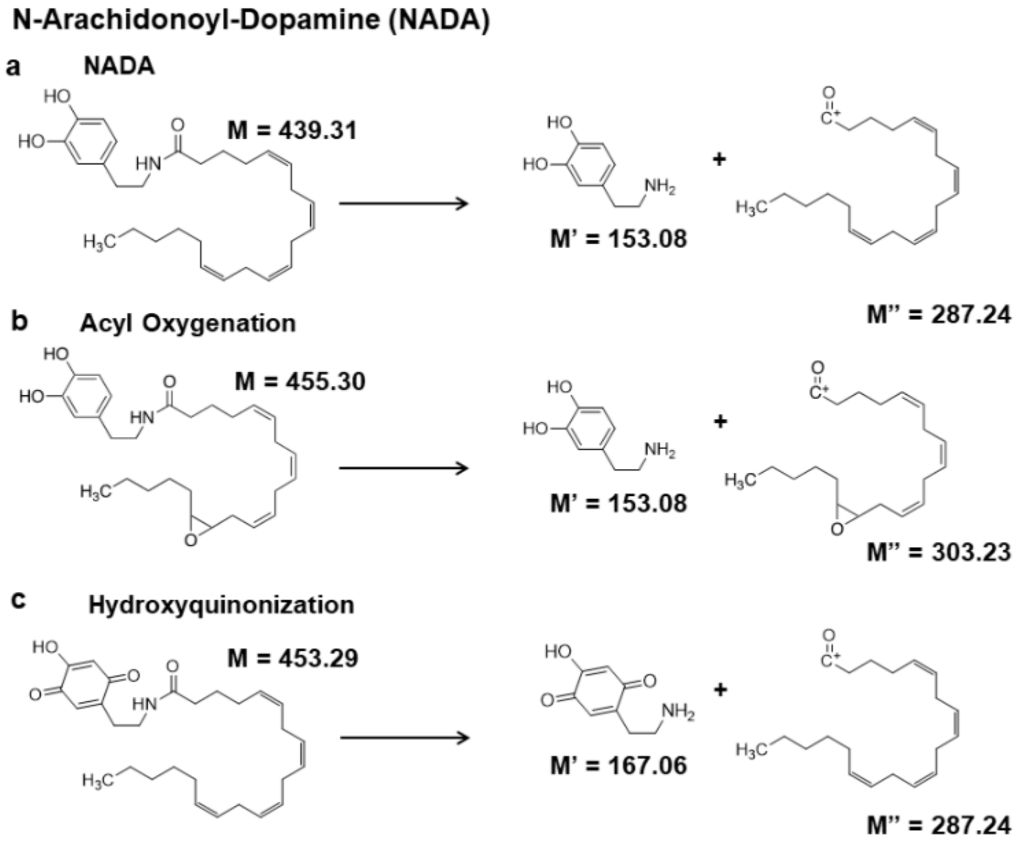

**Figure S7.** Fragmentation scheme of **(a)** NADA, **(b)** acyl oxygenation of NADA (14',15'-epoNADA shown as an example), and **(c)** NADA-HQ.

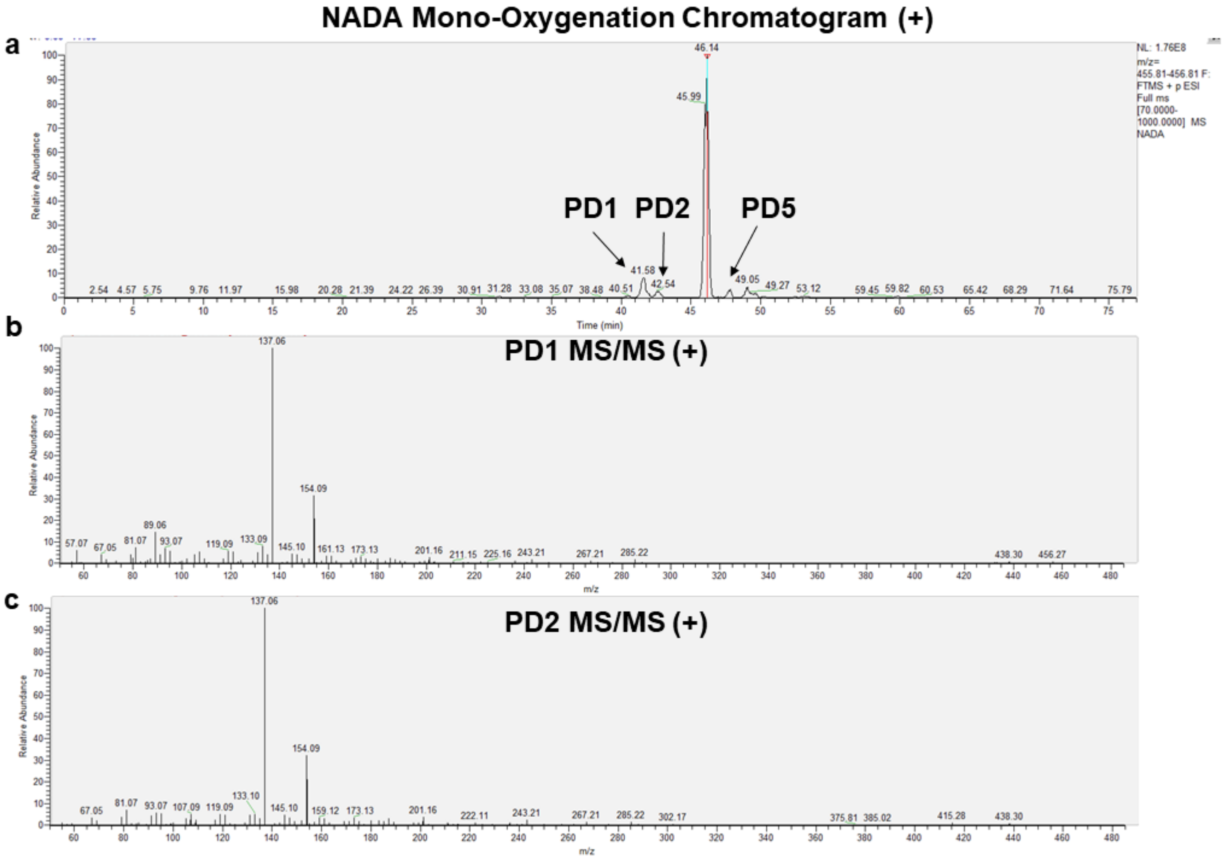

**Figure S8.** *Mono-oxygenation of NADA by CYP2J2*. **(a)** LC-MS/MS chromatogram in positive ion mode. Mass range is ± 5 ppm of the predicted product. **(b-c)** MS/MS spectra obtained from each corresponding product.

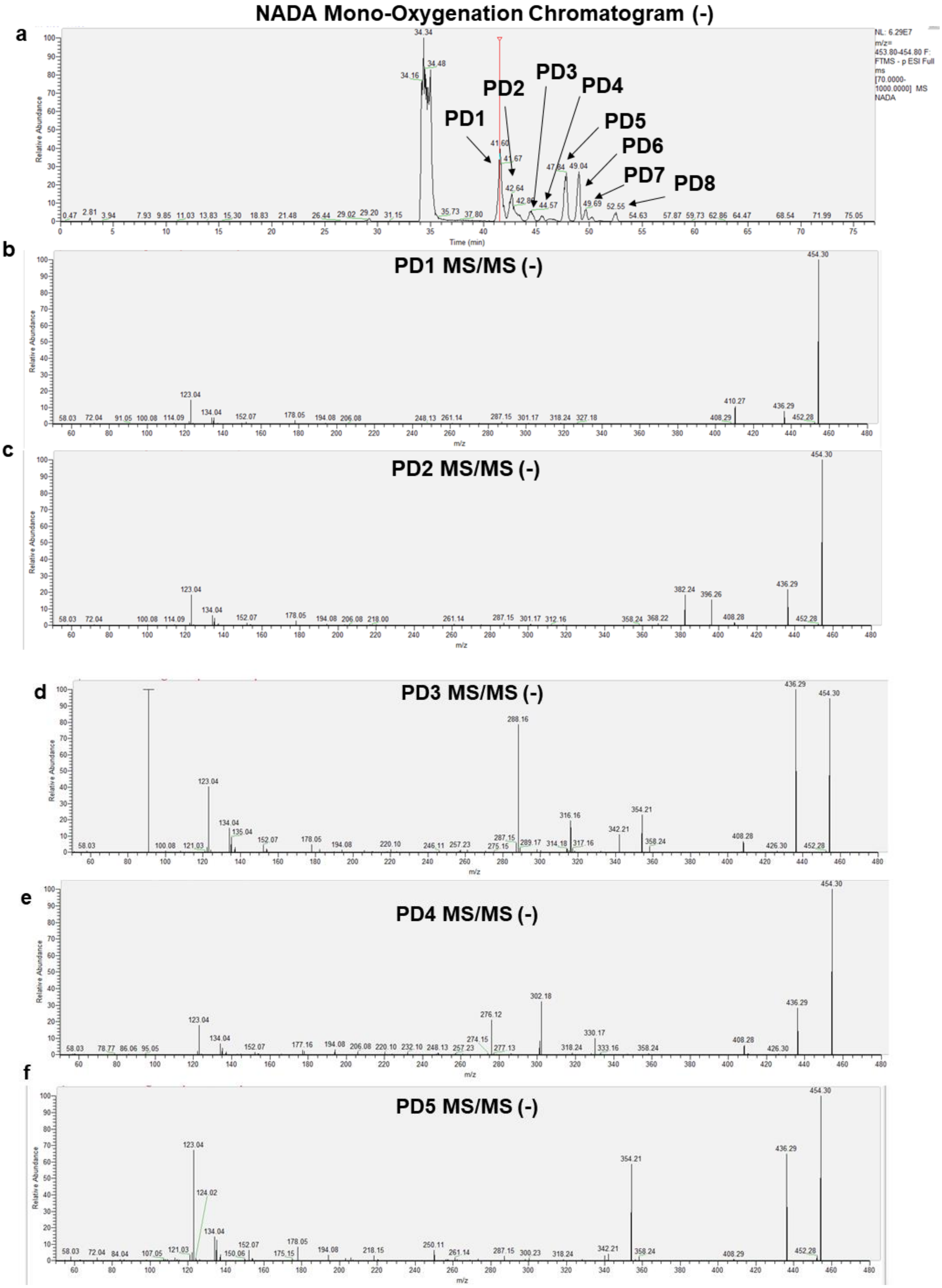
NADA Mono-Oxygenation Chromatogram (-)
a
PD1
PD2
PD3
PD4
PD5
PD6
PD7
PD8
Relative Abundance
Time (min)
b
PD1 MS/MS (-)
c
PD2 MS/MS (-)
d
PD3 MS/MS (-)
e
PD4 MS/MS (-)
f
PD5 MS/MS (-)
m/z

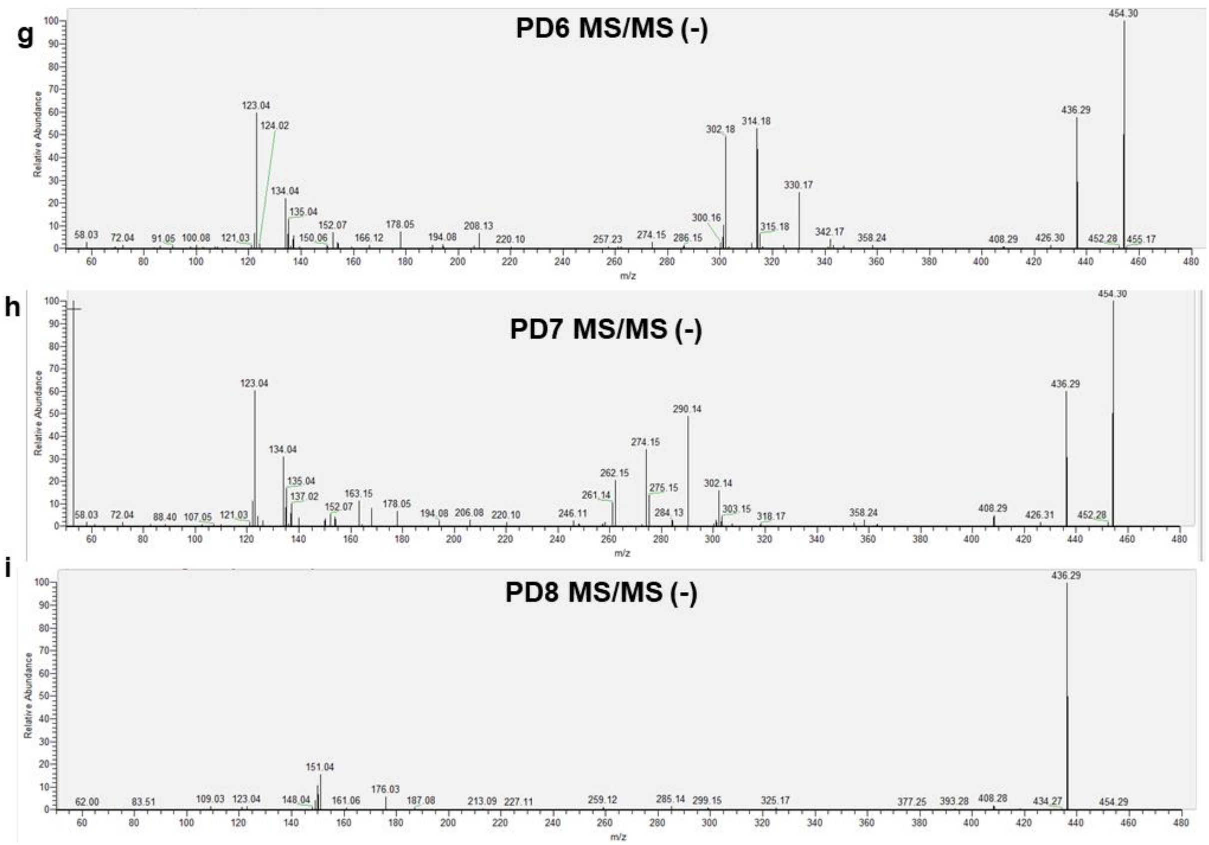

**Figure S9.** *Mono-oxygenation of NADA by CYP2J2*. **(a)** LC-MS/MS chromatogram in negative ion mode. Mass range is ± 5 ppm of the predicted product. **(b-i)** MS/MS spectra obtained from each corresponding product.

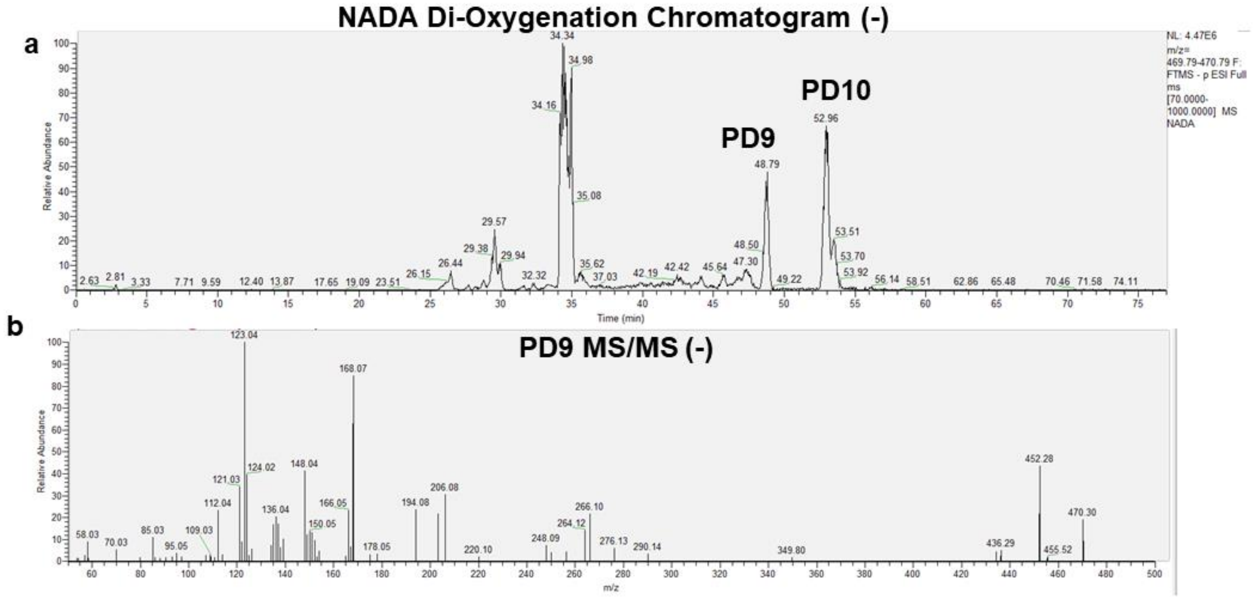

**Figure S10.** *Di-oxygenation of NADA by CYP2J2*. **(a)** LC-MS/MS chromatogram in negative ion mode. Mass range is ± 5 ppm of the predicted product. **(b)** MS/MS spectrum obtained from PD9.

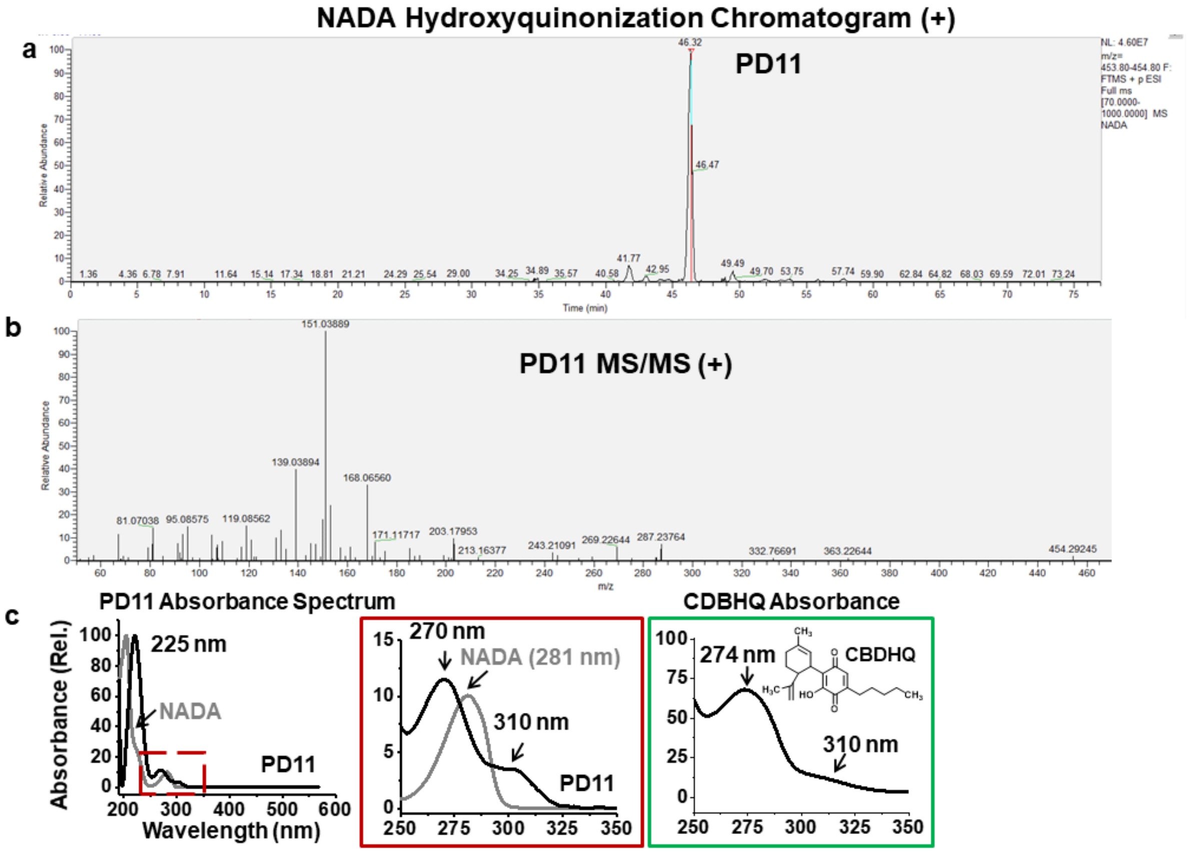

**Figure S11.** *Hydroxyquinonization of NADA by CYP2J2*. **(a)** LC-MS/MS chromatogram in positive ion mode. Mass range is ± 5 ppm of the predicted product. **(b)** MS/MS spectrum of PD11. **(c)** Absorbance spectrum of PD11 obtained by UV-Vis HPLC analysis. NADA and cannabidiol hydroxyquinone (CBDHQ) were analyzed using a similar method and their absorbance spectra are given for comparison.

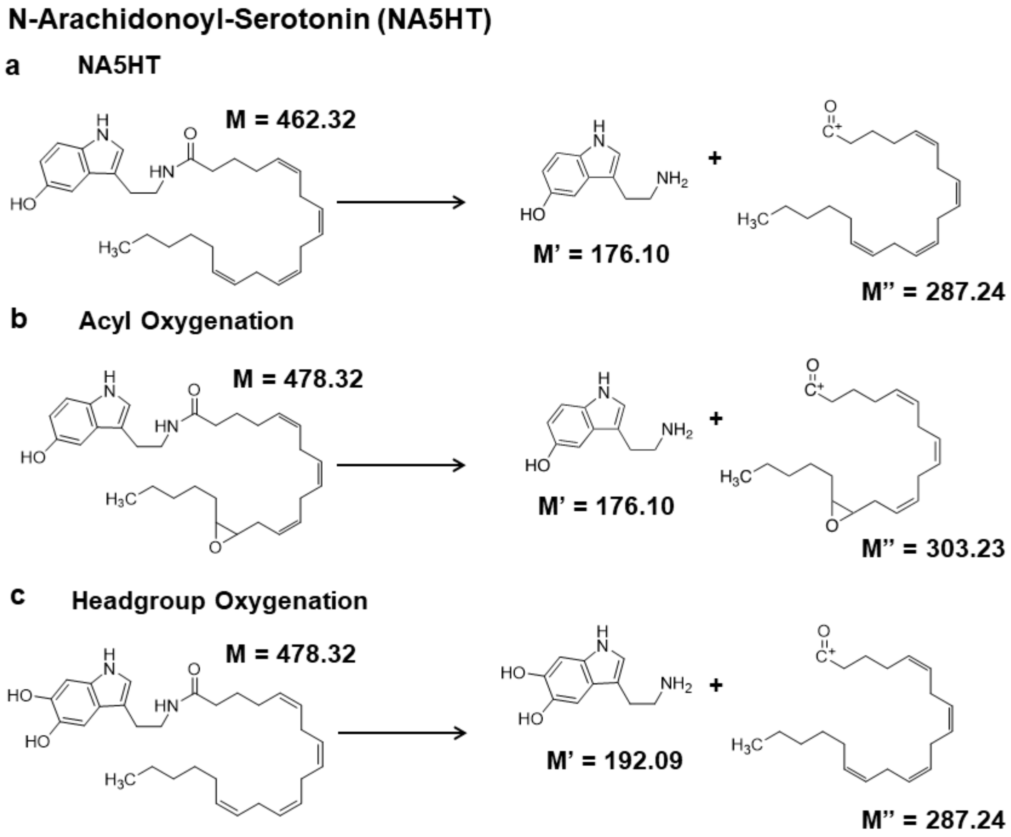

**Figure S12.** Fragmentation scheme of **(a)** NA5HT, **(b)** acyl oxygenation of NA5HT (14',15'-epoNA5HT shown as an example), and **(c)** Headgroup oxygenation of NA5HT.

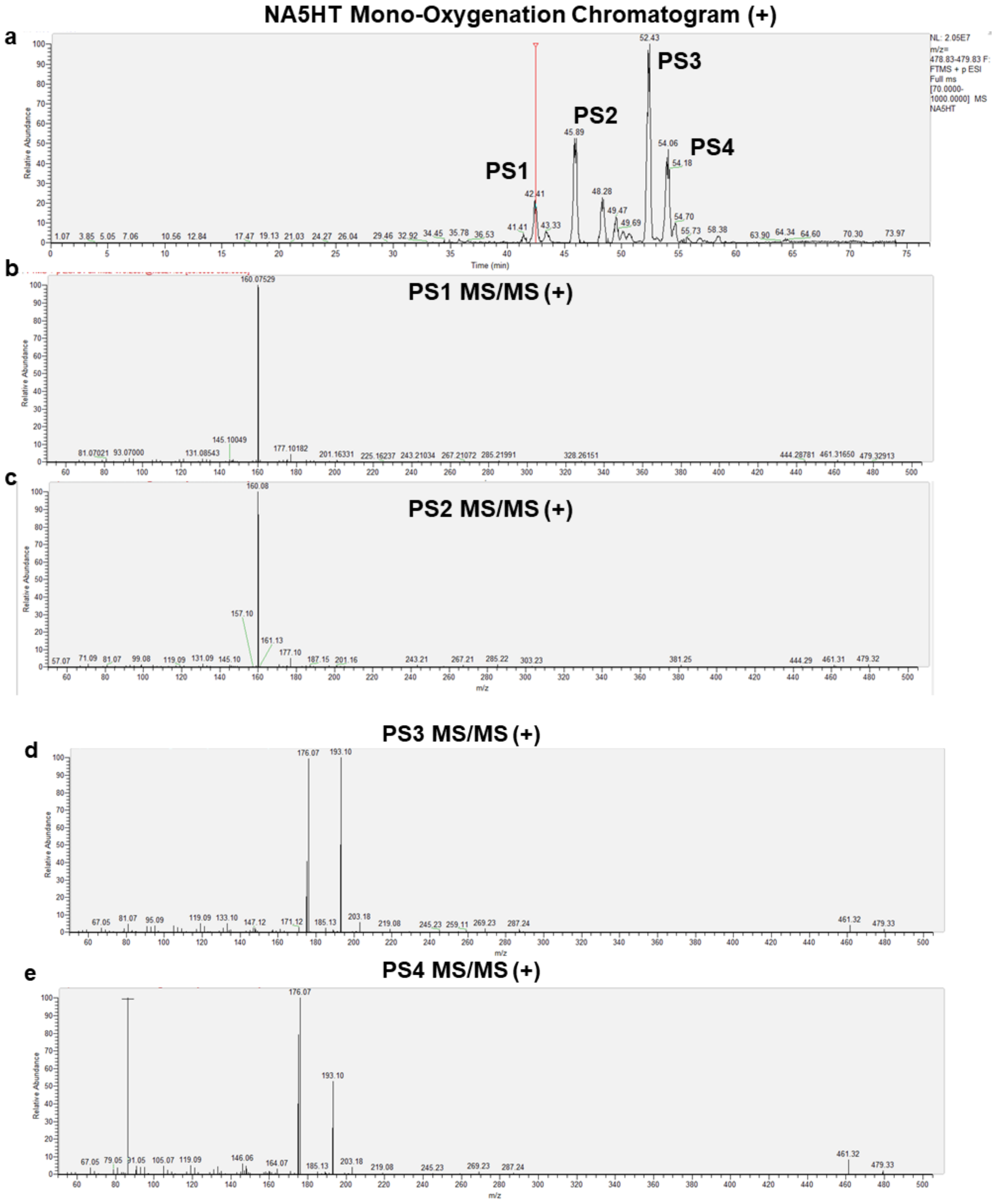

**Figure S13.** *Mono-oxygenation of NA5HT by CYP2J2*. **(a)** LC-MS/MS chromatogram in positive ion mode. Mass range is ± 5 ppm of the predicted product. **(b-e)** MS/MS spectra obtained from each corresponding product.

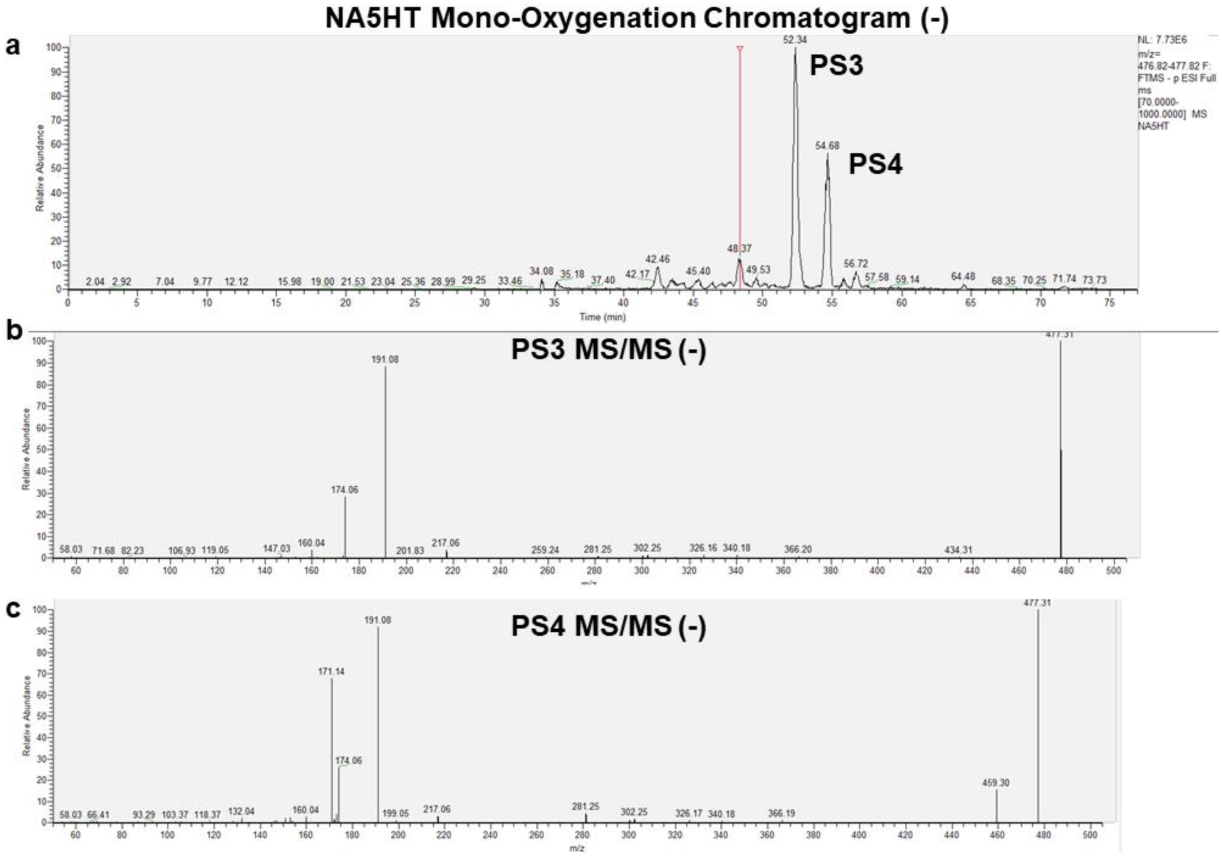

**Figure S14.** *Mono-oxygenation of NA5HT by CYP2J2*. **(a)** LC-MS/MS chromatogram in negative ion mode. Mass range is ± 5 ppm of the predicted product. **(b-c)** MS/MS spectra obtained from each corresponding product.

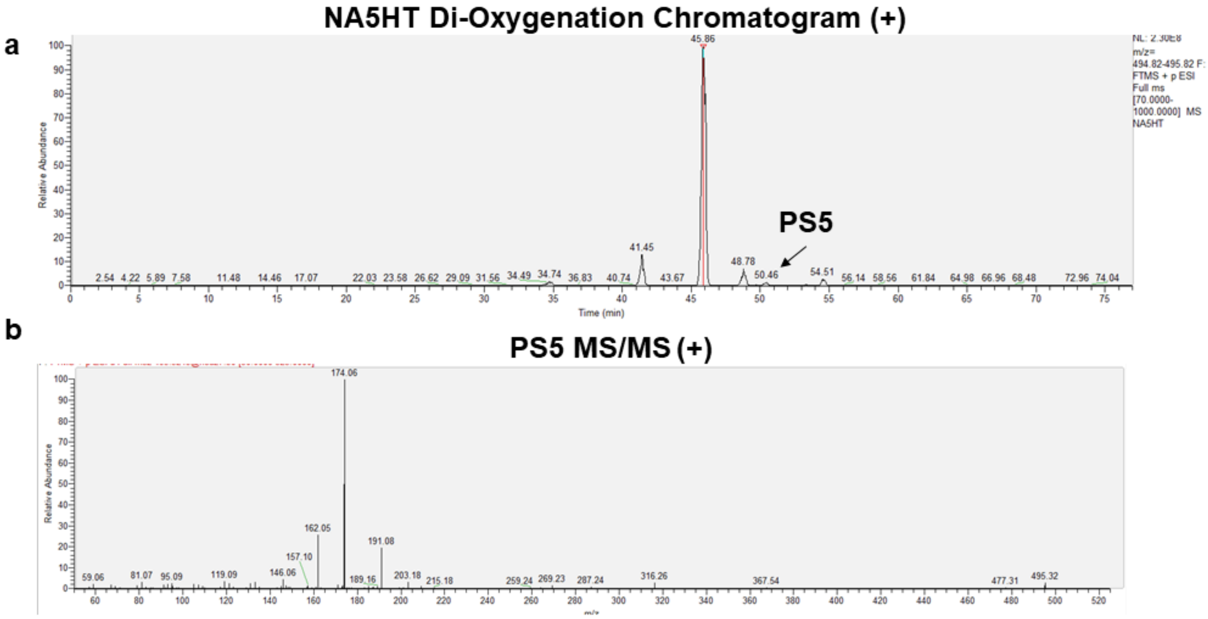

**Figure S15.** *Di-oxygenation of NA5HT by CYP2J2*. **(a)** LC-MS/MS chromatogram in positive ion mode. Mass range is ± 5 ppm of the predicted product. **(b)** MS/MS spectrum of PS5.

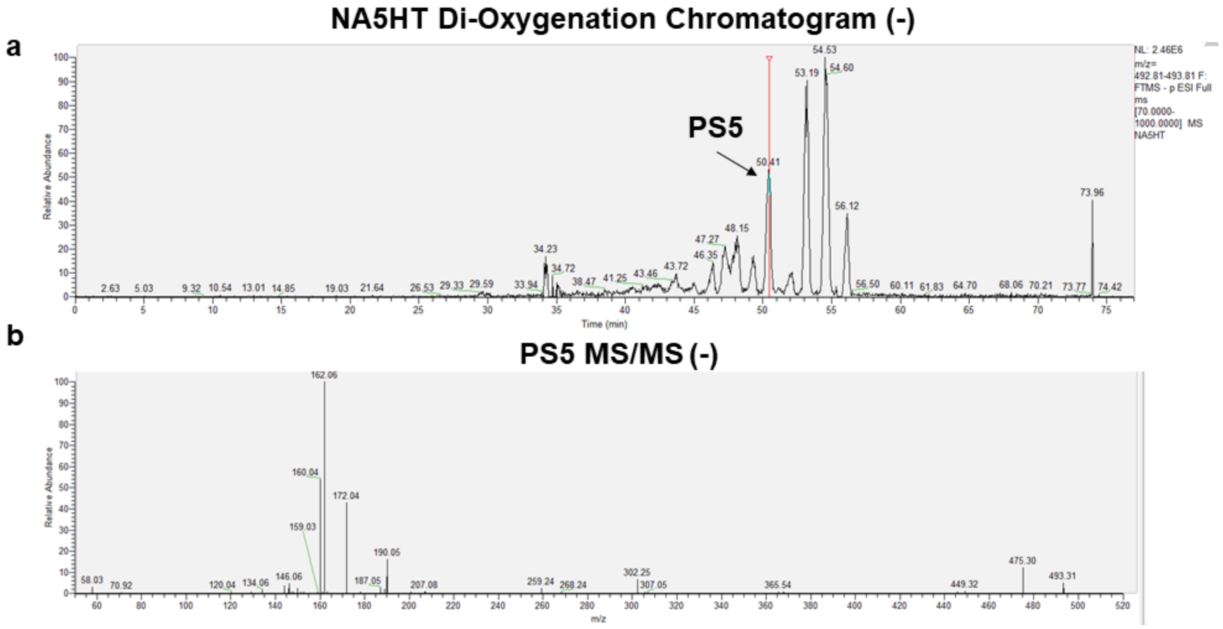

**Figure S16.** *Di-oxygenation of NA5HT by CYP2J2*. **(a)** LC-MS/MS chromatogram in negative ion mode. Mass range is ± 5 ppm of the predicted product. **(b)** MS/MS spectrum of PS5.

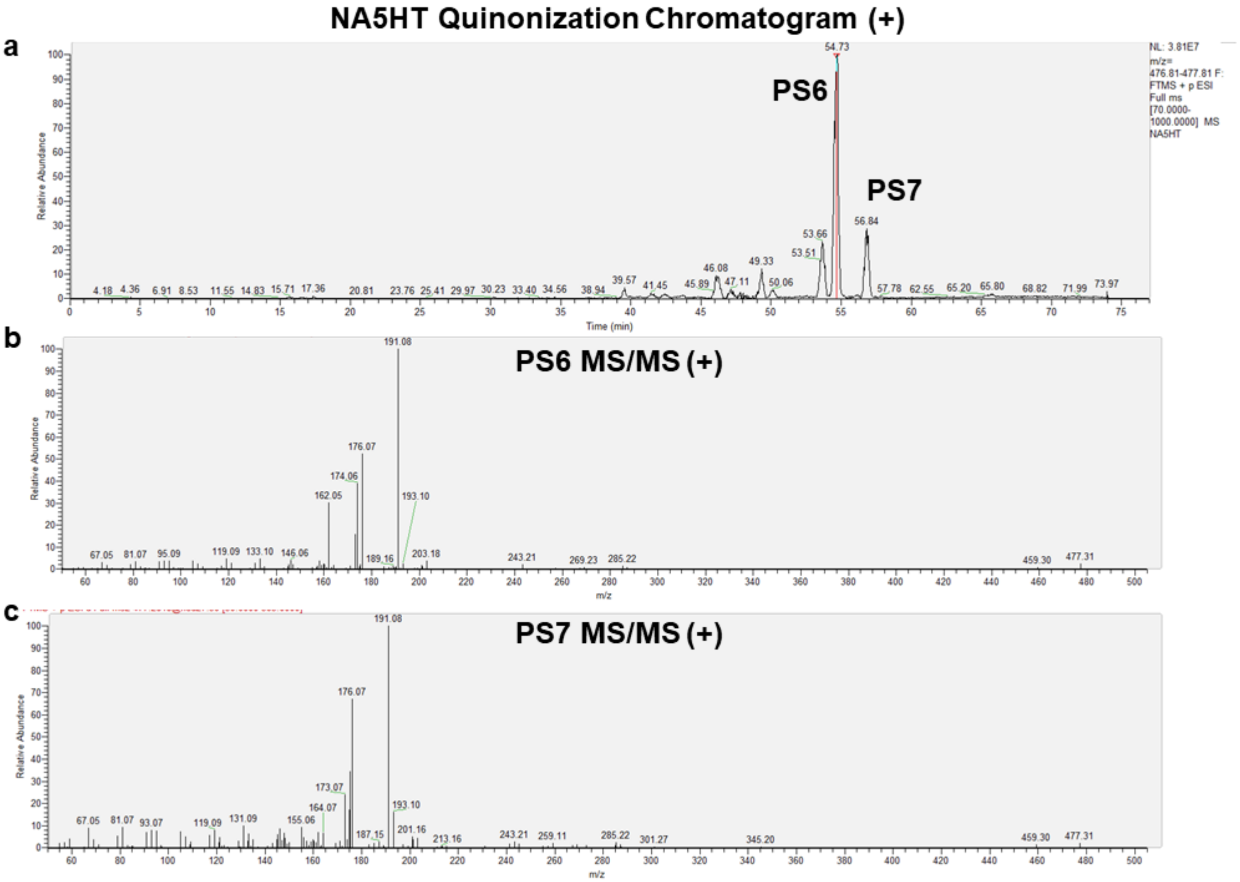

**Figure S17.** *Quinonization of NA5HT by CYP2J2*. Quinonization was determined from the apparent oxygenation of the mono-oxygenation products (PS3 and PS4) indicated by a loss of 2 hydrogen atoms. **(a)** LC-MS/MS chromatogram in positive ion mode. Mass range is ± 5 ppm of the predicted product. **(b-c)** MS/MS spectra obtained from the indicated product.

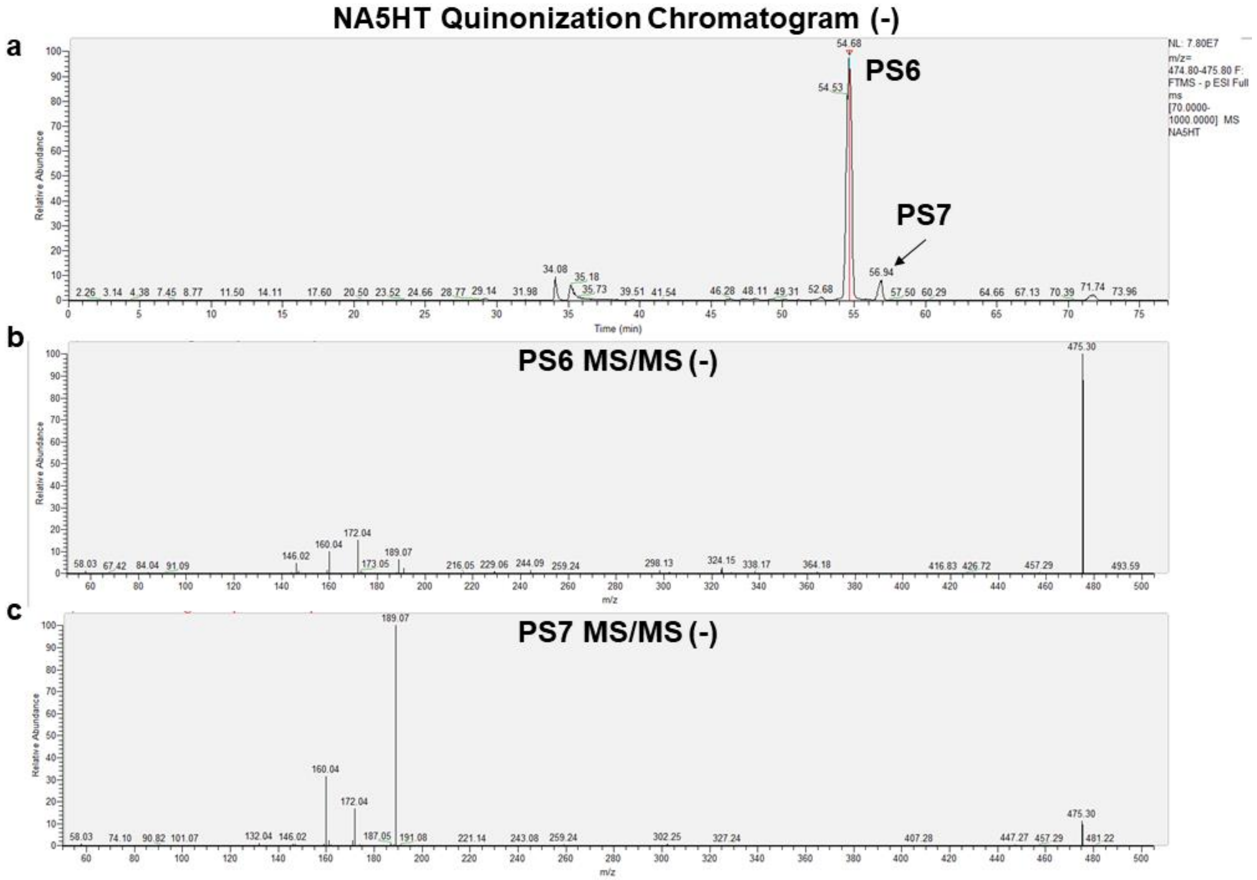

**Figure S18.** *Quinonization of NA5HT by CYP2J2*. Quinonization was determined from the apparent oxygenation of the mono-oxygenation products (PS3 and PS4) indicated by a loss of 2 hydrogen atoms. **(a)** LC-MS/MS chromatogram in negative ion mode. Mass range is ± 5 ppm of the predicted product. **(b-c)** MS/MS spectra obtained from the indicated product.

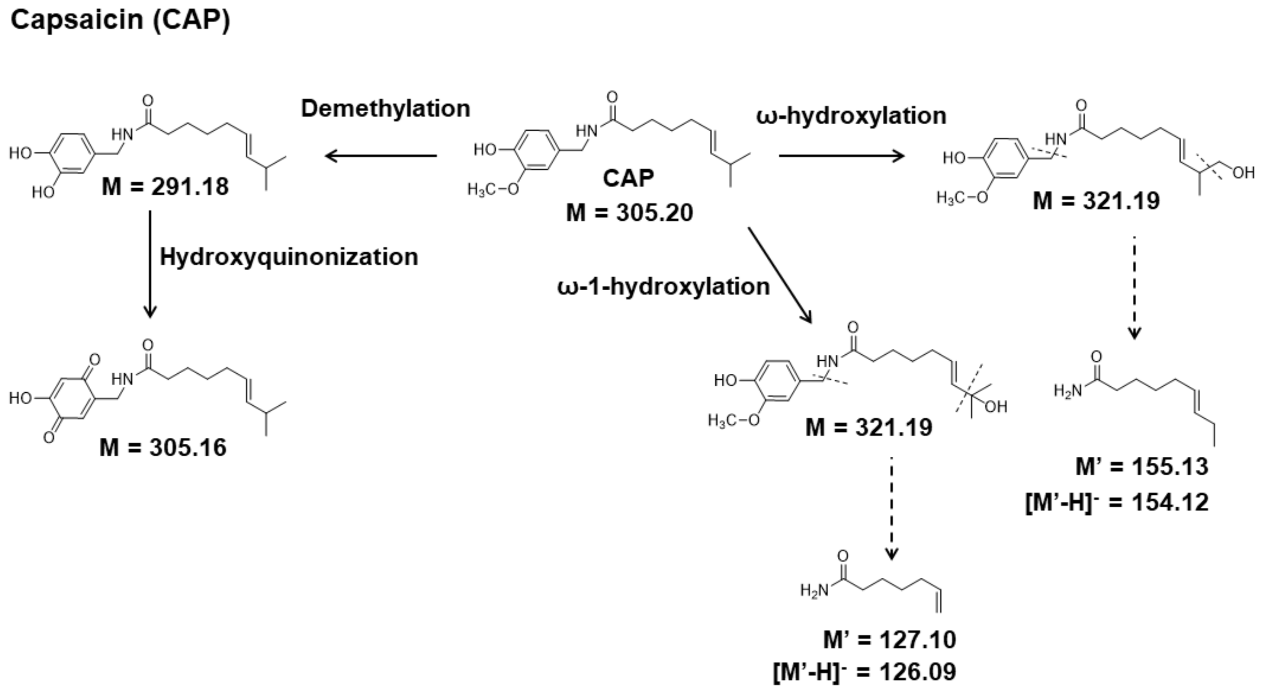

**Figure S19.** Fragmentation scheme of capsaicin oxygenation.

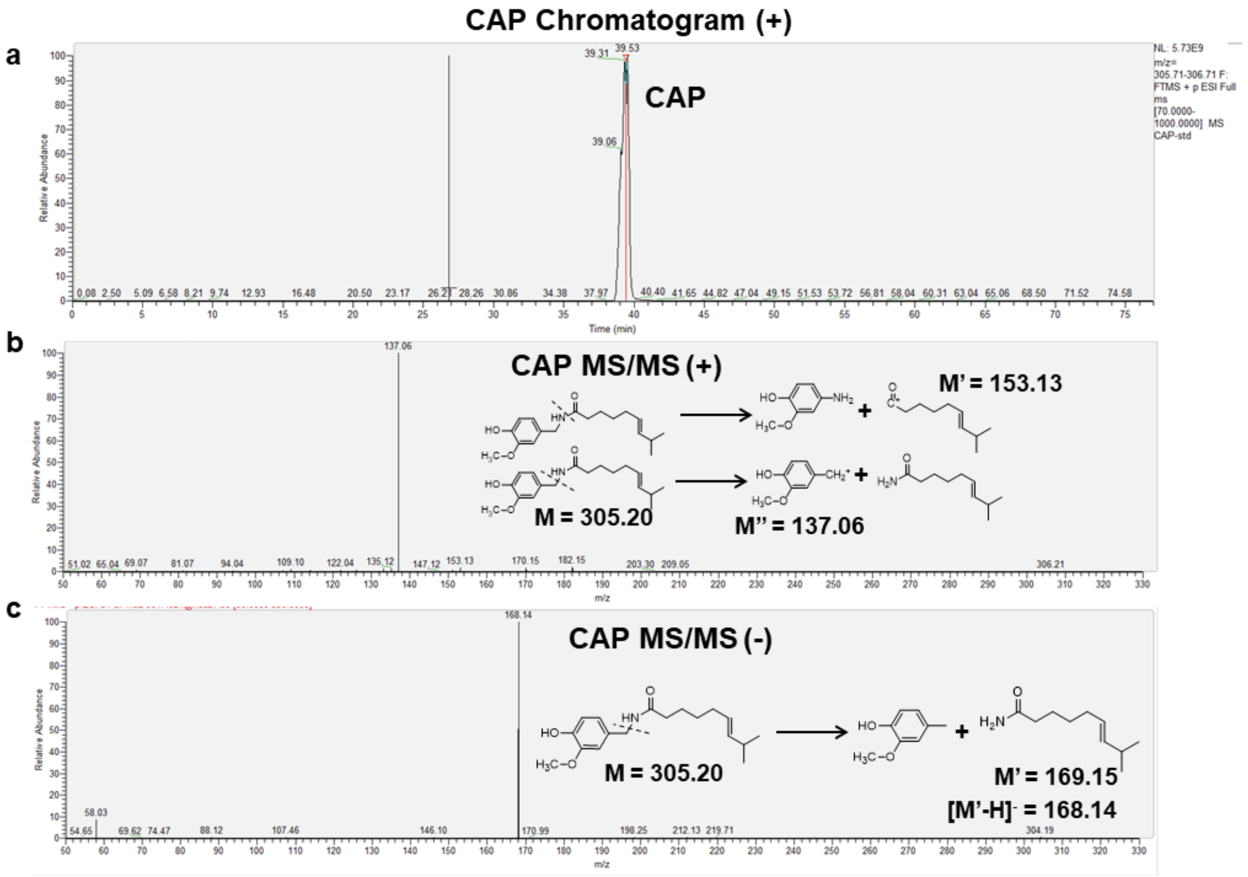

**Figure S20.** *LC-MS/MS analysis of capsaicin (CAP)*. **(a)** LC-MS/MS chromatogram in positive ion mode. Mass range is ± 5 ppm of the predicted product. **(b)** MS/MS spectrum of CAP in positive ion mode. **(c)** MS/MS spectrum of CAP in negative ion mode.

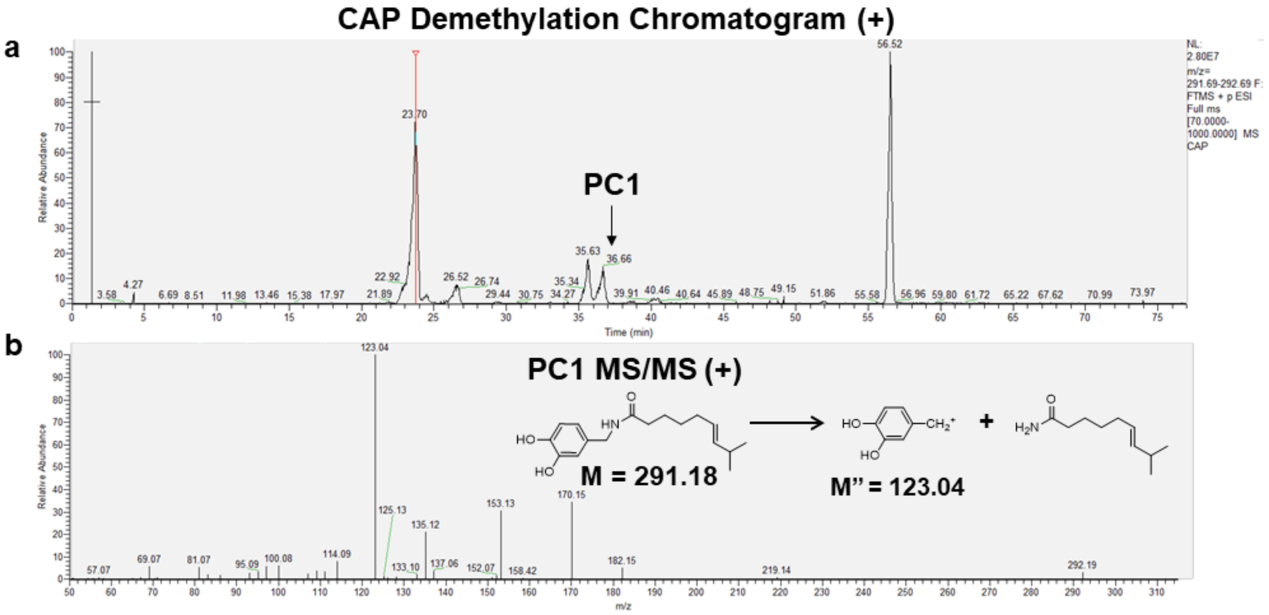

**Figure S21.** *Demethylation of CAP by CYP2J2*. **(a)** LC-MS/MS chromatogram in positive ion mode. Mass range is ± 5 ppm of the predicted product. **(b)** MS/MS spectrum of PC1.

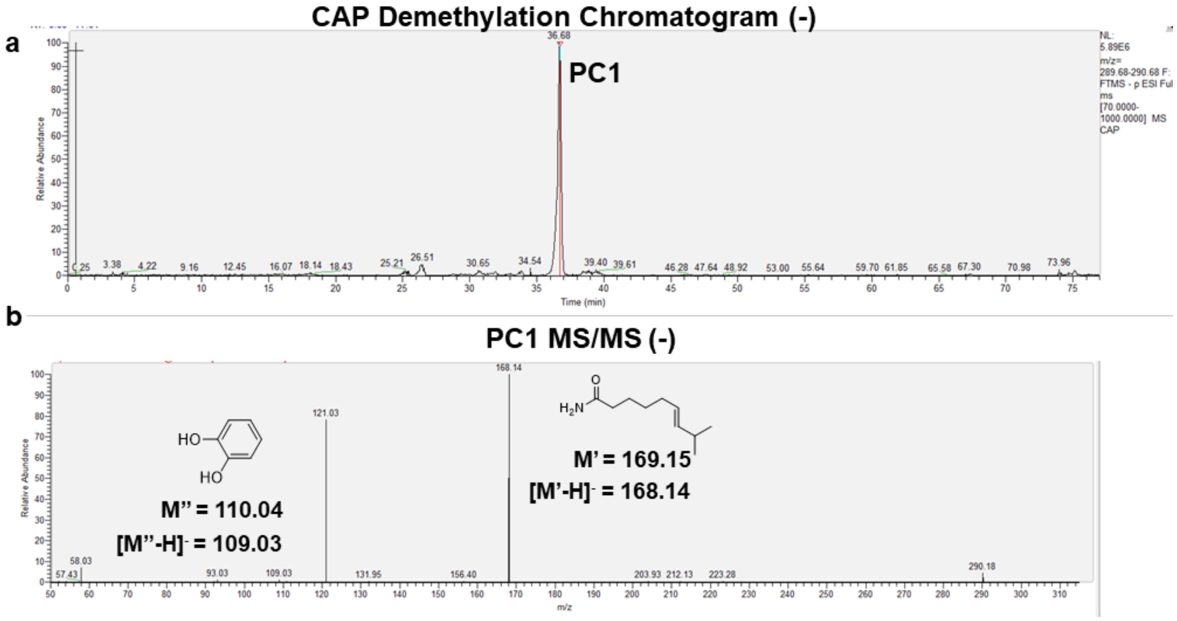

**Figure S22.** *Demethylation of CAP by CYP2J2*. **(a)** LC-MS/MS chromatogram in negative ion mode. Mass range is ± 5 ppm of the predicted product. **(b)** MS/MS spectrum of PC1.

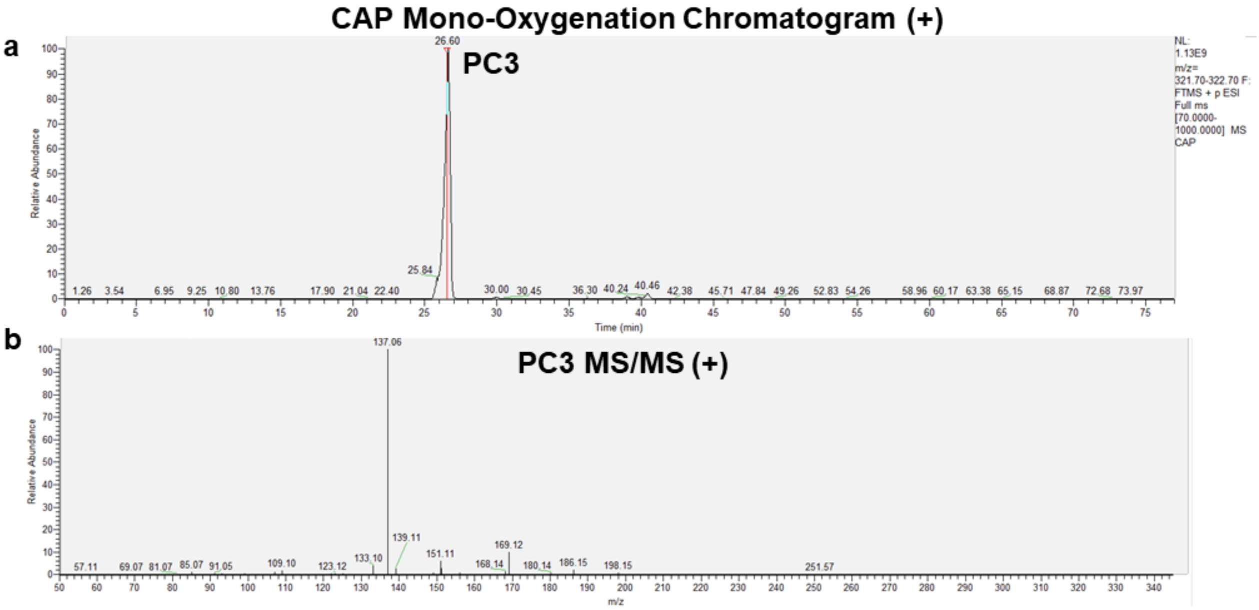

**Figure S23.** *Mono-oxygenation of CAP by CYP2J2*. **(a)** LC-MS/MS chromatogram in positive ion mode. Mass range is ± 5 ppm of the predicted product. **(b)** MS/MS spectrum of PC3.

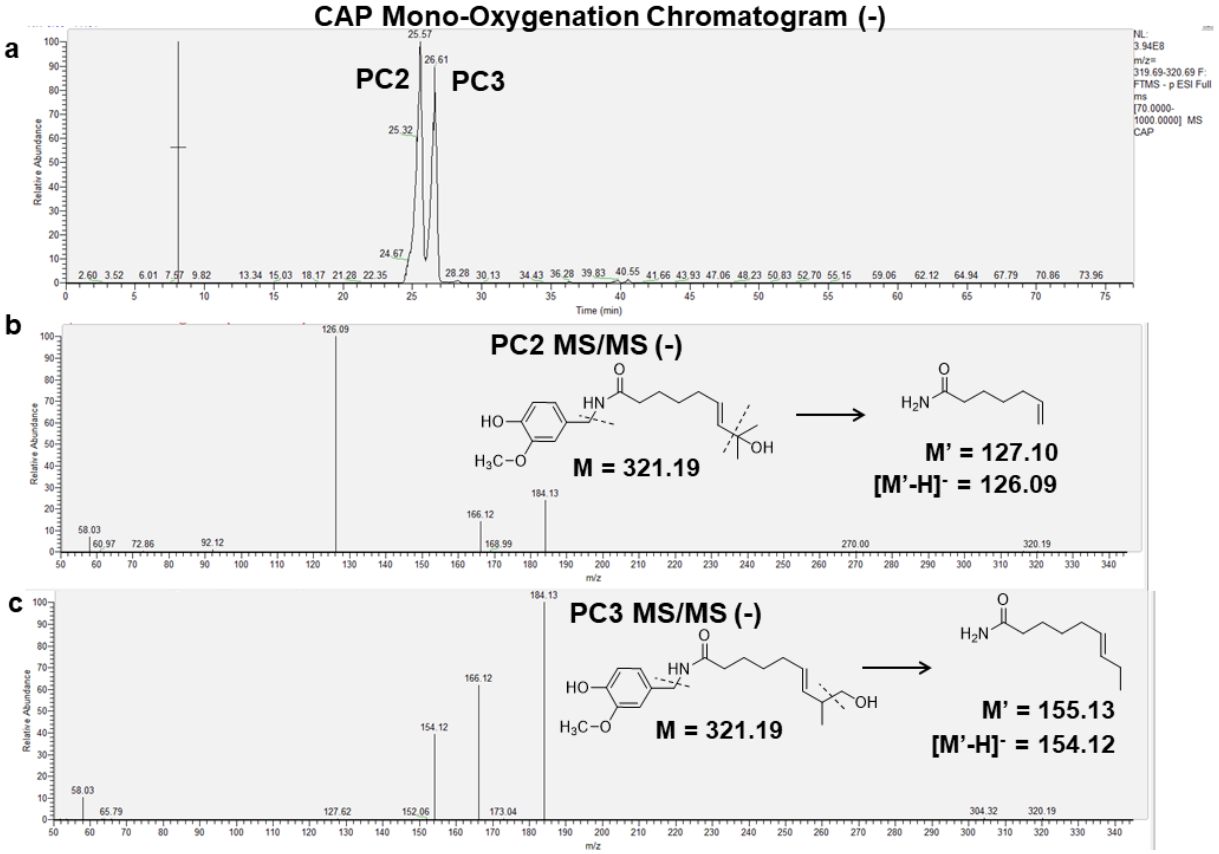

**Figure S24.** *Mono-oxygenation of CAP by CYP2J2*. **(a)** LC-MS/MS chromatogram in negative ion mode. Mass range is ± 5 ppm of the predicted product. **(b-c)** MS/MS spectra of indicated products.

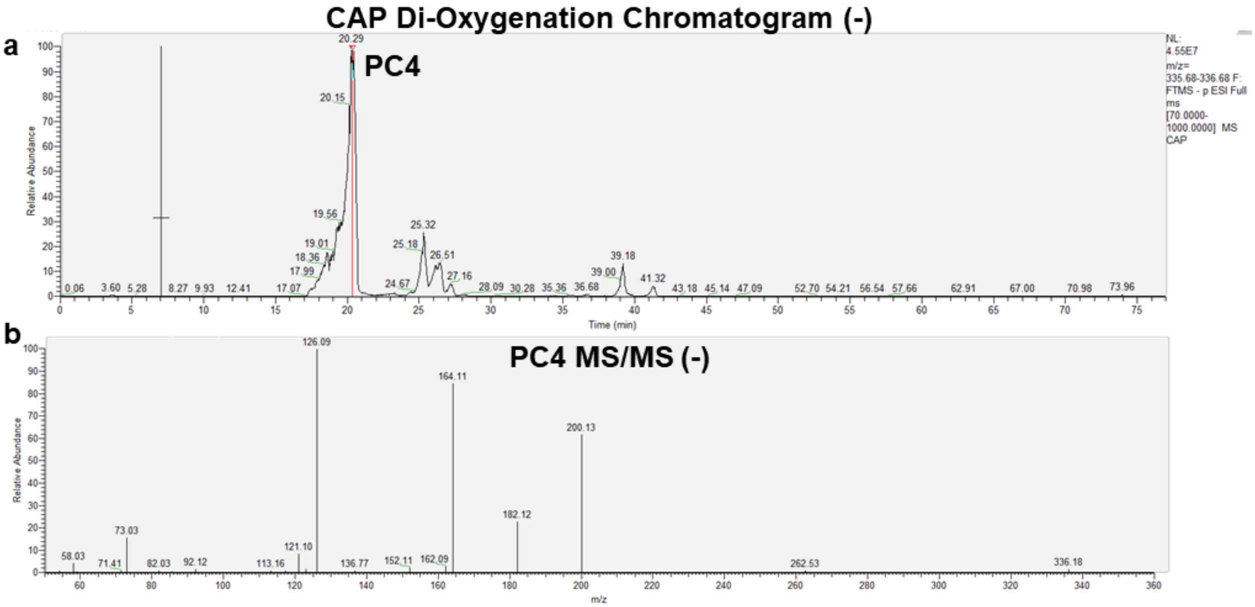

**Figure S25.** *Di-oxygenation of CAP by CYP2J2*. **(a)** LC-MS/MS chromatogram in negative ion mode. Mass range is ± 5 ppm of the predicted product. **(b)** MS/MS spectrum of PC4.

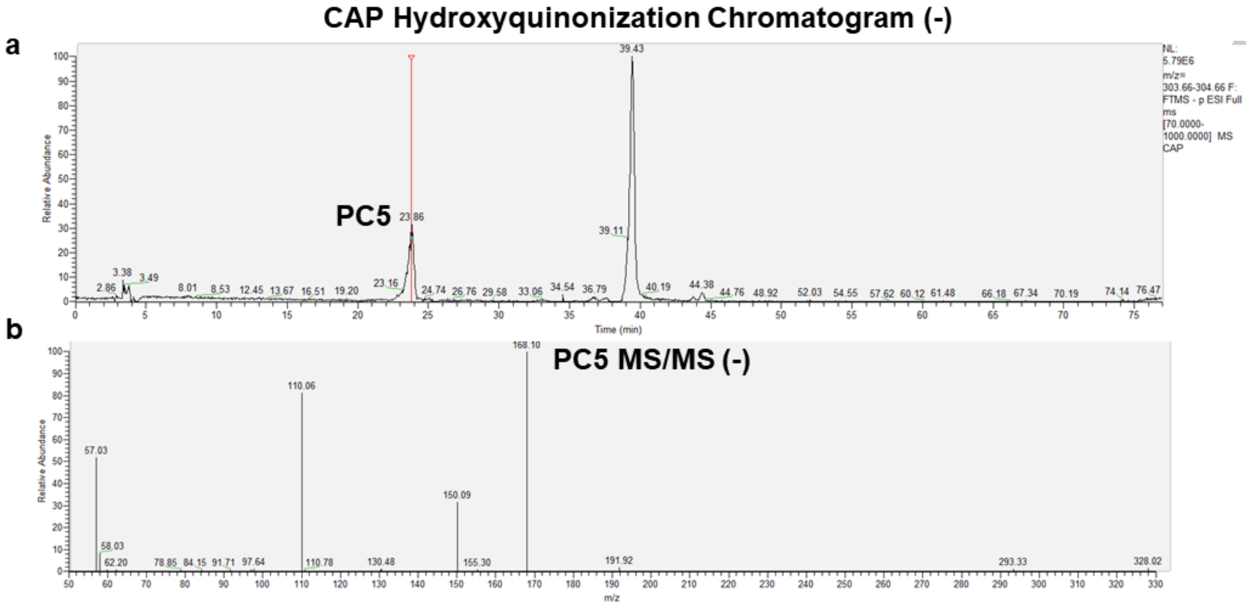

**Figure S26.** *Hydroxyquinonization of CAP by CYP2J2*. Hydroxyquinonization is determined by an oxidation of the demethylated CAP (PC1) indicated by the loss of two hydrogen atoms. **(a)** LC-MS/MS chromatogram in negative ion mode. Mass range is ± 5 ppm of the predicted product. **(b)** MS/MS spectrum of PC5.

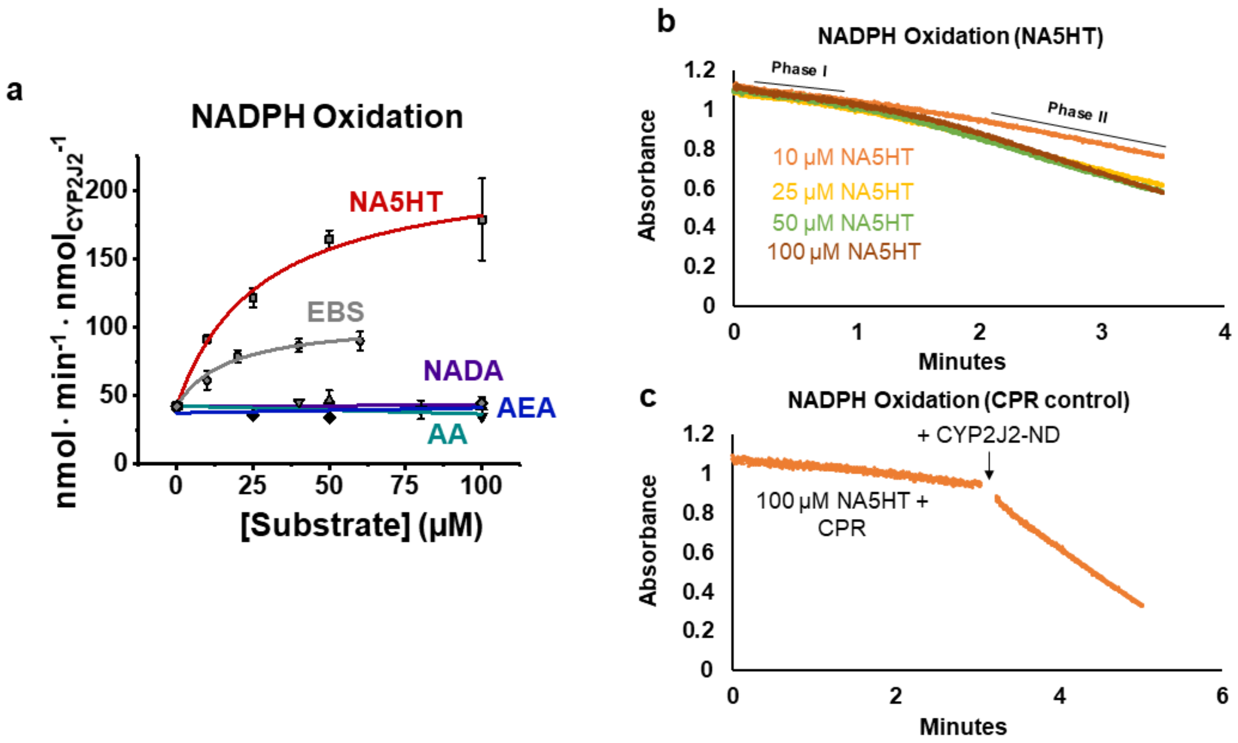

**Figure S27.** *Biphasic NADPH oxidation in the presence of NA5HT*. NADPH oxidation as CYP2J2-ND metabolizes NA5HT was determined by the absorbance of NADPH at 340 nm. Data represents the SEM of three independent experiments. Representative spectra are shown. **(a)** NADPH oxidation rates with the indicated substrates. NA5HT showed biphasic NADPH oxidation kinetics and the rates of Phase II are plotted. Ebastine (EBS) is monophasic. **(b)** Absorbance at 340 nm in the presence of the indicated amounts of NA5HT. Phase I rates are similar to those obtained with CPR and CYP2J2-ND alone. Phase II shows a decrease in NADPH as a function of NA5HT concentration. **(c)** Experiments were repeated with NA5HT and CPR only. Rate of NADPH oxidation is similar to CPR alone. Adding CYP2J2-ND results in a rate of NADPH oxidation similar to Phase II with 100 μM NA5HT as seen in **(b)**.

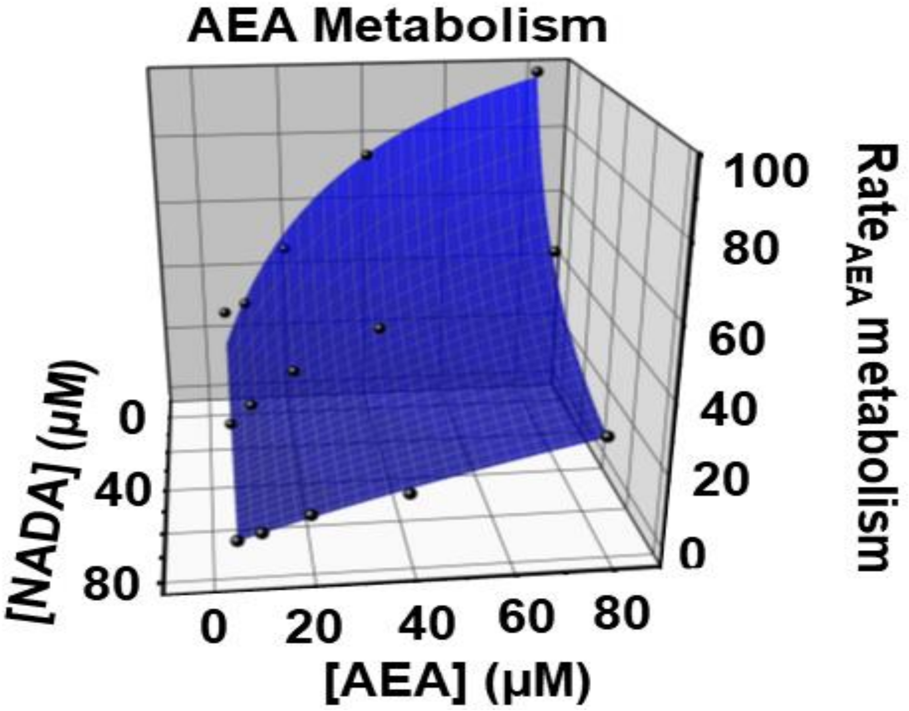

**Figure S28.** 3D global fit of AEA inhibition by NADA as shown in Figure 3f. Rates are in $pmol_{EET\text{-}EAs} \cdot min^{-1} \cdot nmol_{CYP2J2}^{-1}$.

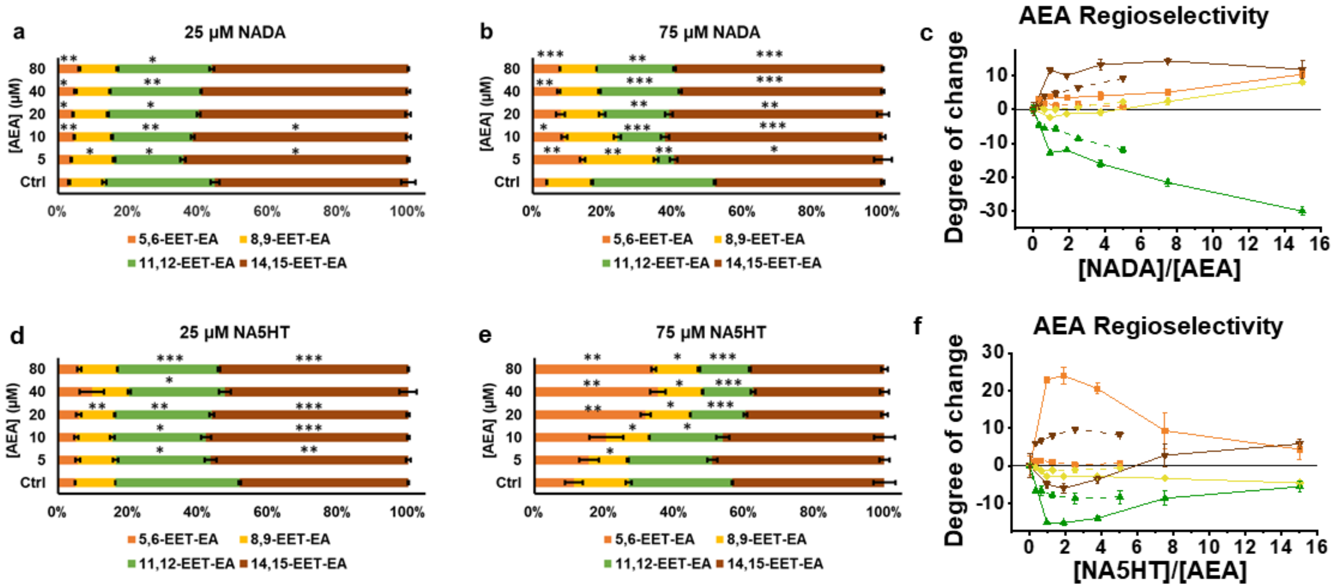

**Figure S29.** *Regioselectivity of AEA epoxidation in the presence of eVDs*. **(a-c)** Regioselectivity in the presence of NADA and **(d-f)** NA5HT at the indicated concentrations. Regioselectivity is shown as a percentage of the total EET-EAs. Control (Ctrl) refers to 40 µM AEA samples without eVDs, which was ran along with each experiment. *p < 0.05; **p < 0.01, ***p < 0.001. Data in **(c)** and **(f)** show the dependence of the regioselectivity change on the concentrations of the eVDs as a function of an increasing ratio of the eVD:AEA. Degree of change is defined as the change in percentage value compared to control. Data from the 25 µM datasets are shown as dashed lines and the data from the 75 µM datasets are shown as solid lines.

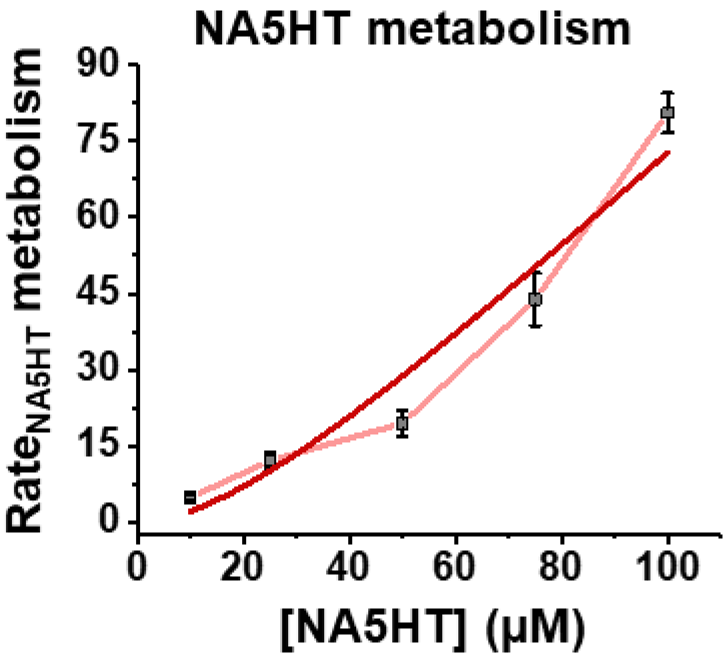

**Figure S30.** NA5HT metabolism data fitted to a two-site (Equation 4) binding equation as detailed in the Results section.

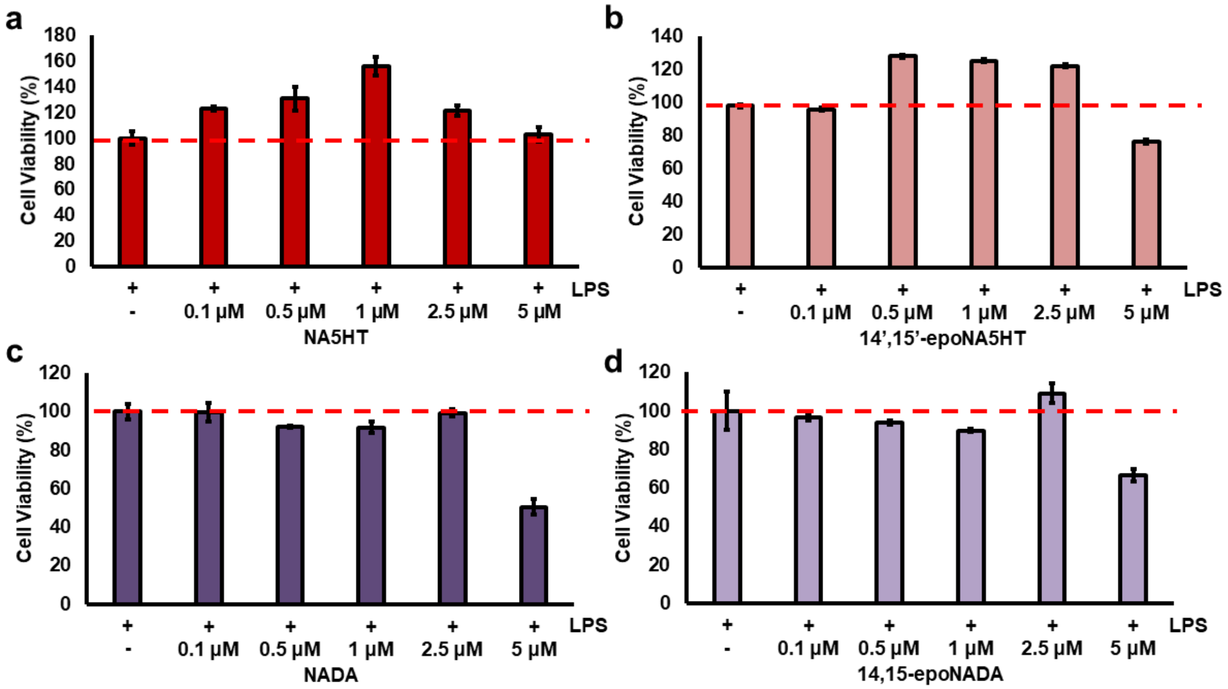

**Figure S31.** *MTT assay to determine cell viability.* BV2 microglial cells seeded at 200,000 cells/well in a 24-well plate were treated with 25 ng/mL LPS for 24 hours and an MTT cell proliferation assay was performed to determine cell viability. Viability was determined as a percent of the formazan absorbance at 570 nm compared to LPS treatment only (100%) in the presence of **(a)** NA5HT, **(b)** 14',15'-epoNA5HT, **(c)** NADA, and **(d)** 14',15'-epoNADA. Data represents the SEM of 3 independent experiments.

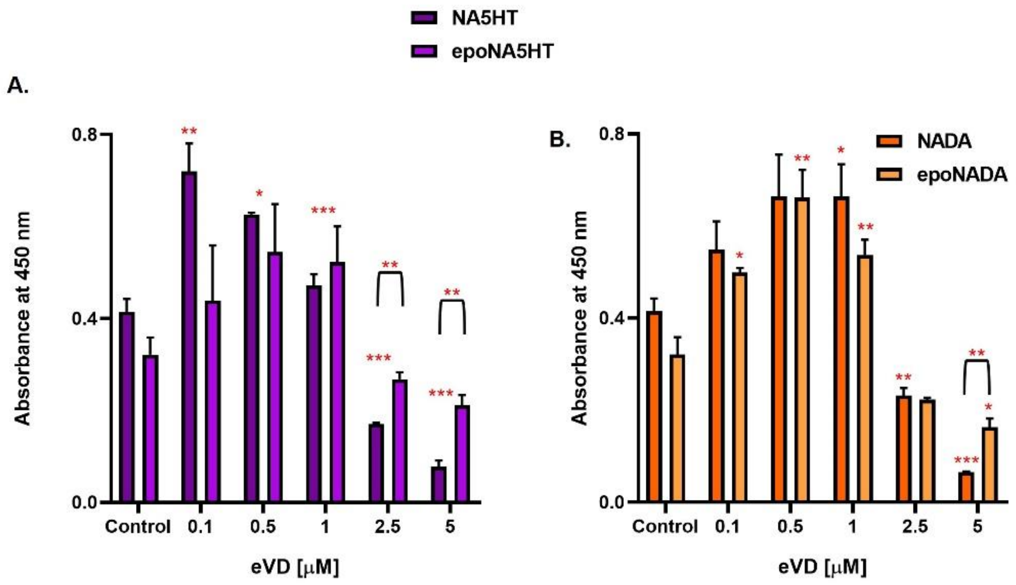

**Figure S32.** *Effects of eVDS and epoxy-eVDs on cell proliferation.* BV-2 microglial cells were pre-incubated with varying concentrations of **(A)** NA5HT and 14,15-epoNA5HT and **(B)** NADA and 14,15-epoNADA for 4 hours followed by LPS (25 ng/mL) stimulation for 24 hours. After 21 hours, BV-2 cell proliferation was measured using BrdU incorporation colorimetric ELISA assay. Values shown are the mean ± SEM of experiments performed in triplicate ($n = 3$). $*P < 0.05$, $**P < 0.01$, and $***P < 0.001$.

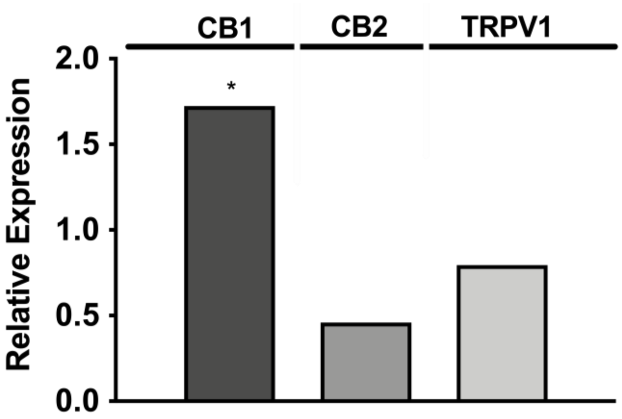

**Figure S33**. *Relative mRNA Expression of CB1, CB2, and TRPV1 in LPS-treated BV2 microglial cells.* BV2 microglial cells seeded at 200,000 cells/well in a 24-well plate were treated with 25 ng/mL LPS for 24 hours. RT-qPCR detected the presence of TRPV1, CB1, and CB2 in BV2 cells. Data is from 3-pooled wells and is performed in technical replicates. Data is reported as the mean relative expression. Statistical analysis was performed using a two-tailed t-test ($\alpha = 0.05$). P-values: 0.0332 (*), 0.0021(**), 0.0002 (***), and < 0.0001 (****).

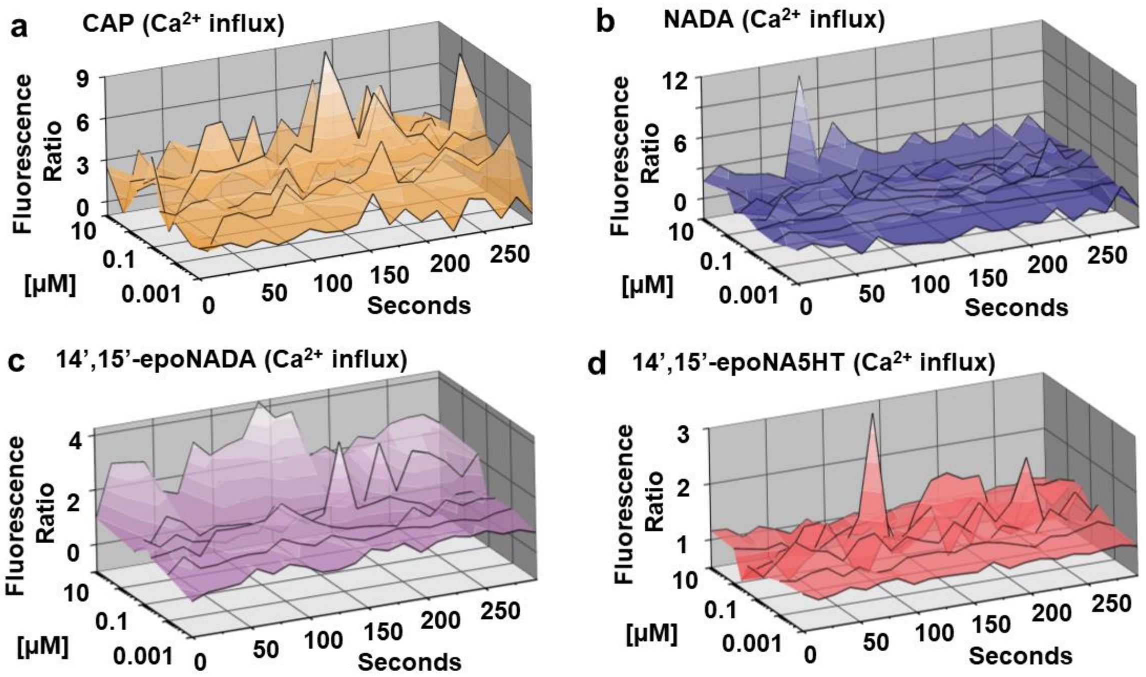

**Figure S34.** *TRPV1 activation by vanilloids.* Relative $Ca^{2+}$ influx upon TRPV1 activation was measured using a Fura-2 AM fluorescence assay. The fluorescence intensity at 510 nm from the $Ca^{2+}$-bound (excitation 340 nm) and $Ca^{2+}$-free (excitation 380 nm) dye was measured over time at varying concentrations of ligands and are reported as a ratio (Fluorescence Ratio). Representative plots of **(a)** Capsaicin (CAP), **(b)** NADA, **(c)** 14',15'-epoNADA, and **(d)** 14',15'-epoNA5HT from 4-6 separate experiments (2 sets of triplicate on separate days) are shown.

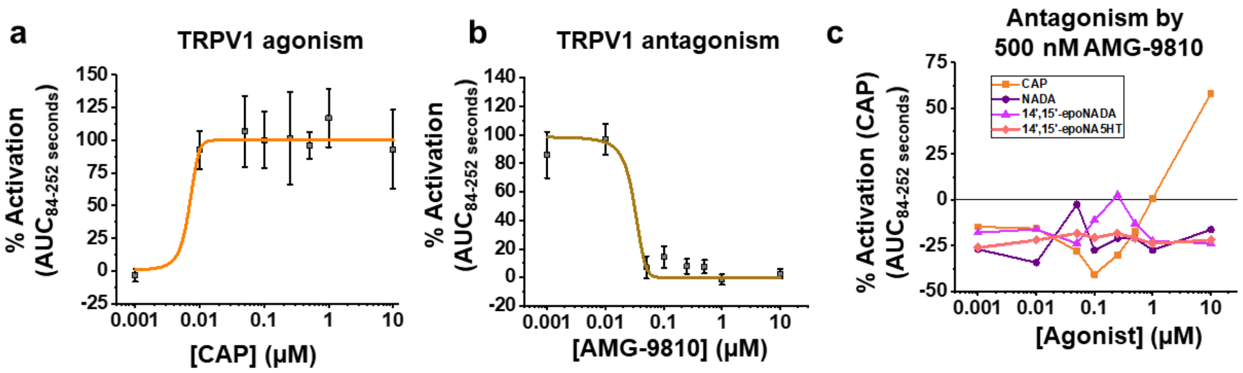

**Figure S35.** *TRPV1-mediated $Ca^{2+}$ influx agonized by CAP and antagonized by AMG-9810.* A Fura 2-AM fluorescence assay was used to determine TRPV1 binding in transfected HEK cells. 100% is defined as the $B_{max}$ of CAP binding. **(a)** CAP agonism data. **(b)** Antagonism of TRPV1 by AMG-9810. 250 nM of CAP was used to activate TRPV1. Data in **(a-b)** is shown as the SEM of 4-6 experiments (2 sets of triplicate performed on separate days). **(c)** HEK-TRPV1 cells were treated with 500 nM of reversible antagonist AMG-9810 30 min prior to stimulating with the indicated agonists. AMG-9810 prevented agonism of TRPV1 for all compounds. Data represents the SEM of 3 independent experiments.

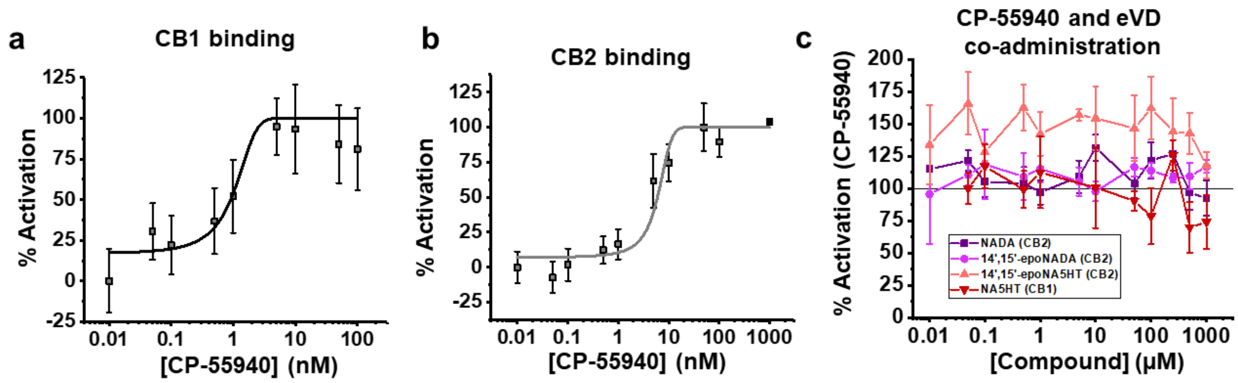

**Figure S36.** *PRESTO-Tango assays of CB1 and CB2 activation.* **(a)** Agonism of CB1 and **(b)** CB2 by CP-55940. Data respresents the SEM of 4-6 separate experiments (2 sets of triplicates from separate days). **(c)** Co-administration of eVDs and CP-55940 to determine eVD binding to CB1 and 2. CB1 and CB2 were stimulated with 50 nM of CP-55940 to determine antagonism by NADA (CB2), 14’,15’-epoNADA (CB2), 14’,15’-epoNA5HT (CB2), and NA5HT (CB1). 100% is defined as the $B_{max}$ of CP-55940 binding. Data represents the SEM of 3 independent experiments.

### Tables

**Table S1.** Primers for Quantitative Polymerase Chain Reaction

| Gene | Forward Primer (5' → 3') | Reverse Primer (5' → 3') | Reference |
|---|---|---|---|
| *Cyp2j9* | AACCGTTTCCCCAGTCAGTC | ATTGCACGCACTCTCTGTCA | 9 |
| *Cyp2j12* | ATGGAAGACAGCCAAGCACA | AGCCAGTGTGCTATCAGCTG | (designed in-house) |
| *Il-6* | CCAGAGATACAAAGAAATGATGG | ACTCCAGAAGACCAGAGGAAAT | 10 |
| *Il-10* | CATGGCCCAGAAATCAAGGA | GGAGAAATCGATGACAGCGC | 10 |
| *Il-1β* | AAATACCTGTGGCCTTGGGC | CTTGGGATCCACACTCTCCAG | 10 |
| *Tnf-α* | GCCACCACGCTCTTCTGCCT | GGCTGATGGTGTGGGTGAGG | 10 |
| *Cnr1* | TGAAGTCGATCTTAGACGGCC | GTGGTGATGGTACGGAAGGTA | 11 |
| *Cnr2* | TGAATGAGCAGACCGACA GG | AGAGATGTTTGCTGG GTGGC | 11 |
| *Trpv1* | TCGTCTACCTCGTGTTCTTGTTTG | CCAGATGTTCTTGCTCTCTTGTGC | 12 |
| *Gapdh* | TCACCACCATGGAGAAGGC | GCTAAGCAGTTGGTGGTGCA | 13 |

**Table S2.** LC-MS/MS metabolite identification

| Substrate | Product | Formulae | m/z (+)<br>m/z (-) | Retention Times (Min) | Metabolite ID |
|---|---|---|---|---|---|
| NADA | Parent | $C_{28}H_{41}NO_3$ | 440.3159<br>438.3014 | 55.4 | |
| | -O | $C_{28}H_{41}NO_4$ | 456.3108<br>454.2963 | 41.6; 42.6; 44.6; 45.6; 47.8; 49.0; 49.6; 52.6 | PD1-PD8 |
| | -$O_2$ | $C_{28}H_{41}NO_5$ | 472.3057<br>470.2912 | 48.8<br>53.0 | PD9 & PD10 |
| | -HQ | $C_{28}H_{39}NO_4$ | 454.2952<br>452.2806 | 46.3 | PD11 |
| NA5HT | Parent | $C_{30}H_{42}N_2O_2$ | 463.3319<br>461.3174 | 55.6 | |
| | -O | $C_{30}H_{42}N_2O_3$ | 479.3268<br>477.3122 | 42.4; 48.3; 52.4; 54.1 | PS1-PS4 |
| | -$O_2$ | $C_{30}H_{42}N_2O_4$ | 495.3217<br>493.3072 | 50.5 | PS5 |
| | -Q | $C_{30}H_{40}N_2O_3$ | 477.3112<br>475.2969 | 54.7<br>56.8 | PS6 & PS7 |
| CAP | Parent | $C_{18}H_{27}NO_3$ | 306.2064<br>304.1918 | 39.5 | |
| | -*O*-de-$CH_3$ | $C_{17}H_{25}NO_3$ | 292.1907<br>290.1762 | 36.7 | PC1 |
| | -O | $C_{18}H_{27}NO_4$ | 322.2013<br>320.1867 | 25.6<br>26.6 | PC2 & PC3 |
| | -$O_2$ | $C_{18}H_{27}NO_5$ | 338.1962<br>336.1816 | 20.3 | PC4 |
| | -HQ | $C_{17}H_{23}NO_4$ | 306.1700<br>304.1554 | 23.8 | PC5 |

**Table S3.** Binding interaction energies for AEA and NADA in Configuration 1.

| **AEA** | | | |
|---|---|---|---|
| | **Residue** | **Interaction energy (kJ/mol)** | **Error (±kJ/mol)** |
| | ARG321 | -8.92481 | 3.47937 |
| | ARG495 | -4.72929 | 1.37637 |
| | ARG484 | -3.76565 | 5.08937 |
| | ASN185 | -3.6608 | 0.91729 |
| | LEU215 | -3.13021 | 0.90826 |
| | HIS494 | -3.10337 | 2.63799 |
| | HIS181 | -2.99208 | 1.25837 |
| | SER493 | -2.98903 | 1.15341 |
| | PRO491 | -2.91059 | 2.22862 |
| | TRP322 | -2.7051 | 1.87051 |
| | VAL492 | -2.34359 | 0.69187 |
| | LEU212 | -2.29354 | 0.802 |
| | THR313 | -2.29139 | 0.656 |
| | LEU481 | -2.25677 | 2.00732 |
| | LEU496 | -1.88488 | 0.98401 |
| | SER480 | -1.76965 | 1.92706 |
| | THR219 | -1.65714 | 0.72366 |
| | ASP216 | -1.53383 | 0.59384 |
| | PHE483 | -1.15568 | 1.16863 |
| **NADA** | | | |
| | **Residue** | **Interaction energy (kJ/mol)** | **Error (±kJ/mol)** |
| | ILE489 | -3.25625 | 0.96471 |
| | GLU222 | -2.81904 | 7.20851 |
| | THR114 | -2.64145 | 1.26045 |
| | ILE127 | -2.4887 | 0.672 |
| | THR488 | -2.42929 | 1.62007 |
| | PHE310 | -2.33006 | 0.68078 |
| | ASN231 | -2.13378 | 1.40423 |
| | ILE487 | -2.12379 | 1.17521 |
| | ARG117 | -2.0808 | 1.03731 |
| | MET400 | -2.0443 | 0.97651 |
| | ILE86 | -1.86892 | 0.99568 |
| | VAL380 | -1.86384 | 0.53806 |
| | ALA311 | -1.75345 | 0.60574 |
| | MET128 | -1.71349 | 0.69304 |
| | PRO381 | -1.52613 | 0.65021 |

| | | | |
|---|---|---|---|
| | LEU83 | -1.51171 | 0.79452 |
| | GLN228 | -1.16917 | 1.08287 |
| | GLU314 | -1.15822 | 0.47136 |
| | THR315 | -0.98946 | 0.40637 |

**Table S4.** Binding interaction energies for AEA and NADA in Configuration 2.

| | | | |
|---|---|---|---|
| **AEA** | | | |
| | **Residue** | **Interaction energy (kJ/mol)** | **Error (±kJ/mol)** |
| | ILE487 | -5.89876 | 1.21444 |
| | GLN228 | -5.83744 | 2.55033 |
| | GLY486 | -4.4337 | 0.85385 |
| | ILE489 | -3.73012 | 1.80267 |
| | THR488 | -2.86717 | 1.23994 |
| | LEU83 | -2.66672 | 1.22142 |
| | PRO381 | -2.01531 | 0.5883 |
| | MET400 | -1.92187 | 0.59975 |
| | ARG111 | -1.86332 | 0.65592 |
| | PHE61 | -1.76106 | 0.70389 |
| | PRO112 | -1.56523 | 0.63967 |
| | THR114 | -1.38301 | 0.59581 |
| | VAL380 | -1.31937 | 0.42989 |
| | LEU81 | -1.29664 | 0.75077 |
| | ILE86 | -1.23497 | 0.69801 |
| | ASN231 | -1.23156 | 0.70985 |
| | GLU82 | -1.21236 | 0.75533 |
| | ILE127 | -0.93357 | 0.4817 |
| | MET485 | -0.90994 | 0.50495 |
| **NADA** | | | |
| | **Residue** | **Interaction energy (kJ/mol)** | **Error (±kJ/mol)** |
| | ASP307 | -24.8669 | 9.39773 |
| | PHE310 | -5.45818 | 1.08501 |
| | SER490 | -4.84377 | 2.57133 |
| | ILE489 | -3.81991 | 1.27441 |
| | ALA311 | -2.55907 | 0.64457 |
| | GLU222 | -2.43978 | 6.04616 |
| | THR315 | -2.4274 | 0.79848 |
| | GLU314 | -1.99078 | 0.60895 |
| | ILE127 | -1.74117 | 0.56921 |
| | VAL380 | -1.56889 | 0.67457 |
| | ILE376 | -1.45199 | 0.64053 |
| | ILE375 | -1.40588 | 0.55081 |
| | THR318 | -1.24705 | 0.4895 |
| | TRP251 | -1.15856 | 0.49279 |
| | PHE121 | -1.06322 | 0.47599 |

| | | | |
|---|---|---|---|
| | THR219 | -0.99602 | 0.86035 |
| | PRO491 | -0.87034 | 0.55172 |
| | LEU126 | -0.76823 | 0.67118 |
| | ARG111 | -0.72046 | 0.51937 |

**Table S5.** Binding interaction energies for AEA and NA5HT in Configuration 1.

| | | | |
|---|---|---|---|
| **AEA** | | | |
| | **Residue** | **Interaction energy (kJ/mol)** | **Error (±kJ/mol)** |
| | ARG321 | -21.757 | 3.21232 |
| | TRP322 | -4.69984 | 1.39869 |
| | PRO491 | -4.12342 | 1.47182 |
| | ARG495 | -3.77759 | 1.6088 |
| | VAL492 | -3.66875 | 0.78594 |
| | SER493 | -2.94951 | 0.78101 |
| | ASN185 | -2.82361 | 0.97184 |
| | THR318 | -2.74648 | 0.88867 |
| | LEU215 | -2.29793 | 0.59603 |
| | LEU496 | -2.1508 | 0.48536 |
| | THR219 | -1.81981 | 0.54936 |
| | ASP216 | -1.45737 | 0.6174 |
| | THR313 | -1.44611 | 0.63856 |
| | HSD181 | -1.23107 | 0.92312 |
| | LEU212 | -1.22474 | 0.5249 |
| | SER317 | -1.13683 | 0.42019 |
| | SER480 | -1.12392 | 0.50546 |
| | SER490 | -1.05437 | 0.5769 |
| | LEU325 | -0.94713 | 0.42667 |
| **NA5HT** | | | |
| | **Residue** | **Interaction energy (kJ/mol)** | **Error (±kJ/mol)** |
| | GLU222 | -5.78558 | 3.11688 |
| | THR488 | -3.52051 | 2.94825 |
| | ILE489 | -3.40178 | 1.07731 |
| | VAL380 | -3.0252 | 0.61197 |
| | GLN228 | -3.00795 | 2.10628 |
| | ARG117 | -2.86239 | 1.37033 |
| | THR114 | -2.8041 | 0.76497 |
| | ARG111 | -2.58118 | 0.899 |
| | ILE127 | -2.16936 | 0.67201 |
| | ASN231 | -1.96704 | 0.91609 |
| | PRO112 | -1.8457 | 0.72025 |
| | PHE310 | -1.78344 | 0.73298 |
| | ALA311 | -1.49405 | 0.50818 |
| | PRO381 | -1.43143 | 0.66267 |
| | SER490 | -1.38003 | 0.99802 |

| | | | |
|---|---|---|---|
| | MET400 | -1.26626 | 0.51045 |
| | ILE487 | -1.22545 | 1.01001 |
| | THR315 | -1.11141 | 0.64492 |
| | GLY486 | -1.00512 | 0.58716 |

**Table S6.** Binding interaction energies for AEA and NA5HT in Configuration 2.

| | | | |
|---|---|---|---|
| **AEA** | | | |
| | **Residue** | **Interaction energy (kJ/mol)** | **Error (±kJ/mol)** |
| | GLN228 | -6.5189 | 1.61495 |
| | ILE487 | -6.4654 | 1.07545 |
| | GLY486 | -5.05945 | 0.69118 |
| | ASN231 | -3.30921 | 1.13293 |
| | THR488 | -2.75129 | 0.70766 |
| | MET400 | -2.60969 | 0.64574 |
| | LEU83 | -2.51603 | 0.82828 |
| | ILE86 | -2.41382 | 0.70615 |
| | ILE489 | -2.06282 | 0.60377 |
| | PHE61 | -1.61387 | 0.53751 |
| | GLU222 | -1.59248 | 1.60739 |
| | LEU81 | -1.54611 | 0.58844 |
| | SER490 | -1.37099 | 0.95213 |
| | GLU82 | -1.32141 | 0.53585 |
| | ALA88 | -1.07485 | 0.34869 |
| | MET485 | -1.04278 | 0.39769 |
| | ALA223 | -1.02855 | 0.51492 |
| | THR114 | -1.0283 | 0.63714 |
| | VAL232 | -0.97153 | 0.50219 |
| **NA5HT** | | | |
| | **Residue** | **Interaction energy (kJ/mol)** | **Error (±kJ/mol)** |
| | ARG117 | -4.62722 | 1.01273 |
| | PHE310 | -4.25033 | 0.78785 |
| | ILE489 | -3.79065 | 0.7503 |
| | PRO112 | -3.53451 | 2.60917 |
| | ILE127 | -3.16155 | 0.66233 |
| | VAL380 | -3.0195 | 0.73603 |
| | ARG111 | -2.8175 | 0.86368 |
| | ALA311 | -2.54837 | 0.69605 |
| | GLU314 | -2.41186 | 0.5614 |
| | THR315 | -2.06731 | 0.79649 |
| | THR114 | -1.5682 | 0.61104 |
| | ILE376 | -1.48917 | 0.57733 |
| | THR318 | -1.26557 | 0.42192 |
| | ILE375 | -1.19388 | 0.4528 |
| | GLU222 | -0.95049 | 0.97809 |

| | | | |
|---|---|---|---|
| | VAL113 | -0.87465 | 0.51344 |
| | SER490 | -0.83479 | 0.43952 |
| | THR488 | -0.62754 | 0.11655 |
| | PRO491 | -0.60725 | 0.5804 |

#### Supplemental Movies

**Movie S1.** *Molecular dynamics (MD) simulations of NADA and AEA in Configuration 1.* MD simulations were performed for 50 ns as described in the Methods section. In this configuration, AEA is bound in the PUFA binding pocket and NADA in the substrate access channel. Green spheres represent the 14’ and 15’ positions of NADA.

**Movie S2.** *Molecular dynamics (MD) simulations of NADA and AEA in Configuration 2.* MD simulations were performed for 50 ns as described in the Methods section. In this configuration, AEA is bound at the entrance of the substrate access channel and stabilizes the binding of NADA near the heme. Green spheres represent the 14’ and 15’ positions of NADA.

**Movie S3.** *Molecular dynamics (MD) simulations of NA5HT and AEA in Configuration 1.* MD simulations were performed for 50 ns as described in the Methods section. In this configuration, AEA is bound in the PUFA binding pocket and NA5HT in the substrate access channel. Green spheres represent the 14’ and 15’ positions of NA5HT.

**Movie S4.** *Molecular dynamics (MD) simulations of NA5HT and AEA in Configuration 2.* MD simulations were performed for 50 ns as described in the Methods section. In this configuration, AEA is bound at the entrance of the substrate access channel and stabilizes the binding of NA5HT near the heme. Green spheres represent the 14’ and 15’ positions of NA5HT.

### Supplementary References

1 McDougle, D. R., Palaria, A., Magnetta, E., Meling, D. D. & Das, A. Functional studies of N-terminally modified CYP2J2 epoxygenase in model lipid bilayers. *Protein Sci* **22**, 964-979, doi:10.1002/pro.2280 (2013).

2 Schmittgen, T. D. & Livak, K. J. Analyzing real-time PCR data by the comparative C(T) method. *Nature protocols* **3**, 1101-1108 (2008).

3 McDougle, D. R. *et al.* Anti-inflammatory omega-3 endocannabinoid epoxides. *Proc Natl Acad Sci U S A* **114**, E6034-E6043, doi:10.1073/pnas.1610325114 (2017).

4 Zelasko, S., Palaria, A. & Das, A. Optimizations to achieve high-level expression of cytochrome P450 proteins using Escherichia coli expression systems. *Protein Expres Purif* **92**, 77-87, doi:DOI 10.1016/j.pep.2013.07.017 (2013).

5 Arnold, W. R., Baylon, J. L., Tajkhorshid, E. & Das, A. Arachidonic Acid Metabolism by Human Cardiovascular CYP2J2 Is Modulated by Doxorubicin. *Biochemistry* **56**, 6700-6712, doi:10.1021/acs.biochem.7b01025 (2017).

6 Arnold, W. R., Weigle, A. T. & Das, A. Cross-talk of cannabinoid and endocannabinoid metabolism is mediated via human cardiac CYP2J2. *Journal of inorganic biochemistry* **184**, 88-99, doi:10.1016/j.jinorgbio.2018.03.016 (2018).

7 Arnold, W. R., Baylon, J. L., Tajkhorshid, E. & Das, A. Asymmetric Binding and Metabolism of Polyunsaturated Fatty Acids (PUFAs) by CYP2J2 Epoxygenase. *Biochemistry* **55**, 6969-6980, doi:10.1021/acs.biochem.6b01037 (2016).

8 Capdevila, J. H. *et al.* The highly stereoselective oxidation of polyunsaturated fatty acids by cytochrome P450BM-3. *J Biol Chem* **271**, 22663-22671 (1996).

9 Graves, J. P. *et al.* Quantitative Polymerase Chain Reaction Analysis of the Mouse Cyp2j Subfamily: Tissue Distribution and Regulation. *Drug Metab Dispos* **43**, 1169-1180, doi:10.1124/dmd.115.064139 (2015).

10 Oh, D. Y. *et al.* GPR120 Is an Omega-3 Fatty Acid Receptor Mediating Potent Anti-inflammatory and Insulin-Sensitizing Effects. *Cell* **142**, 687-698, doi:10.1016/j.cell.2010.07.041 (2010).

11 Zhang, M. *et al.* Modulation of the balance between cannabinoid CB1 and CB2 receptor activation during cerebral ischemic/reperfusion injury. *Neuroscience* **152**, 753-760, doi:10.1016/j.neuroscience.2008.01.022 (2008).

12 Phan, T. X., Ton, H. T., Chen, Y., Basha, M. E. & Ahern, G. P. Sex-dependent expression of TRPV1 in bladder arterioles. *Am J Physiol-Renal* **311**, F1063-F1073, doi:10.1152/ajprenal.00234.2016 (2016).

13 Yen, C. H. *et al.* Characterization of a new murine cell line of sarcomatoid hepatocellular carcinoma and its application for biomarker/therapy development. *Sci Rep* **7**, 3052, doi:10.1038/s41598-017-03164-3 (2017).
